# Changes at V2 apex of HIV-1 Clade C trimer enhance elicitation of autologous neutralizing and broad V1V2-scaffold antibodies

**DOI:** 10.1101/2021.06.15.447100

**Authors:** Anusmita Sahoo, Edgar A. Hodge, Celia LaBranche, Tiffany Turner Styles, Xiaoying Shen, Narayanaiah Cheedarla, Ayalnesh Shiferaw, Gabriel Ozorowski, Wen-Hsin Lee, Andrew B. Ward, Georgia D. Tomaras, David C. Montefiori, Darrell J. Irvine, Kelly K. Lee, Rama Rao Amara

## Abstract

HIV-1 clade C envelope immunogens that elicit both neutralizing and non-neutralizing V1V2-scaffold specific antibodies (protective correlates from RV144 human trial) are urgently needed due to the prevalence of this clade in the most impacted regions worldwide. To achieve this, we introduced structure-guided changes followed by consensus-C sequence-guided optimizations at the V2-region to generate UFO-v2-RQH^173^ trimer. This improved the abundance of native-like trimers and carried an intrinsic dynamic V2-loop. Following immunization of rabbits, the wild-type protein failed to elicit any autologous neutralizing antibodies but UFO-v2-RQH^173^ elicited both autologous neutralizing and broad V1V2-scaffold antibodies. The variant with 173Y modification in V2-region, most prevalent among HIV-1 sequences, showed decreased ability in displaying native-like V1V2 epitope with time *in-vitro* and elicited antibodies with lower neutralizing and higher V1V2-scaffold activities. Our results identify a clade C C.1086-UFO-v2-RQH^173^ trimer capable of eliciting improved neutralizing and V1V2-scaffold antibodies, and reveal the importance of V2-region in tuning this.

## Introduction

Generation of broadly neutralizing antibodies (bnAbs) during the course of HIV-1 infection occurs in 10-25% of the population within 3-4 years after the infection (Binley et al., 2008; Doria-Rose et al., 2009; Hraber et al., 2014; Landais et al., 2016; Li et al., 2009). Robust induction of a similar immune response by vaccination has been a major challenge. Strategies to stabilize the envelope protein in closed trimeric conformation have advanced efforts to induce neutralizing antibodies. The development of SOSIP trimeric design (Sanders et al., 2013), followed by cleavage independent Native Flexibly Linked (NFL) (Sharma et al., 2015), Un-cleaved preFusion Optimized (UFO) (He et al., 2018; Kong et al., 2016) and “germline” bnAb precursor-targeting trimers (reviewed by McGuire, 2019) opened an era of well-ordered, native-like trimers (Aldon et al., 2018; Brouwer et al., 2019; Guenaga et al., 2015b; Guenaga et al., 2017; He et al., 2018; Kulp et al., 2017; Sliepen et al., 2015). Immunization with these different trimeric forms and their variants bearing further stabilizing substitutions as either soluble protein or multivalent nanoparticle display in either homologous or heterologous prime-boost vaccine regimens successfully elicited tier2 homologous envelope specific neutralizing Abs (de Taeye et al., 2015; Escolano et al., 2016; Huang et al., 2020; Klasse et al., 2016; Pauthner et al., 2017; Sanders et al., 2015; Voss et al., 2017). The responses, however, generally had poor neutralization breadth and were primarily directed towards epitopes specific to the immunizing envelope (Arunachalam et al., 2020; Bale et al., 2018; Dubrovskaya et al., 2017; He et al., 2018; Klasse et al., 2018; Pauthner et al., 2017; Sanders and Moore, 2017; Sanders et al., 2015; Torrents de la Pena et al., 2017). However, studies (Bricault et al., 2019; Escolano et al., 2016) including work by Xu et al., 2018, to generate broad neutralization activity towards the fusion-peptide, and Dubrovskaya et al., 2019 to induce cross-neutralizing responses towards CD4bs and gp120-gp41 interface, mark significant steps towards the induction of cross-reactive tier2 neutralizing activities by vaccination.

Antibodies with neutralizing properties can also display Fc-mediated anti-viral effector functions such as ADCC (antibody-dependent cellular cytotoxicity), ADCVI (antibody-dependent cell-mediated viral inhibition), ADCP (antibody-dependent cell-mediated phagocytosis) that help in clearing virus infected cells (Berendam et al., 2021). In addition to the neutralizing Abs, non-neutralizing Abs (non-nAbs) have been reported to contribute significantly towards protection in the RV144 human efficacy trial (Rerks-Ngarm et al., 2009). In the RV144 trial, individuals that generated a strong antibody response to HIV-1 V1V2 loops scaffolded on MuLV gp70 protein (V1V2-scaffold) and against a linear epitope in the V2 loop from residues 166-178 (V2 hotspot region, V2 HS (Tassaneetrithep et al., 2014)) proximal to the α_4_β_7_ binding site showed decreased risk of acquisition of HIV infection (Excler et al., 2014; Haynes et al., 2012; Jones et al., 2019; Robb et al., 2012; Zolla-Pazner et al., 2014; Zolla-Pazner et al., 2013). The protective responses associated with this class of Abs have been linked to (a) Fc-mediated anti-viral effector functions, and (b) their ability to occlude interaction between the host integrin α_4_β_7_ and HIV-1 envelope (known to promote viral pathogenesis (Guzzo et al., 2017; Plotnik et al., 2017)) in-order to control and clear the virus (Gorny et al., 2012; Perez et al., 2017; Yates et al., 2014). Hence, the ability to generate broad V1V2 reactivity has been an important parameter in the immunogen design portfolios.

The design of an immunogen with the potential to elicit both tier2 neutralizing and broad V1V2-scaffold specific binding antibodies has not been explored. A closed stabilized trimer designed to generate neutralizing Abs is less likely to expose the conformation presented by the V1V2-scaffolds which is more flexible and lacks the quaternary contacts, making it difficult to design an immunogen with a fine balance between the two kinds of responses. Although studies have reported the significance of V2 region in regulating the closed-state of the envelope and neutralization sensitivity (Cimbro et al., 2014; Guzzo et al., 2018), understanding how these changes maintain native apex integrity with time without impacting trimer integrity is unexplored. How do such changes affect the balance between inducing a neutralizing versus a broad V1V2-scaffold specific response is not known. We attempted to address these by engineering a stabilized clade C trimeric immunogen. Clade C immunogens are critically needed since a large proportion (∼48%) of global HIV-1 burden results from infection by viruses within this clade (Geretti et al., 2009). We focused our efforts on C.1086 envelope since a gp120 version of this protein is used in human phase 2b/3 trials by the HIV Vaccine Trials Network (HVTN) in HVTN702. There are only a couple of studies describing efforts to stabilize C.1086 in native-like trimeric form (Guenaga et al., 2017; He et al., 2018). The study by Geunaga et al., 2017 showed the addition of glycine substitutions in gp41 region of C.1086 NFL in combination with multiple trimer derived changes (Guenaga et al., 2017) to improve the generation of well-ordered trimers. However, immunization studies with such stabilized trimers are yet to be reported. Here, we explored alternative strategies systematically to generate stable, prefusion, native-like C.1086 trimer, compared the immune responses of the design variants in rabbits to monitor if the designs could successfully induce any neutralizing as well as broad V1V2-scaffold specific responses, monitor if optimized changes introduced at the V2 region could maintain native apex integrity with time without affecting the trimer integrity and influence the generation of neutralizing, V1V2-scaffold specific responses. A wildtype (WT) C.1086 gp140 protein that does not maintain the native-like conformation failed to induce autologous neutralizing antibodies (Burton et al., 2019; Kasturi et al., 2017; Styles et al., 2019).

Given the importance of V2 region in regulating the closed-state of the envelope (Cimbro et al., 2014; Guzzo et al., 2018), we carried out an integrated approach using consensus clade C V2 hotspot (V2-HS, res. 166-173 in this study) sequence-guided mutational screening to design UFO-v2-RQH^173^ variant that improved expression of the trimeric fraction and enhanced antigenic properties associated with a well-formed trimer. We analyzed the immune responses elicited by the C.1086 variants in rabbits and showed that the optimized UFO-v2-RQH^173^ variant enhanced induction of V1V2-scaffold specific antibodies and autologous neutralizing responses. UFO-v2-RQH^173^ carried an intrinsic dynamic V2 loop, which otherwise was relatively less dynamic in well-ordered clade A BG505 SOSIP.664 trimer. Furthermore, substituting Y at 173 (the most prevalent variant in HIV-1 sequences), demonstrated decreased ability in displaying native-like V1V2 epitope reactive to V1V2 specific bnAbs relative to UFO-v2-RQH^173^ with time *in-vitro*, despite carrying similar antigenic properties and overall structural organization. UFO-v2-RQY^173^ induced significantly higher V1V2-scaffold specific antibody responses than UFO-v2-RQH^173^, and modestly reduced neutralizing and ADCVI specific functional antibodies than its counterpart UFO-v2-RQH^173^. These results report the development of an optimized clade C trimeric immunogen, UFO-v2-RQH^173^ capable of inducing improved autologous neutralizing and broader V1V2-scaffold specific responses, and highlight the contribution of residues in V2-HS in modulating these two kinds of responses.

## Results

### Screening of C.1086 base variants to select UFO form as the parent construct for stabilization strategies

As initial steps to generate a stable C.1086 trimeric protein, we generated the SOSIP(Sanders et al., 2013), NFL (Native Flexibly Linked with I559P)(Sharma et al., 2015), and UFO (Uncleaved preFusion Optimized with no disulfide linkage SOS between A501 and T605)(Kong et al., 2016) variants (schematic shown in Figure 1A) and compared their ability to form trimers using size exclusion chromatography (SEC). The proteins were expressed in 293F cells and purified by *Galanthus nivalis* lectin (GNL) mediated affinity purification, followed by isolation of trimers by SEC. The majority of the SOSIP variant resulted in aggregates (only 23% trimer proportion) similar to the WT protein and was not characterized further (Figure 1B). Both NFL and UFO variants formed higher proportion of trimers (mean 61% and 50% respectively, Figures 1B and 1C) relative to the SOSIP and WT proteins with net protein yield of 1.8 and 5.5mg/L, respectively. In addition to higher protein yield, the UFO form showed a ∼60-fold improved association kinetics against V1V2 apex specific bnAb, PG16 (Pejchal et al., 2010) resulting in a 44-fold improved binding (K_D_ UFO 252nM, NFL 11µM, Figures 1D, 1G, S1A, S1B, Table S1) against PG16; and comparable binding to other bNAbs (±3-fold) relative to NFL, prompting us to select UFO as the parent design to further improve the trimer.

**Figure 1.**
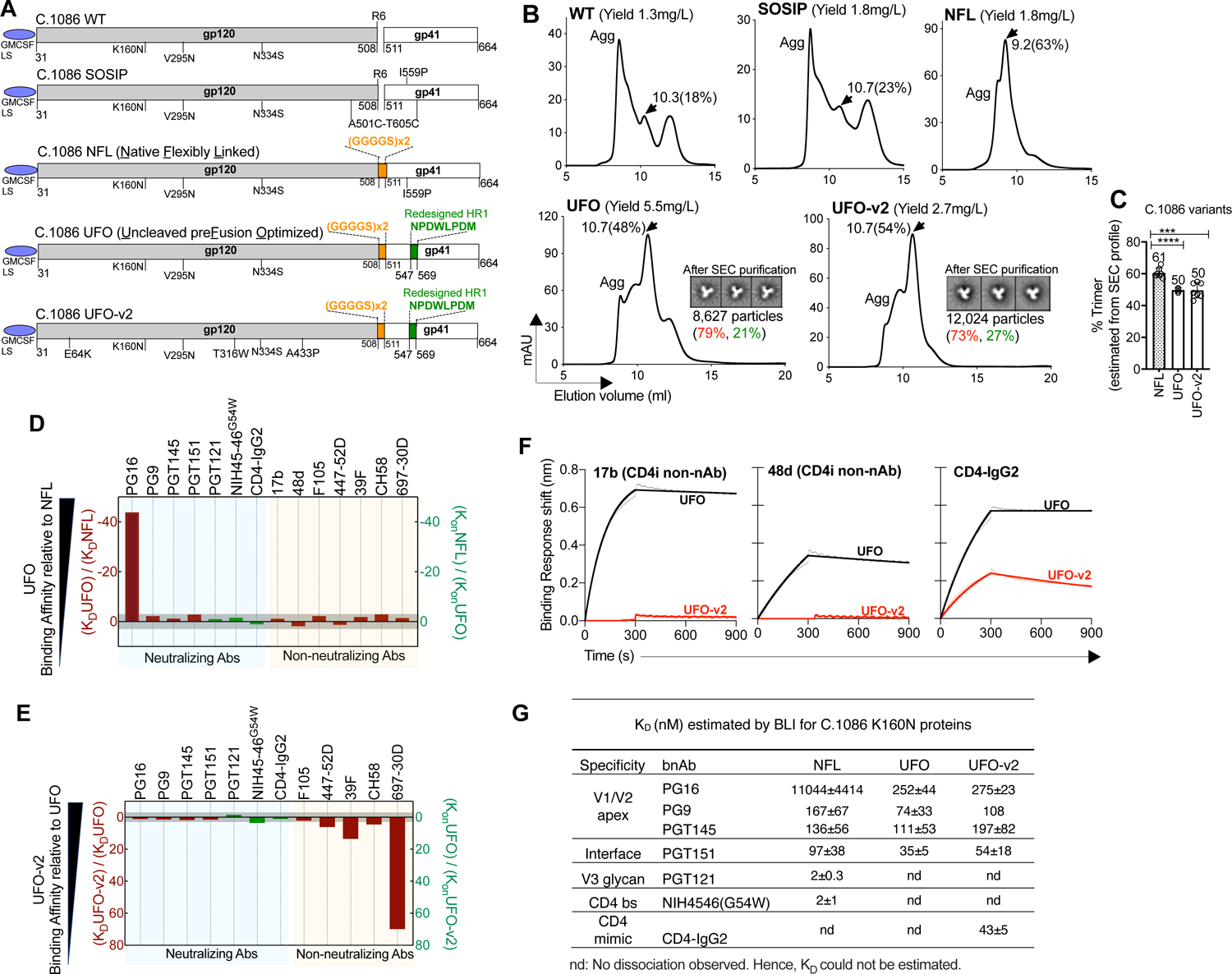
Biophysical characterization of C.1086 base constructs. **(A)** Schematic representation of different C.1086 base constructs tested in the study. GM-CSF leader sequence (LS) was used as secretory signal. (**B)** SEC profiles of 293F expressed, GNL affinity purified C.1086 base constructs. Trimeric peak (black arrow) elution volume and proportion (%, AUC), “Agg” Aggregate/Oligomeric peak. 2D-class averages of the UFO and UFO-v2 trimers (purified after SEC) monitored by negative stain electron-microscopy shown at bottom right of corresponding traces with total particles imaged, %native-like (red) and non-native like malformed trimers (green) indicated. (**C)** Trimeric proportion of C.1086 NFL, UFO and UFO-v2 variants (mean (value indicated on top of bar) ± SD (error bars)) of at-least three independent transfection/purifications, estimated from SEC profiles. Student’s t test (two tailed) for statistical comparisons, *p < 0.05, **p< 0.01, ***p<0.001, ****p<0.000. (**D)** Comparison of binding affinities of UFO and NFL trimers for various envelope specific mAbs. K_D_UFO/K_D_NFL (left y-axis) and K_on_NFL/K_on_UFO (right y-axis, for instances where K_D_ could not be calculated due to no observable dissociation in the experimental setting). Values<1 were inverted and multiplied by −1 for ease of visualization. All plotted values >0 or lower affinity than NFL and vice-versa. −3<Fold-change<3 (gray shaded area) were not considered as significant change. (**E)** Similar plot as described in (C), but comparing UFO-v2 vs UFO. (**F)** Bio-Layer Interferometry (BLI) responses of 200nM C.1086 UFO and UFO-v2 designs to CD4i non-nAbs 17b, 48d, and CD4-IgG2. (**G)** K_D_ (nM) of C.1086 NFL, UFO, UFO-v2 against various bnAbs. Mean ± SD. “nd” refers to “no dissociation” observed, and hence K_D_ could not be calculated. (**D-G)** All binding affinities estimated by BLI. All C.1086 constructs carried K160N/V295N/N334S changes.

### Structure guided mutations in UFO backbone (UFO-v2) decrease exposure of non-neutralizing epitopes and binding to CD4

Although the UFO form improved binding to PG16, it did not reduce binding to non-neutralizing antibodies relative to NFL form (Figure 1D, Table S1). To minimize the exposure of non-neutralizing epitopes, we introduced three structure guided mutations T316W, E64K (de Taeye et al., 2015) and A433P (Kwon et al., 2015) that have been reported in the context of BG505 SOSIP.664 trimer to improve hydrophobic interactions at the apex, restrict CD4 induced conformational changes, and reduce the exposure of off-target V3 region. We named this variant as UFO-v2 (Figure 1A). The WT and NFL-TD (i.e. NFL + “trimer-derived” TD stabilizing substitutions only) forms of the protein have been reported to be insufficient in forming any native-like trimers by negative stain electron microscopy (EM) (Bontempo et al., 2020; Guenaga et al., 2017), while NFL-TD with additional stabilizing glycine substitutions L568G or T569G in HR1 region efficiently formed native trimers (63% and 62%, respectively) by EM (Guenaga et al., 2017). Here, both UFO and UFO-v2 constructs yielded similar proportions of trimers (50% for both variants, Figures 1B and 1C) as assessed by SEC and >70% native-like trimers by EM analyses (79% and 73% respectively, Figures 1B, and S2). However, UFO-v2 displayed markedly reduced binding to V2i 697-30D non-nAb (70-fold) and other non-nAbs (447-52D, 39F, CH58; 6, 14 and 5-fold decrease in K_D_ respectively); along with no apparent binding to CD4i 17b and 48d non-nAbs relative to UFO, supporting reduced exposure of non-neutralizing epitopes (Figures 1E and 1F). UFO-v2 also displayed reduced binding to CD4-IgG2, which however did not affect recognition by CD4 binding site bnAb NIH-45-46^G54W^ (Figures 1E, 1F, S1C, Table S). This could be due to the introduction of E64K substitution, which has been observed earlier to reduce binding to CD4 (Liu et al., 2017). Both UFO and UFO-v2 exhibited similar binding (± 3-fold change in affinity) to neutralizing Abs (Figures 1E and 1G, Table S1). Thus, structure guided mutations introduced in the backbone of C.1086 UFO (i.e., UFO-v2) trimer decreased the exposure of non-neutralizing epitopes.

### Sequence guided mutations at V2 hotspot region improve antigenicity of the V2 apex on C.1086 trimers

Despite the minimized exposure of non-neutralizing epitopes, the UFO-v2 protein exhibited moderate affinity towards V1V2 apex specific bnAbs (K_D_>100nM, Figures 1G, S1C, Table S1). In order to improve binding to this class of bnAbs and thereby stabilize the protein into a closed native-like trimer, we optimized the V2 hotspot region (V2-HS, res 166-173 in the current study) on envelope harboring the signature binding residues of V1V2 apex-specific bnAbs (Bricault et al., 2019) by mutational screening. For this, we identified four positions viz. 166, 170, 172, 173 in C.1086 (UFO-v2) differing from Clade C consensus V2-HS region (n= 22,415 sequences, Figure 2A) and substituted them with either dominant or sub-dominant amino acid present in the consensus sequence; in the form of single, double, triple or quadruple mutants (listed in Figure 2B left). These mutants (referred as UFO-v2 V2-HS mutants) were screened by ELISA to identify variants which enhanced binding to PGT145 and other V1V2 targeting bnAbs.

**Figure 2.**
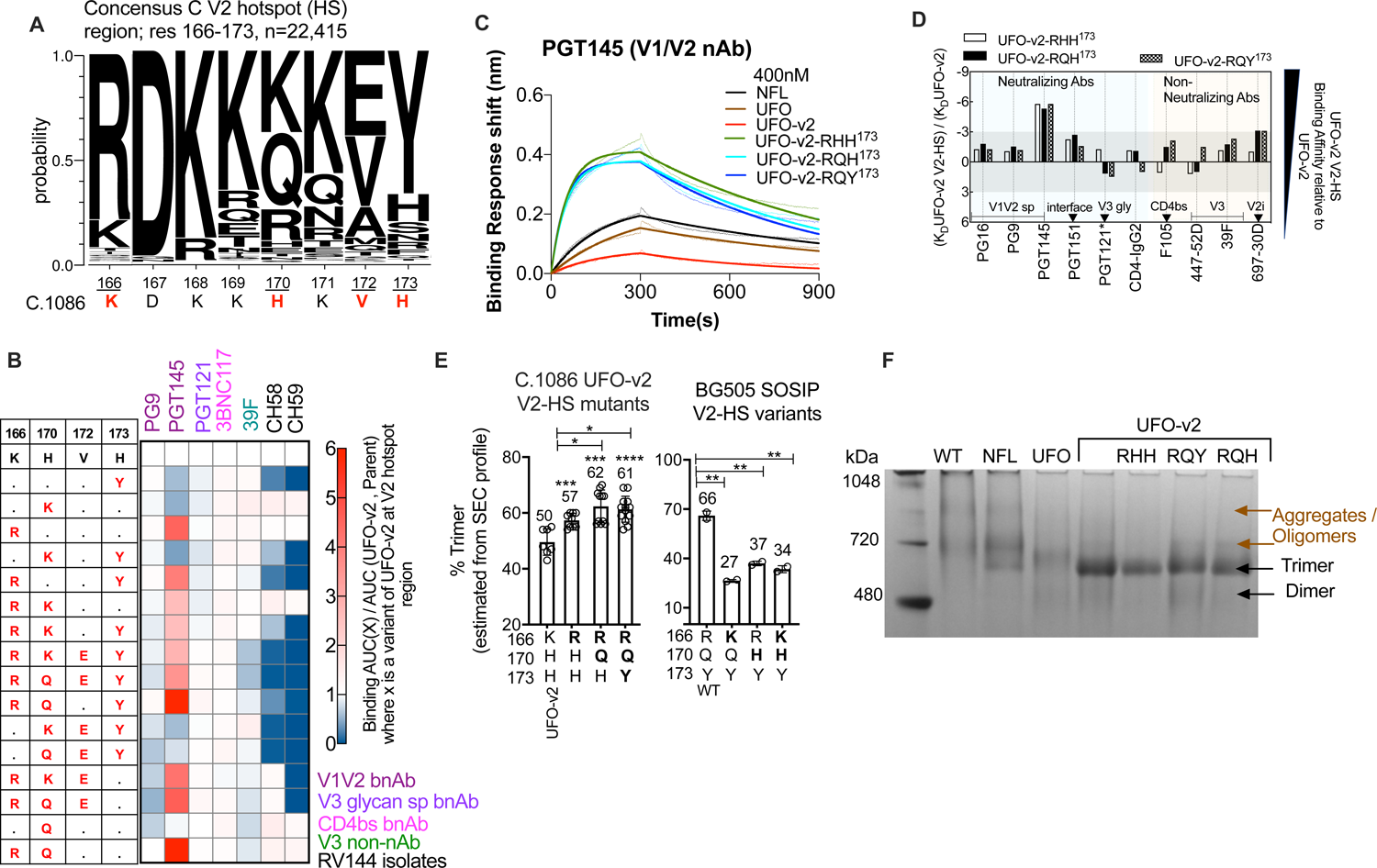
Sequence guided mutations at V2 hotspot region improve properties of native-like trimers. **(A)** V2 hotspot (HS) region (res. 166-173) of Clade C Consensus sequence (n=22415) with corresponding C.1086 region mentioned below. C.1086 residues differing from the consensus sequence highlighted in red. (**B)** Influence of indicated V2 HS modifications in C.1086 UFO-v2 protein on binding to various mAbs. The data are average of more than two independent experiments. (**C)** BLI measurements of different C.1086 purified designs (400nM each) against V1/V2 trimer apex specific PGT145 bnAb. (**D)** Comparison between binding affinities of optimized UFO-v2 V2-HS mutants relative to UFO-v2 (K_D_ UFO-v2 V2-HS / K_D_ UFO-v2) against various envelope specific mAbs. *K_on_UFO-v2 / K_on_ UFO-v2 V2-HS for PGT121 as K_D_ could not be calculated due to no observable dissociation. Values<1 were inverted and multiplied by −1 for ease of visualization. All plotted values >0 or lower affinity than UFO-v2 and vice-versa. ±3-fold change in values (gray shaded area) were not considered significant. (**E)** Trimeric proportion of C.1086 UFO-v2 (left) and BG505 SOSIP.664 V2-HS variants (right, mean (value indicated) ± SD (error bars) of at-least two independent purifications, and estimated from SEC profiles. (**F)** Blue native PAGE (BN-PAGE) of purified C.1086 proteins with molecular weight standard. *, p < 0.05; **, p< 0.01; ***, p<0.001; ****, p<0.0001 (Student’s t test (two tailed), p-value indicated on top of bar (**E**, **left**) corresponds comparison with UFO-v2 which has 166K,170H,173H.

We observed all designs bearing K166R to significantly improve binding to PGT145 with minimal effect on binding to other envelope specific Abs (monitored by ELISA, Figures 2B and S3A). The result was justified as arginine at 166 position directly interacts with the electronegative moieties on PGT145 HCDR3 loop based on the structure solved previously for PGT145-BG505 SOSIP trimer (Lee et al., 2017), and its preference at 166 position for recognition by V1V2 apex directed bnAbs (Bricault et al., 2019). V2-HS mutants viz. K166<u>R</u>/H170<u>Q</u>/<u>H173</u> (referred to as UFO-v2-RQH^173^) and K166<u>R</u>/H170<u>Q</u>/H<u>173Y</u> (referred to as UFO-v2-RQY^173^) showed strongest binding to PGT145 by ELISA (Figures 2B and S3A), and hence were studied further. We used UFO-v2-RHH^173^ (K166<u>R</u>/<u>H</u>170/<u>H173</u>) as a control to examine if K166R had an influence on trimer characteristics. Bio-layer interferometry (BLI) responses monitored for these purified proteins confirmed 5 to 6-fold improved binding to PGT145 (average KD 34nM, Figures 2C, 2D, S1D-F, Table S1) with no significant change in affinity (±3-fold) towards other V1V2 specific bnAbs and other epitope targeting bnAbs and non-nAbs (Figure 2D, Table S1) compared to the parent UFO-v2. Binding affinities of these V2-HS proteins measured against V1V2 apex specific bnAbs PGT145, PGDM1400 (Table S2) were similar to that reported for stabilized C.1086 NFL TD variant (KD 41nM PGT145, 40nM PGDM1400 (Guenaga et al., 2017)), suggesting an alternative strategy of stabilizing C.1086 trimer. Both 173 variants UFO-v2-RQ(H/Y)^173^ displayed similar antibody binding profiles (Table S1, Figure 2D). This was further supported by data from hydrogen-deuterium exchange mass spectrometry (HDX-MS) experiments, showing overall similar backbone amide dynamics of the UFO-v2-RQ(H/Y)^173^ variants (Figure S4). Additionally, UFO-RQ(H/Y)^173^ variants yielded high trimeric protein (∼4mg/L). In summary, consensus C V2-HS sequence guided changes introduced, specifically K166R to generate UFO-v2-RQ(H/Y)^173^ constructs enhanced binding to V2 specific PGT145.

### 166R, 170Q in V2-HS improve protein trimer fraction

We explored whether the enhanced binding to apex-directed PGT145 bnAb by the optimized V2-HS changes was an outcome of improved trimeric protein fraction, besides restoring interaction between PGT145 CDRH3 and 166R (Lee et al., 2017). To answer this, we examined the trimeric proportions of C.1086 UFO-v2 V2-HS mutants from their SEC profiles. We noticed only K166R and H170Q V2-HS substitutions in UFO-v2-RQ(H/Y)^173^ to significantly improve the trimeric fraction of the proteins relative to UFO-v2 (which has K166/H170) and UFO-v2-RHH^173^ (which has 166R/H170) (Figures 2E left, 1B, S3B top). We did not observe any difference in the proportion of trimers between the two 173 (H/Y) variants (UFO-v2-RQ(H/Y)^173^, Figures 2E left, S3B top), indicating that 173(H/Y) did not have any effect on this property of C.1086 trimer. In-order to see if the observed effect of 166R/170Q to increase the proportion of trimers could be translated to a more distant Clade A BG505 envelope, we generated BG505 SOSIP.664 V2-HS mutants at 166 and 170 positions and compared their SEC profiles. We sequentially modified 166R and 170Q present in the WT BG505 sequence to those present in WT C.1086 (contains 166K and 170H) and noticed substantial reduction in the trimeric fraction (Figure 2E right, S3B bottom). These results suggested plausible role of 166R and 170Q in enhancing folding of protein and thereby the proportion of trimers.

We monitored the purity of the C.1086 trimers by blue native PAGE (BN-PAGE). Though the purification steps (after GNL and SEC) yielded >95% trimers (inset in S3B top Fig), we noticed higher order oligomers and dimers by BN-PAGE (Figure 2F) and non-native like malformed trimers by EM (Figure S2) in the purified trimers. The differences in the proportion of unwanted higher-order species among the C.1086 UFO-RQ(H/Y)^173^ variants (by BN-PAGE, Figure 2F) were likely the cause of differences in the proportion of native-like trimers monitored for these variants by NE-EM (S2 Fig). We anticipate additional purification steps involving anion-exchange, hydrophobic interaction chromatography (Verkerke et al., 2016), bnAb, non-nAb based positive and negative selection (Guenaga et al., 2015a) respectively would likely reduce the unwanted species observed in the process. Nevertheless, overall, we found 166R, 170Q V2-HS residues to improve the trimeric proportion of C.1086 and BG505 envelopes.

### Enhanced V1V2 dynamics of C.1086 UFO-v2 trimers compared to BG505 SOSIP.664 trimer

Previous studies have laid the importance of Y173 in influencing the neutralization sensitivity of envelope (Cimbro et al., 2014; Guzzo et al., 2018). Substitution of Y173 to either Ala or Phe 173 altered susceptibility of the envelopes towards neutralization and increasing vulnerability towards adopting an open conformation (Guzzo et al., 2018). To identify any potential structural differences between C.1086 UFO-v2-RQ(H/Y)^173^ trimer variants, we measured their backbone dynamics by HDX-MS. HDX-MS provides a sensitive means of monitoring changes in local structural ordering, particularly involving solvent accessibility and secondary structure of backbone amide groups. To eliminate spurious signals in the HDX-MS experiment, we repurified the proteins by hydrophobic interaction chromatography (HIC) to obtain highly monodisperse trimers. Both C.1086 UFO-v2-RQ(H/Y)^173^ trimers displayed similar HDX profiles for the peptides we could monitor, including res. 176-179 (proximal to 173 position) in the V2 region; indicating no differences in global and local structural organization of the UFO-v2-RQ(H/Y)^173^ trimers; which was indeed consistent with their similar antigenic profiles (Figure S4, Table S1).

A BG505 SOSIP.664 trimer was used as a reference for HDX-MS experiments due to its extensive structural characterization in past studies (de Taeye et al., 2015; Guttman et al., 2014). The homologous peptic peptides between C.1086 and BG505 envelopes present at the gp120-gp41 interface; res. 35-52 and res. 484-501 were observed to be protected (Figures S4, 3A, 3B), which indicated that the trimers were well-folded and native-like(Verkerke et al., 2016). The C.1086 UFO-v2-RQ(H/Y)^173^ trimers showed overall high local structural dynamics at the apex and in portions of gp41 (Figure 3C), and also relative to BG505 SOSIP trimer (Figures 3A, 3B, S4). Notably, the base of V1 ^112^WDESLKPCVKLTPL^126^ and the peptide segment ^176^FYKL^179^ present in the V2 loop of envelope was found to be more dynamic in C.1086 UFO-v2-RQ(H/Y)^173^ than BG505 SOSIP trimer, which is known to have a well-ordered V1V2 apex (Figurse 3A, 3B, S4); C.1086 ^176^FYKL^179^ being the most dynamic (Figures 3A, S4). This increased flexibility of the C.1086 V1V2 region was further supported by observable binding of CH58 (prefers H over Y at 173 position (Liao et al., 2013)) V2p non-Ab to both C.1086 UFO-v2-RQ(H/Y)^173^ trimers (Table S1, Figures S1E and S1F), which was not observed against BG505 SOSIP (has 166R/170Q/173Y) trimer (Bontempo et al., 2020); given the fact that CH58 recognizes a linear epitope on the V2 C β-strand generally displayed on a more flexible V2 antigen such as gp70-V1V2 scaffold that lacks the native display of variable loops (Figure S1G). As expected, binding of CH58 to gp70-C.1086 V1V2-RQ(H/Y)^173^ was higher than its affinity for the closed trimer conformation (Figures S1G and S1H). In summary, His and Tyr variants at 173 position of C.1086 (UFO-v2-RQ(H/Y)^173^) trimers displayed similar overall structural dynamics. The proteins exhibited an inherent higher V2 loop dynamics and accessibility relative to well characterized BG505 SOSIP.664 trimer.

**Figure 3.**
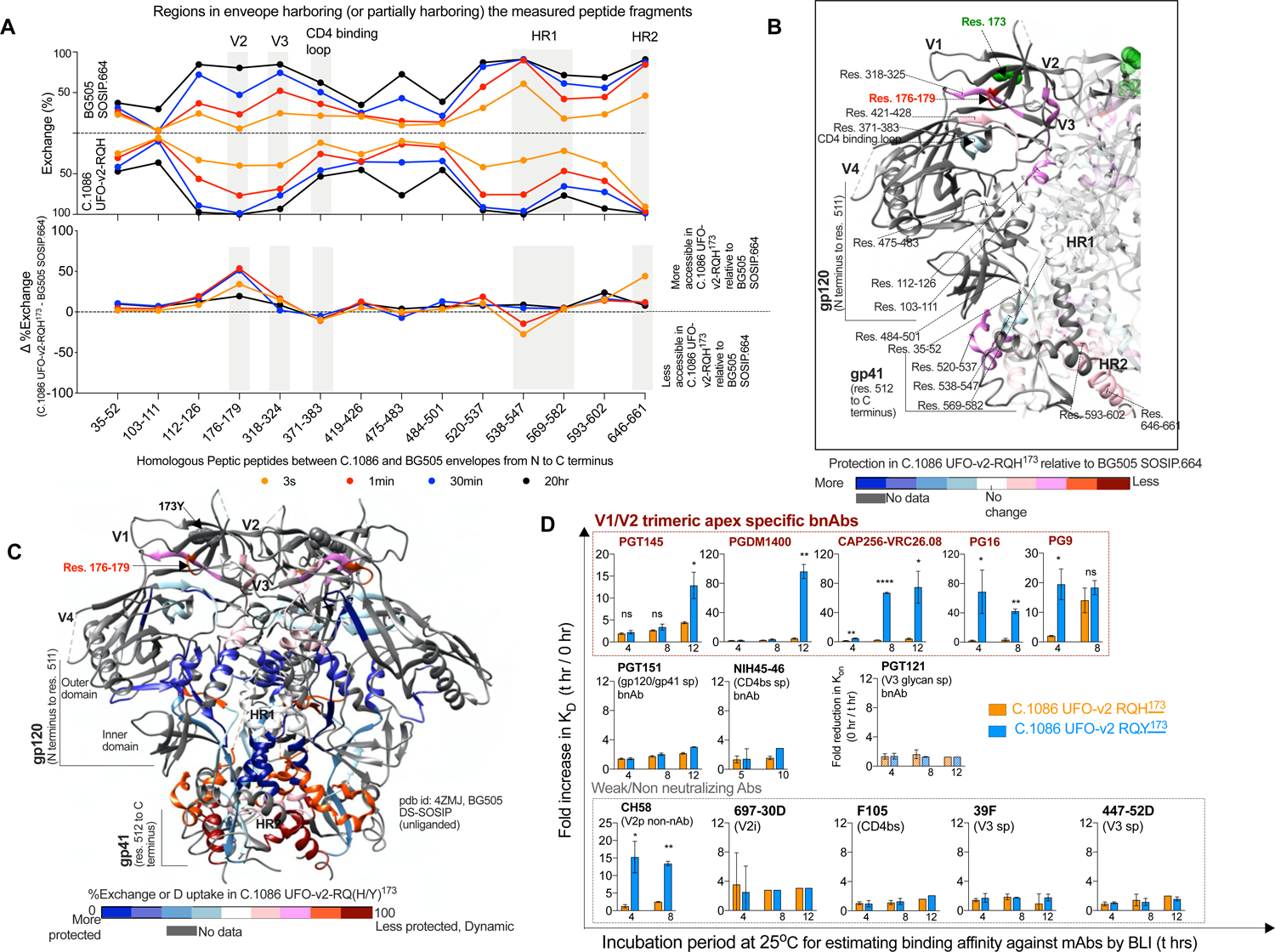
Structural properties of C.1086 UFO-v2-RQ(H/Y)^173^ trimers measured by HDX-MS and time dependent BLI experiments. **(A)** Top, Butterfly plots of C.1086 UFO-v2-RQH^173^ and BG505 SOSIP.664 proteins comparing the %exchange or deuterium uptake for homologous peptide segments (indicated from N to C terminus) of the trimers detected in the experiment at each time point. Bottom, Differences in %exchange or deuterium uptake for the homologous peptide segments of C.1086 UFO-v2-RQH^173^ relative to BG505 SOSIP.664 trimer detected at each time point of the experiment. (**B)** These differences in exchange between C.1086 and BG505 SOSIP trimers have been mapped on the unliganded structure of BG505 SOSIP.DS trimer (pdb id 4ZMJ (Kwon et al., 2015)). (**C)** The exchange kinetics of C.1086 UFO-v2-RQH^173^ were similar to UFO-v2-RQY^173^ (see Figure S4) and hence the average %Deuterium uptake after 1minute of exchange of peptide segments of C.1086 UFO-v2-RQ(H/Y)^173^ trimers were mapped onto the structure of unliganded BG505 SOSIP.DS trimer (pdb id 4ZMJ(Kwon et al., 2015)). (**B,C)** made using UCSF Chimera v1.14 (Pettersen et al., 2004). (**D)** Time dependent fold reduction in binding affinity (K_D_ t hr / K_D_ 0 hr) of C.1086 UFO-v2-RQH^173^ and UFO-v2-RQY^173^ immunogens against various envelope specific mAbs measured by BLI, at 25°C. The purified proteins were incubated for specific time periods (t hr) at 25°C prior to affinity measurements. In case of PGT121, fold reduction in association rate (K_on_ 0hr / K_on_ t hr) was plotted as K_D_ could not be calculated due to no observable dissociation. Bars show Mean, error bars SD of at-least three independent experiments. Student’s t test for statistical comparisons, *p < 0.05, **p< 0.01, ***p<0.001, ****p<0.0001.

### C.1086 UFO-v2-RQH^173^ demonstrated enhanced durability in displaying the epitope reactive to V1V2 bNAbs compared to UFO-v2-RQY^173^ *in-vitro*

Mechanistic understanding of how alterations in the V1V2 region protect or influence native integrity of the apex with time without impacting the trimer integrity is an important but unexplored area. As a preliminary step to probe the influence of 173H and 173Y in the context of C.1086 trimers, we explored any salient differences between UFO-v2-RQ(H/Y)^173^ trimers in their ability to maintain the display of epitopes desirable for a well-ordered trimer, over time. To address this, we incubated UFO-v2-RQ(H/Y)^173^ immunogens at 25°C for time periods ranging from 0 to 15 hrs and measured their binding kinetics against different bnAbs and non-nAbs by BLI. We noticed the largest reduction in binding affinity by V1V2 apex targeted bnAbs after 12hrs incubation at 25°C relative to other immunodominant epitope-specific bnAbs and non-nAbs, which in contrast were marginally affected (Table S2, Figure 3D). Specifically, comparison between UFO-v2-RQ(H/Y)^173^ proteins revealed a dramatic 52, 19, and 5-fold mean reduction in binding affinity of UFO-v2-RQY^173^ against V1V2 specific PG16, PG9, CAP256-VRC26.08 bnAbs, respectively as early as 4hrs, in-contrast to only <2-fold reduction observed for UFO-v2-RQH^173^ (Figure 3D, Table S2). Similar observation was seen for other V1V2 bnAbs PGT145, PGDM1400 (after 12hrs, UFO-v2-RQY^173^ 13, 96-fold reduction respectively; UFO-v2-RQH^173^ 4-fold reduction in affinity for both Abs), and V2p CH58 (after 4hrs, UFO-v2-RQY^173^ 15-fold reduction; UFO-v2-RQH^173^ no observable change in affinity). This time dependent reduction in affinity was observed to be driven by reduced association rate, without much influence on the dissociation rate (Figure S5A, Table S2). Thus, although both the C.1086 UFO-v2-RQ(H/Y)^173^ variants displayed reduced affinity to V2 targeting Abs after 12hrs relative to time 0, this effect was more pronounced for 173Y (UFO-v2-RQY^173^) compared to 173H (UFO-v2-RQH^173^) protein variant. Binding affinities of the 173(H/Y) variants remained similar and unchanged (<3-fold reduction) towards other immunodominant epitopes targeted by neutralizing and non-neutralizing Abs, e.g., 39F, 447-52D non-nAbs targeting the V3 loop sequestered beneath the V1V2 loops (Figures 3D, S5A, Table S2, epitopes mapped on the structure in Figure S5B). V2 B-C β-strands carry the binding footprint of the V1V2 apex specific bnAbs (Bricault et al., 2019) and V2 C strand harbors the 173 position (Figure S5C). CH58 V2p non-nAb targets a linear V2 epitope present in helix/coil conformation and map to V2 C β-strand on trimer (Liao et al., 2013) (Figure S5D). Thus, the observations suggested prolonged incubation of the C.1086 UFO-v2-RQ(H/Y)^173^ immunogens at 25°C primarily affected the display or accessibility of V2 B-C β-strands (based on the Abs tested); and the 173Y trimer variant in this context demonstrated reduced durability in displaying the epitope reactive to V1V2 bNAbs with time compared to UFO-v2-RQH^173^ *in-vitro*, although the two C.1086 UFO-v2-RQ(H/Y)^173^ variants displayed similar antibody binding profiles for V1V2 apex specific bnAbs (Table S1) at baseline time 0.

We next performed dynamic light scattering (DLS) analysis of the HIC-purified trimers to test for the formation of protein aggregates following incubation at room temperature which could have contributed to the decreased binding to V2 specific neutralizing antibodies by the UFO-v2-RQ(H/Y)^173^ trimers *in-vitro.* Following room temperature incubation for 4 hours, we observed the appearance of higher order species in both variants (Figure S5E) but they constituted only a minor proportion. More heterogeneous, larger species were observed to appear in UFO-v2-RQY^173^ at a slightly higher proportion relative to UFO-v2-RQH^173^. Both proteins remained predominantly intact trimers after 4hrs (hydrodynamic radius 70±3Å). It is possible that the increased abundance of higher order species in UFO-v2-RQY^173^ or transient interactions at the apex not observed in HDX data, can lead to decreased accessibility of V1/V2 epitopes relative to UFO-v2-RQH^173^ with time and plausibly influence the immune responses when immunized into animal models.

In summary, V1V2 point alterations 173H and 173Y in the context of C.1086 UFO-v2-RQ(H/Y)^173^ did not alter the structural organization of the proteins, but exhibited differences in their resilience to display V1V2 epitope accessible to native apex targeting bnAbs over time (*in-vitro*) without any effect on the trimer integrity.

### C.1086 UFO-v2 variants induce higher trimer specific responses in rabbits

To monitor the immunogenicity of the optimized C.1086 trimers, female New Zealand rabbits (10-12 weeks old, n=4 per group) were vaccinated subcutaneously with either (1) WT, (2) UFO, (3) UFO-v2, (4) UFO-v2-RQH^173^, or (5) UFO-v2-RQY^173^ proteins (30µg each immunization, 375U ISCOM as adjuvant) at weeks 0, 8, 24 and 40, and bled two weeks after each immunization for serum analyses (Figure 4A). All rabbits, regardless of the immunization group elicited similar binding antibody responses against WT C.1086 gp140, monitored two weeks after the final protein boost (Figure 4B left). All groups manifested moderate (ranging 10^4^-10^6^ ELISA end-point titer without outliers) trimer specific antibody responses after first immunization, followed by a maximum of 237-fold average boost in responses after the second protein, and 14, 27-fold average boost after third and fourth protein respectively (all responses at two weeks post the immunization time, S6A Fig). Encouragingly, UFO-v2-RQH^173^ and UFO-v2-RQY^173^ groups displayed 39-fold higher trimer specific responses than WT (p=0.03 in each case, Figures 4B right and S6B). Although all stabilized trimeric immunogens induced higher trimer-specific binding antibody than WT, comparable responses against the unwanted V3 peptide were observed (Figure 4C left). This resulted in a significantly lower ratio of V3 over total trimer specific responses in all UFO backbone bearing groups relative to WT (Figure 4C right). The induction of similar levels of V3 specific Abs in UFO-v2-RQH^173^ and UFO-v2-RQY^173^ groups was supported by similar V3 backbone dynamics (^318^YATGDIIG^324^ region) of the proteins monitored by HDX experiments (S4 Fig). Thus, a higher proportion of trimer-specific binding Abs was induced by all C.1086 UFO variants, while unwanted V3 targeting Abs normalized to total trimer specific Abs were reduced in the UFO variants relative to WT protein.

**Figure 4.**
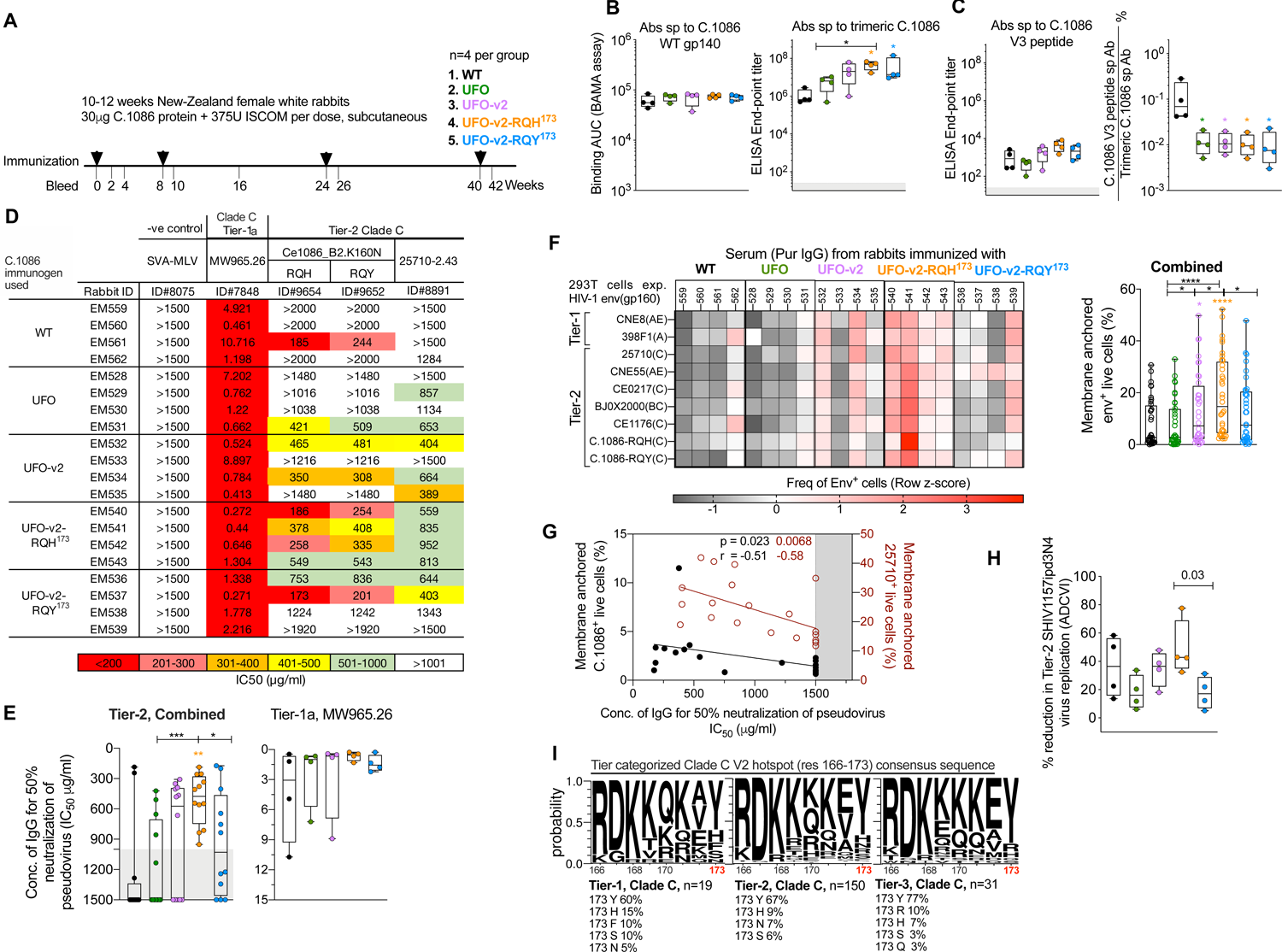
166R, 170Q modifications in V2-HS of UFO-v2 (UFO-v2-RQH^173^) enhance induction of moderate anti-viral responses and binding to membrane anchored tier2 envelopes. **(A)** Schematic overview of the immunogenicity regimen tested in rabbits. (**B)** Serum binding antibody responses against C.1086 WT and trimeric UFO-v2-RQH^173^ proteins measured by Binding Antibody mediated Multiplex Assay (BAMA, indicated by area under the curve (AUC)) and ELISA respectively. (**C)** Serum binding antibody responses against V3 peptide (left) and normalized to total trimer specific responses (right) measured by ELISA. (**D, E)** Neutralizing antibody titer against tier1 and tier2 HIV-1 envelopes. Concentration of purified IgG required to achieve 50% neutralization (IC_50_ µg/ml) is shown. IC_50_ >1480 were assigned 1500 in (**E)**. (**F)** Binding of purified IgG (1µg/ml) from immunized rabbits to broad multi-clade HIV-1 full length envelopes expressed on transiently transfected 293T cells. See Figure S6A for representative flow plot and gating of env^+^ live cells. Row z-scores of the frequency of env^+^ live cells plotted. Env^+^ frequencies measured against each envelope in the panel for each rabbit has been shown in combined plot. (**G)** Spearman’s correlation between binding responses to membrane anchored envelopes (C.1086 and 25710) and concentration of IgG required for 50% neutralization of the corresponding envelope specific pseudovirus. (**H)** Percent reduction in replication of Clade C tier2 SHIV1157ipd3N4 virus in human PBMCs by purified IgGs (250 µg/ml) from various immunized groups (% ADCVI activity). Each data point represents averaged data for a rabbit IgG from three independent experiments, each done in duplicates. (**B-H)** All analyses correspond to serum collected two weeks post final protein boost. Refer to panel (**A)** for color coding of the immunization groups. (**I**) V2 hotspot (HS) regions (res. 166-173) of Clade C tier1(n=19), tier2(n=150) and tier3(n=31) (Rademeyer et al., 2016)mentioned. (**B, E, F, H)** Box and whiskers plots where box extends from 25th to 75th percentile, median indicated by line, minimum and maximum values indicated by whiskers. Statistical comparisons between groups by Mann-Whitney test (*p < 0.05, **p< 0.01, ***p<0.001, ****p<0.0001, p values color coded by group correspond to comparison with WT). All values plotted are the average of at-least two independent experiments.

### 166R, 170Q modifications in V2-HS of UFO-v2 (UFO-v2-RQH^173^) enhance induction of anti-viral antibody responses and binding to membrane anchored tier2 envelopes

We next investigated if the different modifications introduced in C.1086 UFO designs influenced the quality of immune responses induced by vaccination, e.g., generation of (a) neutralizing antibodies, and (b) recognition of membrane anchored diverse tier1 and tier2 full-length envelopes. In addition, we were interested (will be discussed separately below for clarity) in understanding if differences at 173 position i.e., H or Y (as in UFO-v2-RQH^173^ and UFO-v2-RQY^173^) would influence the immune responses, despite exhibiting similar antigenicity. We assayed neutralizing antibody responses against homologous and heterologous tier2 pseudotyped envelopes and binding to membrane anchored gp160s constituting tier1 and tier2 global panel. We used purified IgG to eliminate sporadic low-level background and to be sure that the activity was mediated by IgG. IgG from all animals showed strong neutralizing antibody titers against tier1 envelope MW965.26 (Figures 4D, 4E right, S6F, Table S3A). WT C.1086 has been studied previously to be inefficient in eliciting homologous neutralization titers(Kasturi et al., 2017; Styles et al., 2019). IgG from WT and UFO immunized animals induced only sporadic clade C tier2 responses and increased only marginally in UFO-v2 immunized animals. However, UFO-v2 V2-HS changes in UFO-v2-RQH^173^ resulted in induction of overall better neutralization titers against homologous (C.1086 K160N RQH and C.1086 K160N RQY) and heterologous (25710) pseudotyped viruses than WT, UFO in a combined analysis (p=0.003, 0.0006 respectively) (Figures 4D, 4E left). Additionally, groups immunized with UFO variants showed sporadic induction of moderately better neutralizing responses against tier2 X1632 (Clade G) pseudovirus than the WT (Table S3B).

In an attempt to evaluate the ability of the serum to recognize diverse full-length envelopes, we monitored binding of purified serum IgGs to 293T cells expressing membrane anchored gp160 including tier1 and tier2 Global Panel (deCamp et al., 2014) of envelopes (representative plots in S7A Fig). Reactivity to the envelope panel was poor in WT and UFO groups and increased significantly in UFO-v2 and UFO-v2-RQH^173^ groups with UFO-v2-RQH^173^ showing the highest responses (Figure 4F). As anticipated, the serum antibodies recognizing the membrane anchored envelope correlated positively with neutralization responses (Figure 4G). These results demonstrated the UFO-v2 changes introduced to stabilize the trimer, resulted in induction of better autologous neutralizing and cell surface env binding antibody responses relative to wild-type protein; and the V2-HS changes K166R, H170Q in UFO-v2 (i.e., UFO-v2-RQH^173^) further enhanced these responses.

### 173Y modification in UFO-v2-RQH^173^ marginally reduces induction of functional antibody responses

To understand the contribution of 173Y in the induction of functional antibody responses, we compared neutralizing, antibody-dependent cell-mediated virus inhibition (ADCVI), and cell surface envelope binding activity of IgGs purified from the UFO-v2-RQY^173^ group. In all these cases, we observed modest decrease in responses in the UFO-v2-RQY^173^ group compared to UFO-v2-RQH^173^ group (Figures 4D, 4E left, 4F, 4H), indicating that 173Y modification can negatively influence the induction of functional antibody responses in the context of UFO-v2-RQH^173^. This observation prompted us to investigate if 173Y would be selected over 173H in natural HIV-1 isolates as it elicited weaker anti-viral responses? To address this, we analyzed V2-HS region (res 166-173) of Clade C consensus sequences from tier1 (n=19), tier2 (n=150) and tier3 (n=31) isolates categorized by Rademeyer, C., et al. 2016(Rademeyer et al., 2016). We observed increased occupancy of highly conserved Tyr and reduced occupancy of the sub-dominant His residue at 173 position as the tier level (or difficulty level to neutralize the virus) increased from tier1 to tier3 (Figure 4I, tier1 Y 60% and H 15%, tier2 Y 67% and H 9%, tier3 Y 77% and H 7%). The data corroborated with possible selection of Tyr at 173 position in the viral envelopes to increase its chances to escape the immune system by decreasing tendency to elicit anti-viral responsive Abs, relative to 173H. Presence of Tyr or His at 173 position in the backbone of C.1086 envelope did not influence its infectivity, in the absence of any immune pressure (S7B Fig). It will be important to monitor infectivity of the variants in other cell types. It should be noted that the sequence analysis was limited by the number of tier-categorized clade C sequences available, and 173 was not the only position being influenced by the tier level. In summary, the choice of residue (here, H/Y) at 173 position in C.1086 UFO-v2 V2-HS protein was “capable” of tuning the generation of antibodies with anti-viral functions; an important implication for vaccine design.

### 173Y modification in UFO-v2-RQH^173^ enhances the breadth of V1V2 scaffold specific responses

Immune correlates in the RV144 clinical trial, multiple SIV and SHIV challenge studies in rhesus macaques have shown the positive association of V1V2-scaffold specific antibodies with reduced infection risk (Barouch et al., 2012; Haynes et al., 2012; Roederer et al., 2014; Zolla-Pazner et al., 2019; Zolla-Pazner et al., 2013) ability to generate broad V1V2 responses as an important property of an immunogen. Moreover, UFO-v2-RQ(H/Y)^173^ displayed (a) higher V2 loop dynamics, accessibility of this region on the C.1086 trimers relative to well-ordered BG505 SOSIP, and (b) differences in ability to display V1V2 epitope targeted by specific bnAbs with time; some features likely to influence the elicitation of non-native scaffold V1V2 specific responses. We thereby evaluated if the mutations and V2-HS changes influenced the generation of broad V1V2-scaffold specific Abs in rabbits. To investigate the breadth of V1V2-scaffold reactivity, we assayed binding to gp70-V1V2 scaffold proteins from 16 HIV-1 strains spanning diverse clades by Binding Antibody Multiplex Assay (BAMA) (Zolla-Pazner et al., 2014), in addition to monitoring this against the homologous strain by ELISA. Rabbits immunized with the UFO variants induced higher homologous V1V2-scaffold specific responses than the WT protein; notably highest titers measured by ELISA were seen in UFO-v2 V2-HS groups (Figure 5A). All immunogens generated cross-reactive V1V2-scaffold specific antibodies which were overall significantly higher and broad in the UFO-v2 variants than the WT (Figures 5B, 5C, S6G). Interestingly, UFO-v2-RQY^173^ elicited markedly higher V1V2-scaffold specific responses (Figure 5B), with higher recognition breadth (p<0.001, Figure 5C) compared to UFO-v2-RQH^173^. Rabbit #537 present in UFO-v2-RQY^173^ immunized group, exhibited the highest V1V2 breadth. The results indicated favorable contribution of UFO-v2 in enhancing the cross-reactive V1V2-scaffold breadth, which was further enhanced by 173Y in combination with the V2-HS changes (K166R/H170Q). In-terms of binding responses measured against C.1086 V2-HS (res. 166-180) and V2 cyclic (V2 cyc) peptides, all groups showed weak binding titers, and the V2 cyc specific responses correlated positively with gp70 C.1086 V1V2 titers (S6C-E Fig). The inherent V2 loop dynamics and accessibility of this region on C.1086 trimers measured by HDX-MS may potentially explain the ability of C.1086 UFO-v2 native-like trimers to elicit antibodies that show a broad range of V1V2-scaffold binding. Moreover, differences observed between UFO-RQ(H/Y)^173^ immunogens in terms of their ability to display V1V2 epitope targeted by V1V2 specific bnAbs with time could be one of the factors leading to the differences in immune responses elicited in the study.

**Figure 5.**
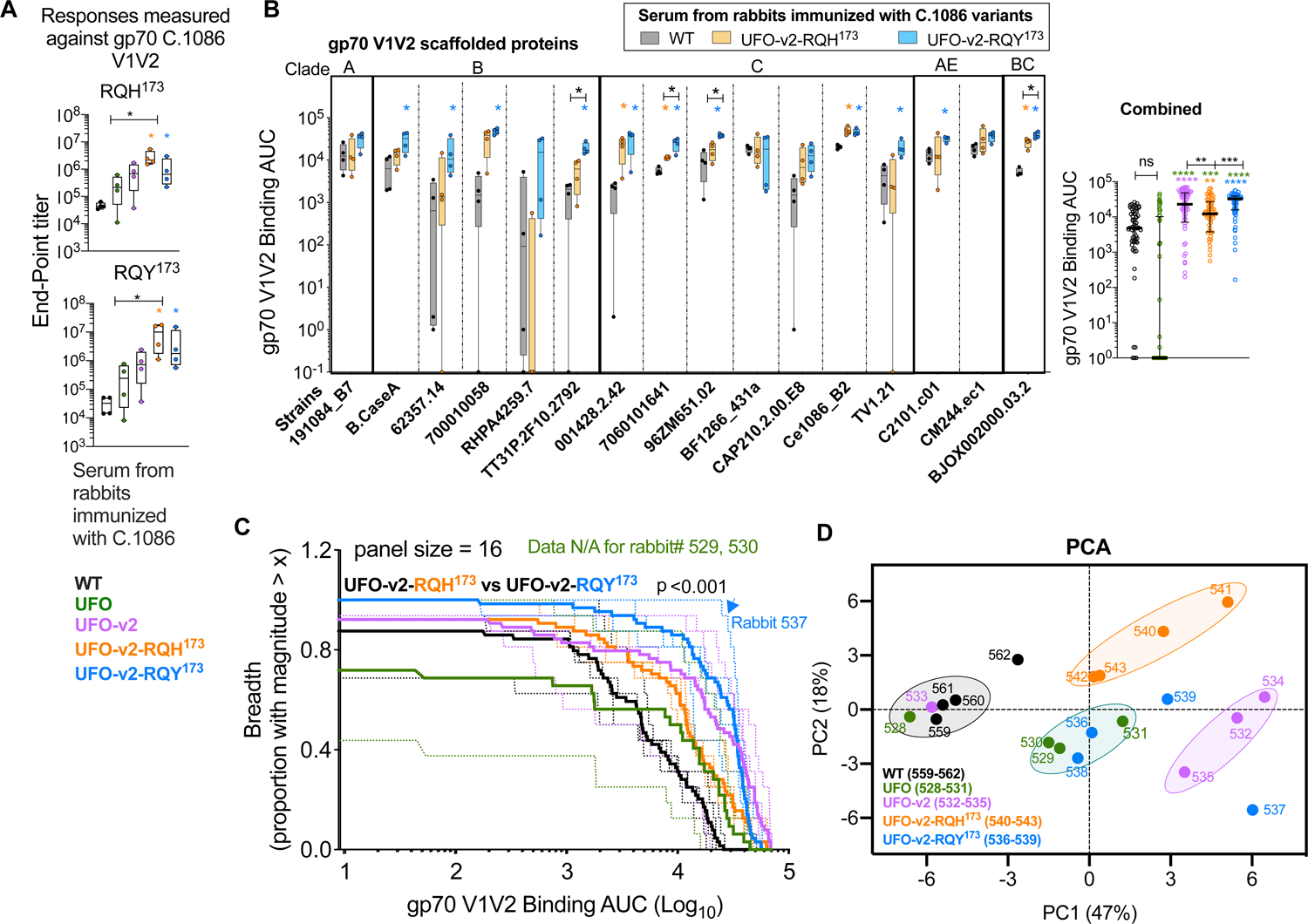
173Y modification in UFO-v2-RQH^173^ enhances the breadth of V1V2 scaffold specific responses. **(A)** Serum binding antibody responses of the immunized groups against gp70 C.1086 RQ(H/Y)^173^ V1V2 proteins by ELISA. Plotted values are the average of two independent experiments. (**B)** BAMA analyses (binding AUC) of serially diluted WT, UFO-v2-RQH^173^ and UFO-v2-RQY^173^ immunized serum to V1V2-scaffolds from 16 cross-clade HIV-1 isolates. Combined plot uses the AUC values obtained against the panel for each rabbit. (**C)** V1V2 Breadth magnitude curves of all immunized rabbits (dotted line) with mean response of a group (bold line). Refer to panel (**A)** for color coding of the immunization groups. Unpaired two-tailed Kolmogorov Smirnov test to see statistical difference between UFO-v2-RQH^173^ and UFO-v2-RQY^173^ groups. (**D)** Principal Component Analyses (PCA using R) of serum characterization data obtained for all immunized rabbits. All analyses correspond to serum collected two weeks post final protein boost. Mann-Whitney test for statistical comparisons between groups (**A, B)** *p < 0.05, **p< 0.01, ***p<0.001, ****p<0.0001. p-values color coded by group correspond to comparison with WT, and in (**B right**) those colored green corresponds to comparison with UFO group. Box and whiskers plots **(A, B left)** where box extends from 25th to 75th percentile, median indicated by line, minimum and maximum values indicated by whiskers, **B right** median indicated by line with interquartile ranges.

PCA analysis of the immunogenicity data segregated UFO-v2-RQH^173^ and UFO-v2-RQY^173^ into two separate clusters, implying true differences in immune responses elicited due to changes at 173 position (Figure 5D). Encouragingly, UFO-v2 and UFO-v2-RQH^173^ groups clustered separately indicating amino-acid differences at K166R and H170Q; previously discussed to be modulating native like trimer folding, resulting in measurably different immune responses as well. However, V2-HS changes and optimizations in the presence of 173Y (UFO-v2-RQY^173^) did not seem to additionally influence immunogenicity and thereby clustered with the ancestor UFO group, rabbit #537 being the outlier. Most notable variables significantly (p<0.05) contributing towards these differences in PCA responses were binding to V1V2-scaffolds of tier2 homologous and heterologous envelops, membrane anchored tier2 25710 env, and neutralization against 25710 pseudotyped virus (Figure S6Hs). Overall, the data suggested an important role of V2-HS residues, particularly 173 position in modulating the immune responses, with nature of the residue as the governing player.

### Trimer stabilizing modifications induce antibodies that compete with CD4bs specific broadly neutralizing antibodies

To gain insights into the epitope specificity of the antibody response induced by these immunogens and determine epitope-specificity induced by our trimer modifications, we performed competition binding experiments between rabbit serum and a panel of well-characterized bnAbs by BLI. The epitope targeted by immunized rabbit IgG was identified by a reduction in binding of C.1086 trimer (or competition) to specific mAb (immobilized on biosensor) in the presence and absence of rabbit IgG isolated two weeks post final protein boost (representative BLI responses shown in Figure 6A). Responses from trimer + pre-bleed IgG served as negative control, while rabbit IgG in presence of only buffer accounted for the background signal. Encouragingly, we observed generation of CD4bs specific Abs (competition with NIH45-46^G54W^) in all UFO-v2 backbone bearing groups (Figure 6B) and correlated positively with homologous C.1086 neutralization IC_50_; concentration of IgG required for 50% neutralization of pseudovirus (p=0.002, r= −0.6, Figure 6C). All rabbits with an IC_50_<400µg/ml also showed modest competition with HJ16 CD4bs bnAb. These results demonstrated early signs of plausible presence of CD4bs neutralizing Abs in the serum. Hence, we created a series of CD4bs mutants in the 1086.C K160N RQH background: N279Q, N280D, S365K, I371A, G458Y, G459(E/P); only N279Q and N280D yielded infectious units. We evaluated neutralization sensitivity of these mutants to sera from the immunized rabbits and observed reduced neutralization of N279Q (in few) and N280D KO mutant viruses (5-fold median reduction in IC_50_ relative to parent C.1086 K160N RQH in all the purified IgGs tested, regardless of the immunization group, Supplementary Table S3c). This suggested the presence of CD4bs Abs in the serum which were different from VRC01-class of bnAbs (VRC01-class CD4bs bnAb activity is dependent on N280(Lynch et al., 2015)); further analyses are needed to identify the fine specificity of the neutralizing activity in these sera. Interestingly, rabbit #537 (IC_50_ against C.1086 RQH=173µg/ml) immunized with UFO-v2-RQY^173^ seemed to have elicited V1V2 specific Abs competing against prototype PG16 (99%), PG9 (86%), VRC26.09 (64%), PGDM1400 (54%) and PGT145 (48%) bnAbs, besides eliciting CD4bs specific Abs in the serum (Figures 6A, 6B). None of the rabbits elicited V3 glycan specific (PGT121 like) antibodies. Most rabbits bearing CD4bs bnAb epitope specificity also exhibited non-neutralizing V3 (39F like), V2p (CH58 like) and V2i (697-30D) targeting antibodies. CAP228-16H V2p Ab recognizes both 173H and 173Y bearing V2 peptides unlike CH58 and exhibit potent ADCC activities(van Eeden et al., 2018). Encouragingly, our data suggested the presence of CAP228-16H like Abs in all rabbits vaccinated with UFO-v2-RQH^173^, in contrast to only one out of four rabbits (rabbit #537) in UFO-v2-RQY^173^ group. Overall, the stabilized UFO variants were able to induce CD4bs-directed and, one instance of V1V2 binding Abs which competed for binding with bnAbs specific for those epitopes.

**Figure 6.**
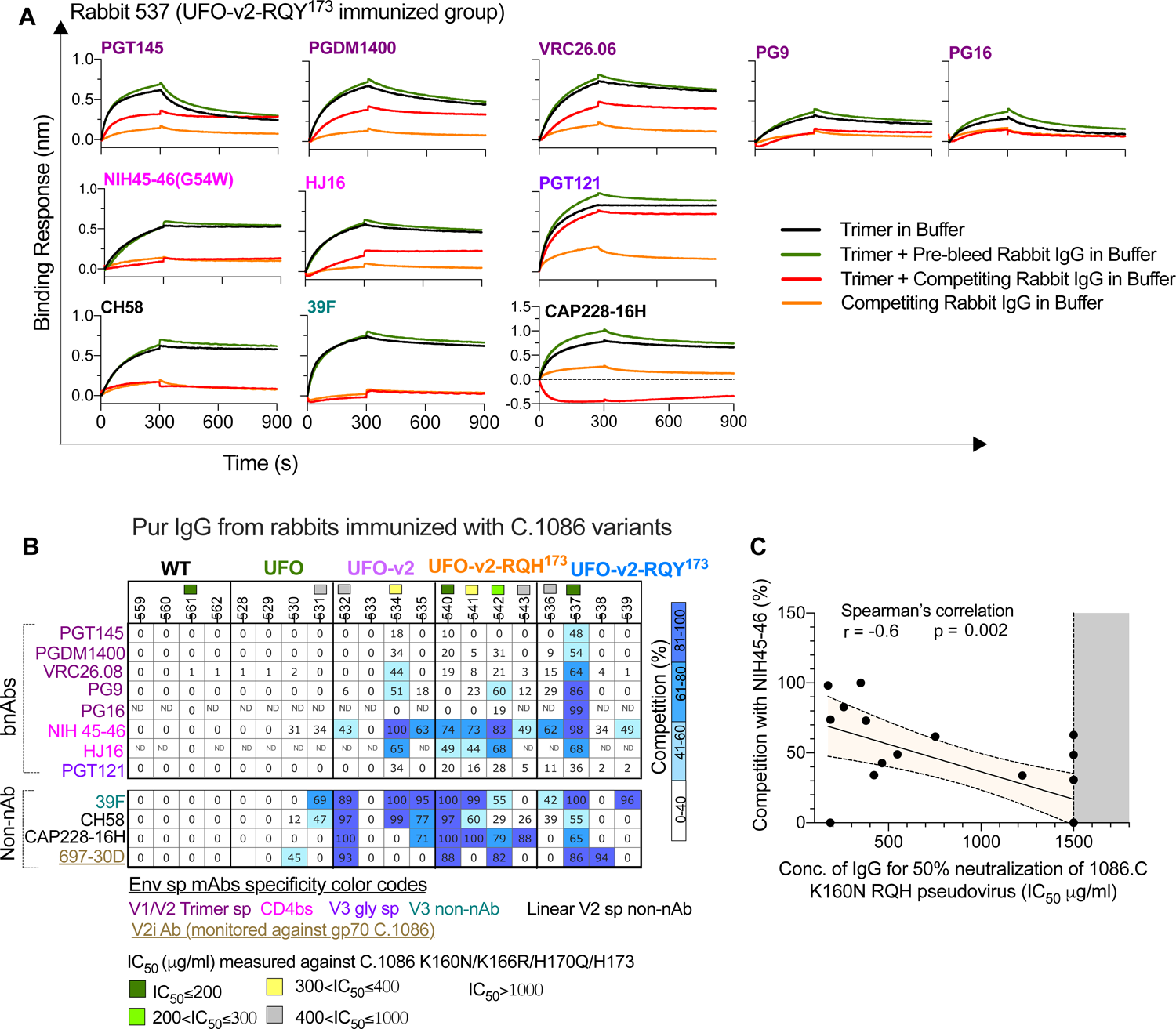
Mapping binding specificity of antibodies elicited in rabbits immunized with C.1086 variants. **(A)** Representative BLI traces of competition binding experiment for purified IgG (rabbit #537) against envelope specific mAbs. (**B) %**Competition of rabbit purified IgGs with envelope specific mAbs to bind C.1086 UFO-v2-RQH^173^ trimers and gp70 C.1086 V1V2 (for 697-30D V2i mAb), represented as heatmap. Purified IgGs from 2 weeks after final protein boost used for analyses (**A, B**). Values <40%, false positive. “ND” denotes not determined due to <1.5-fold difference in signals between background (buffer + purified IgG) and positive (buffer + trimer) control binding signals for the mAb. C.1086 K160N RQH neutralization IC_50_ values (concentration of IgG required for 50% neutralization of the virus) of rabbit purified IgG (filled squares, color coded based on IC_50_ values, see Figure 3D) mentioned above the rabbit codes. Data representative of at-least three independent experiments. (**C)** Spearman’s correlation between C.1086 K160N neutralization IC_50_ and competition (%) with NIH45-46 (estimated in **B**). Shaded orange area represents 95% confidence band. IC_50_ >1480 were assigned 1500 (shaded gray area).

## Discussion

Here, using structure and sequence-guided strategies we engineered the C.1086 envelope protein (referred as UFO-v2-RQH^173^) to elicit antibodies in an animal model with improved functional tier2 anti-viral activities and broad V1V2 scaffold specific binding responses compared to WT protein, where past attempts using WT variants were reported to be inefficient in eliciting any homologous neutralization responses (Burton et al., 2019; Kasturi et al., 2017; Styles et al., 2019). The SOSIP variant of C.1086 mostly yielded aggregates as observed earlier for other envelope sequences (Guenaga et al., 2015a). Disulfide bond-stabilized I201C-A433C substitutions have been widely used to reduce the CD4 triggered conformational change and yield better temporal stability to the HIV-1 trimer than its substitute A433P (Joyce et al., 2017; Kwon et al., 2015). Here, we observed similar binding profiles of DS and 433P variants of C.1086 UFO towards env specific mAbs (data not shown), and we thereby investigated the immunogenicity of A433P in combination with E64K/T316W changes i.e. UFO-v2 variants in rabbits. All UFO-v2 variants showed reduced exposure of non-neutralizing epitopes, which resulted in lower undesired V3 specific immune responses. The UFO-v2 variant also displayed ability to induce higher tier2 virus (envelope) targeting functional antibodies and broad V1V2-scaffold reactive antibodies. We further engineered the V2 region of C.1086 UFO-v2 trimer to improve the abundance of well-formed trimers displaying enhanced binding to V1V2 apex specific bnAbs; PGT145. This was guided by consensus Clade C V2 hotspot sequence based mutational screens and lead to the generation of UFO-v2-RQ(H/Y)^173^ variants, with K166R/H170Q/173(H/Y) changes. The variable V1V2 loops in C.1086 UFO-v2 trimers were more dynamic relative to BG505 SOSIP trimer, which is known to have a well-structured apex V1V2, and hence a potential factor responsible for induction of antibodies by C.1086 UFO-v2 variants targeting flexible V1V2 antigen, such as V1V2-scaffolds.

The native pre-fusion state of the envelope is critically regulated by the V1V2 conformation; consisting of inter V2 β-strand interactions shielded by highly dense glycan network and a cationic cleft in the three-fold trimer axis. Changes at these and proximal positions are likely to influence the dynamics, quaternary epitope and inter-protomer interactions at the apex. For instance, point substitutions M161A, L165A, D167A in the V1V2 region of BG505 SOSIP have been previously reported to form mis-folded trimers (Lee et al., 2017). In the current study, we found R166K, Q170H V2 changes in C.1086 UFO-v2 and BG505 SOSIP envelope proteins to increase the proportion of misfolded higher-order oligomers/aggregates, presumably by destabilizing the quaternary epitope, inter-V2-strand interactions at the apex, and play a role in folding of the envelope. Molecular understanding of how changes at the V2 apex influence the envelope protein folding kinetics is an unanswered area.

Previous studies have highlighted enhanced sulfation of Tyr at 173 position to increase recognition by trimer specific bnAbs, and substituting 173 Tyr to either Ala or Phe to alter neutralization sensitivity specific to an “open” conformation (Cimbro et al., 2014; Guzzo et al., 2018), suggesting important role of 173 position in regulating native envelope conformation. Characterization of the serum elicited by the C.1086 UFO-v2-RQ(H/Y)^173^ immunogens highlighted the importance of sequence-guided V2 hotspot changes in tuning the immune responses, specifically choice of residues His or Tyr at 173 position. The HDX-MS data indicated that the local and global conformation of the trimer was not altered by the H173Y modification for the peptides monitored. Our BLI results suggested, the choice of residues at 173 position (in context with C.1086 UFO-v2-RQ(H/Y^)173^ proteins) could influence the durability in maintaining the V1V2 apex specific antigenicity (monitored at baseline) for an extended time-period, *in-vitro*. The 173H variant (UFO-v2-RQH^173^) displayed the reactive epitope (V2 B-C β-strands) present at the apex of trimer for an extended period, at 25°C *in-vitro* compared to its 173Y counterpart. Located in the V2 C β-strand, residue 173 is adjacent to N156 and its associated glycan, which is spatially proximal to glycan at N160. N156 has been shown to be involved in recognition by V2 apex targeting bnAbs, including PG16, PGT145 (Lee et al., 2017; Pancera et al., 2013). Loss of N156 GlcNAc on the envelope has been reported to alter the inter-V2 strand interaction, form aberrant trimers (Lee et al., 2017) and non-infectious units (observed in this study). Though the amino acid type at 173 position did not alter local structural order of proximal peptide (“FYKL” res. 176-179) monitored by HDX-MS, it is conceivable that the side chain choice at N173 might influence the disposition of the glycan chain at N156 and its associated interactions with the V1V2 trimer specific bnAbs; 173H favoring this interaction over the 173Y in the context of C.1086 UFO-v2-RQ(H/Y)^173^ trimers. Additionally, it is possible that amino acid type at this position may influence native-like trimer’s ability to form higher order oligomers such as larger assemblies of trimers when the proteins are incubated at 25°C for an extended time-period, and occlude/alter accessibility of the V1V2 epitope for recognition by V1V2 trimer specific bnAbs. Such an interaction would be consistent with (a) the formation of larger and more heterogeneous species over time by UFO-v2-RQY^173^, as monitored by DLS at 25°C at relatively higher frequency than UFO-v2-RQH^173^. This would agree with the time dependent V1V2 trimer specific affinity differences; particularly against PGT145, measured for the UFO-v2-RQ(H/Y)^173^ proteins by BLI. These structural differences, as one of the potential factors, may help explain some of differences in immune responses observed for the 173 position variants, *in-vivo* in this study. Accessibility of the V1V2 quaternary epitope present at the apex of the trimer could influence the induction of antibodies with anti-viral functional activities e.g. neutralization, ADCVI, ADCC responses; and V1V2-scaffold specific binding responses, and could influence efficacy of the immunogen in an animal challenge study. Increased V1V2-scaffold Abs have been correlated with reduced risk of viral acquisition and anti-viral activities e.g. ADCC. However, UFO-v2-RQY^173^ immunized group exhibiting highest V1V2-scaffold binding Abs showed minimal anti-viral ADCVI activity. This could be due to (a) differences in the assays, or (b) nature of the V1V2-scaffold Abs induced by 173Y and other studies, including those generated by 173H variant which needs in-depth study.

In summary, we generated clade C C.1086 trimer capable of simultaneously inducing functional autologous tier2 virus reactive antibodies as well as broad V1V2-scaffold specific responses in rabbits. In addition, our results highlighted the influence of V2 residues 166R, and 170Q in the formation of well-folded C.1086 and BG505 trimers in solution, His or Tyr at 173 position in altering durability in displaying V1V2 epitope reactive to V1V2 bNAbs with time *in-vitro*, and in combination with the highly dynamic feature of the V1V2 region of C.1086 trimers to modulate the induction of V1V2-scaffold specific responses and functional antibody responses. Identification of 173(H/Y) like residues by high throughput screening and sequence analyses across various regions of the envelope surface may improve our fundamental understanding of linking relationship between fine-tuned antigen design and immune outcome.

## Materials and Methods

### Immunizations in rabbits

Five C.1086 immunogens viz. WT, UFO, UFO-v2, UFO-v2-RQH^173^, UFO-v2-RQY^173^ were tested for their immunogenicity responses in female New-Zealand white rabbits (10-12 weeks old, n=4 per group). The rabbits were housed and immunized at Covance Laboratories, Inc., Denver, PA, USA in compliance with IUCUC protocol 0065-18. Each rabbit was immunized subcutaneously with group specific protein (30μg/dose) on the neck (dorsal area), formulated with 375U of ISCOM as adjuvant (from Darrel J. Irvine, Howard Hughes Medical Institute) on weeks 0, 8, 24 and 40. Serum was collected before (pre-bleed) and two weeks after each immunization to monitor the antibody responses.

### Design of C.1086 constructs and mutagenesis

All C.1086 proteins generated in the study correspond to 31-664 residues (HxB2 numbering) of C.1086 sequence (Genebank id FJ444392.1) and contain K160N (improve binding to PG9 bnAb), V295N (2G12 binding) and N334S (improve binding to PGT121 and 10-1074) mutations. C.1086 WT (^508^RRRRRR^511^ or ^508^R6^511^ to increase furin cleavage efficiency), SOSIP (A501C/T605C, ^508^RRRRRR^511^)(Sanders et al., 2013), NFL (Native Flexibly Linked, I559P, ^508^(GGGGS)2^511^)(Sharma et al., 2015), UFO (Uncleaved preFusion Optimized, ^508^(GGGGS)2^511^, ^547^NPDWLPDM^569^, no disulphide linkage A501C-T605C)(Kong et al., 2016), UFO-v2 (^508^(GGGGS)2^511^, ^547^NPDWLPDM^569^, E64K/T316W/A433P) envelope inserts with GMCSF leader sequence (MWLQGLLLLGTVACSIS) were synthesized by GenScript and subcloned between ClaI and NheI sites of pGA1 vector (Kan^R^). For generating V2 hotspot (HS) mutants, residues 166-173 in V2 hotspot region of C.1086 differing from the Clade C Consensus sequence i.e. positions 166, 170, 172 and 173 were mutated to either dominant and/or sub-dominant amino acids present in the consensus sequence. Mutagenesis was done by inverse PCR using non-overlapping forward and reverse primers. The mutant codon was present at the 5’ end of the forward primer(Jain and Varadarajan, 2014). pGA1 vector (Kan^R^) expressing C.1086 UFO-v2 was used as the parent/template for V2-HS PCR reactions. NEB® 5-alpha E. coli cells (NEB, catalog no C2987H) and Sanger sequencing were used to transform, screen and confirm the positive clones respectively. Mutant pCDNA3.1 full length C.1086 K160N envelope bearing plasmids (used in neutralization mapping assays) were generated by introducing mutant codons as described above using pCDNA3.1 C.1086 K160N vector (Amp^R^) as template for PCR reactions and transformation of the blunt ligated mixture in NEB® Stable Competent *E. coli* cells (NEB, catalog no C3040H, 30°C). To generate gp70 C.1086 V1V2 RQ(H/Y)^173^ constructs, gp70 (design of gp70 V1V2 has been described previously (Zolla-Pazner et al., 2014)) was synthesized by Genescript in frame with C.1086 V1V2 (res. 120-204; K166R/H170Q for C.1086 V1V2 RQH^173^ and K166R/H170Q/H173Y for C.1086 V1V2 RQY^173^) at its C terminus, and GMCSF leader sequence followed by 6xHis tag in-frame at its N terminus, and sub-cloned between ClaI and NheI sites in pGA1 vector. The BG505 SOSIP envelope insert (Genebank id ANG65466.1, res. 31-664, A501C/T605C/T332N, ^508^RRRRRR^511^,(Sanders et al., 2013)) was synthesized by Genescript with GMCSF leader sequence at its N terminus and sub-cloned between ClaI and NheI sites of pGA1 vector. To generate V2-HS mutants of BG505 SOSIP, we used inverse PCR using non-overlapping forward and reverse primers as described above for generating C.1086 V2-HS mutants.

Gp70_sequence:^N_terminus^VYNITWEVTNGDRETVWAISGNHPLWTWWPVLTPDLCMLALSGPPH WGLEYQAPYSSPPGPPCCSGSSGSSAGCSRDCDEPLTSLTPRCNTAWNRLKLDQVTHKSSEGFYVCPGSHRPREAKSCGGPDSFYCASWGCETTGRVYWKPSSSWDYITVDNNLTTSQAVQVCKDNKWCNPLAIQFTNAGKQVTSWTTGHYWGLRLYVSGRDPGLTFGIRLRYQNLGPRVPIGPNPV LADQLSLPRPNPLPKPAKSPP^C_terminus^.

### Purification of protein

The envelope gp140 glycoproteins cloned in Kan^R^ pGA1 plasmid were transiently expressed from Expi293F cells using the Expifectamine^TM^ 293 transfection kit (ThermoScientific) per manufacture’s protocol and grown at 37°C, 8% CO_2_ at 130rpm. The supernatant was harvested 72hrs after transfection in presence of EDTA free protease inhibitor (Millipore Sigma, catalog no 11836170001) and affinity purified by *Galanthus nivalis* lectin agarose (Vector Labs, catalog no AL-1243-5, pre-equilibrated with PBS). Bound protein was eluted in presence of 1M methyl α-D-mannopyranoside (Sigma). The protein was dialyzed against PBS and subjected to size-exclusion chromatography using a Superdex 200 Increase 10/300 GL (Sigma, GE Healthcare product) column on an Akta^TM^ Pure (GE) system. The trimeric peak was collected, concentrated using Amicon Ultra-4, MWCO 100kDa, and quantified by BCA assay (Pierce™, ThermoScientific). To obtain highly monodisperse trimeric population for HDX-MS and DLS experiments, an additional hydrophobic Interaction chromatography (HIC) based purification (described by Verkerke, et. al., 2016 (Verkerke et al., 2016)) was done on SEC purified protein samples prior to experiments. The proteins purified from SEC were dialyzed against a high salt buffer A (2M NH_4_SO_4_, 100mM NaH_2_PO_4_, 0.02% sodium azide; pH 7.4), bound to a pre-equilibrated (high-salt buffer A) Hitrap Phenyl HP column (Cytiva, catalog no 17519501), and eluted using a linear gradient (0-100%) of low salt buffer B (0.1M NaH_2_P04, sodium azide 0.02%, pH 7.4) using an Akta^TM^ Pure system (GE). The eluates containing trimeric peak of interest were pooled, dialyzed against PBS, concentrated and quantified. In all cases, the trimeric status and purity of the proteins were confirmed by BN-PAGE (NuPAGE™, 4-12% Bis-Tris Protein Gels, ThermoScientific). The gp70 V1V2 proteins were purified by His based affinity purification, using HisPur™ Ni-NTA resin (ThermoFisher, catalog no 88221), as described in the manufacturer’s protocol. Briefly, the supernatant was diluted (1:1) in equilibration buffer (20mM sodium phosphate, 300mM sodium chloride, 10mM imidazole in PBS, pH 7.4) and bound to HisPur™ Ni-NTA column (pre-equilibrated with equilibration buffer). The column was washed with wash buffer (25mM imidazole in PBS; pH 7.4) to remove non-specific loosely bound proteins, followed by elution of the protein in presence of 250mM imidazole in PBS, pH 7.4. The eluted protein was dialyzed against PBS, concentrated using Amicon Ultra-4, MWCO 10kDa, and quantified by BCA assay. The purity of the protein was monitored by SDS-PAGE, and western blot. BG505 SOSIP T332N protein used as reference in mass spectrometry-based experiments was expressed from pPPI4 vector (kindly provided by Dr. John P. Moore, Cornell University, NY, USA) in 293F cells and purified as described above for C.1086 envelope gp140 constructs. For all assays using rabbit purified IgGs, IgG was purified from immunized rabbit serum using Pierce™ Protein A IgG Purification Kit (ThermoFisher, cat. No. 44667) and dialyzed against PBS, as per manufacturer’s instructions. The protein was concentrated using Amicon Ultra-4, MWCO 30kDa, and quantified by BCA assay.

### Negative stain Electron Microscopy (NS-EM)

Protein samples were diluted to 0.02 mg/ml, applied to a carbon coated Cu400 grid, and stained with 2% (w/v) uranyl formate for 30-60 s. Data were collected on an FEI Tecnai Spirit T12 transmission electron microscope operating at 120 keV and equipped with a Tietz TVIPS CMOS camera. A magnification of 52,000x was used, resulting in a physical pixel size at the specimen plane of 2.05 Å. Data was collected using the Leginon software package (Suloway et al., 2005), and processing (particle picking and stack creation) was performed in Appion (Lander et al., 2009). Two-dimensional classifications were performed using MSA/MRA method described by (Ogura et al., 2003). Class averages were inspected manually and compared to previously published 2D class averages of HIV-1 Envelope SOSIP trimers (for example see (de Taeye et al., 2015)).

### Enzyme-linked Immunosorbent Assay (ELISA) to screen V2 hotspot (HS) mutants for enhanced binding to V1V2 specific bnAbs

C.1086 V2 hotspot envelope mutant supernatants collected 48hrs after transient transfection of 293T cells were screened for increase in binding to PGT145 than the parent C.1086 UFO-v2. Envelope supernatants were immobilized onto ConA (Sigma, catalog no C2272, 25μg/ml in HEPES buffer i.e.,10mM HEPES, 151mM NaCl, 4.7mM KCl, 2mM CaCl2, 1.2mM MgCl2, 7.8mM Glucose, pH 8.5) coated ELISA maxisorp microtiter plates (ThermoFisher, catalog no 439454) at RT for 2hrs. The plates were blocked with 5% BSA + 4% Whey in PBS, RT, 1hr. Serially diluted envelope specific mAbs PGT145, PG9, PGT121, 3BNC117, 39F, CH58, CH59 Abs were added to the plates at RT, 1hr, after which plates were incubated with mouse anti-human biotin (BD Biosciences, catalog no 555785, 1:5000 dilution in 4% Whey buffer) and streptavidin horseradish peroxidase (Vector Labs, catalog no SA5004, 1:1000 dilution) as secondary and tertiary Abs respectively for 1hr at each step, RT. The plates were washed with PBS containing 0.05%Tween-20 (6 times) after each incubation step. The plates were developed with 3,3’,5,5’-tetramethylbenzidine chromogenic substrate solution (KPL TMB Peroxidase substrate, SeraCare) in dark. The reactions were quenched with phosphoric acid. Equal concentrations of the envelope proteins (in supernatants) were added to the ELISA plates. The concentration was estimated by densitometric analyses of the envelope specific bands monitored by western blot of SDS-PAGE (+DTT) gel. ID6 (NIH AIDS reagents program, 0.5μg/ml), goat anti-mouse-HRP (SouthernBiotech, catalog no 1030-05, 1:40,000 dilution in 4% Whey buffer) were used as primary and secondary Abs for western blot. A_450nm_ recorded from the developed ELISA plates were analyzed to measure binding area under the curve (AUC) against each mAb tested.

### ELISA to estimate C.1086 specific responses in immunized serum

To monitor trimer specific responses, ELISA microtiter plates were coated with trimeric C.1086 UFO-v2-RQH^173^ protein at 2μg/ml (in PBS), overnight at 4°C. The plates were blocked with 5% BSA + 4% Whey in PBS, RT, 1hr. The plates were incubated with serially diluted serum at RT, 2hrs, followed by incubation with goat anti-rabbit HRP (SouthernBiotech, catalog no 4010-05, dilution 1:4000), RT, 1hr. The plates were washed with PBS containing 0.05%Tween-20 (6 times) after each incubation step and developed as described above. To monitor C.1086 V3 responses, ELISA plates coated overnight with V3 peptide (res. 296-33, 1μg/ml in PBS) were used. gp70 V1V2 specific responses were monitored by using plates coated with 2μg/ml His-tag purified gp70 C.1086 V1V2 RQH (K166<u>R</u>/H170<u>Q</u>/<u>H</u>173) and RQY (K166<u>R</u>/H170<u>Q</u>/H173<u>Y</u>) (Zolla-Pazner et al., 2014) proteins. V2 hotspot specific responses were monitored using V2 peptides, V2-HS 173H (^166^KDKKHKV<u>H</u>ALFYKLD^180^) and V2-HS 173Y (^166^RDKKQKV<u>Y</u>ALFYKLD^180^) (1μg/ml in PBS coated on ELISA microtiter plates). To monitor V2 responses cyclized C.1086 V2 (cV2, synthesized by GenScript) ^157^CSFNATTELKDKKHKVHALFYKLDVVPLNGNSSSSGEYRLINC^196^ was used at 1μg/ml in PBS for coating. A_450nm_ recorded from the developed ELISA plates were analyzed to measure end-point titre associated with the serum responses.

### Binding antibody multiplex assay (BAMA)

Binding antibody multiplex assay (BAMA) of the serum was performed as described previously (Tomaras et al., 2008; Zolla-Pazner et al., 2014). Briefly, serially diluted rabbit serum (5-fold dilution starting with 1:80) were tested for binding to color-coded beads by Bio-plex (Biorad). The beads were coated with avi tagged C.1086 WT, UFO-v2-RQH^173^, UFO-v2-RQY^173^ gp140 and gp70 scaffolded proteins with V1V2 grafted from B.CaseA, 7060101641, CM244.ec1, TV1.21, 001428.2.42, CAP210.2.00.E8, C2101.c01, BJOX002000.03.2, BF1266_431a, 96ZM651.02, RHPA4259.7, Ce1086_B2, 62357.14, 700010058, 191084_B7, and TT31P.2F10.2792 HIV-1 strains. Biotinylated anti-rabbit IgG was used as secondary antibody followed by streptavidin conjugated fluorophore to monitor mean fluorescence intensity (MFI) signal at the dilutions tested. This was subsequently used to calculate binding area under curve (AUC) for analyses. Instances satisfying the following criteria were considered positive, i.e. (a)MFI at 1:80 dilution >100, (b) MFI at 1:80 > antigen specific cut-off (95th percentile of all prebleed for the study for each antigen), and (c) MFI > 3-fold that of the matched baseline or pre-bleed samples, both before and after blank bead subtraction.

### Binding of purified serum IgGs to cell surface bound gp160 envelopes

293T cells were transfected with full-length envelope expressing plasmid using lipofectamine2000 (ThermoFisher, catalog no 11668027) as per the manufacturer’s protocol. The cells were harvested 36 hours after transfection and incubated with 1µg/ml of purified rabbit IgG (week 42 i.e. two weeks after the final protein boost) for 45minutes, RT in presence of live/dead fixable stain (ThermoFisher, catalog no L34975). Envelope specific mAbs PGT145 (1µg/ml), PGT121 (0.5µg/ml), PG9 (0.5µg/ml), PG16 (1µg/ml) were used instead of rabbit IgG as controls to monitor expression of envelope on the cell surface. Goat anti-rabbit IgG PE (SouthernBiotech, cat. no. 4030-09, 1:1000 dilution) was used as secondary antibody, 30minutes, RT for monitoring binding to immunized rabbit IgG. To monitor binding to mAbs, mouse anti-human biotin (BD Biosciences, catalog no 555785, 1:5000 dilution) was used as secondary antibody followed by streptavidin conjugated PE (BD Biosciences, cat. no. 554061, 1:5000 dilution), 30minutes at RT each step. The cells were washed twice at 1500rpm, 5minutes, RT with BD FACS buffer after each incubation step. The samples were fixed in presence of 1% paraformaldehyde (in PBS) and acquired on LSRII flow cytometer. pSV-A-MLV-envelope (NIH AIDS Reagent program) was used as negative control. Binding of purified rabbit IgG to this was used as reference to gate binding of corresponding rabbit IgG to live envelope positive cells. All envelope plasmids tested in the study constituting the global panel of HIV-1 envelopes were obtained from NIH AIDS Reagent program.

### Bio-layer Interferometry (BLI)

The assay was done in 384 well format on Octet Red384 platform, Pall ForteBio. Envelope specific monoclonal Abs (NIH-AIDS Reagent program) were immobilized onto anti-human Fc coated biosensors at 5µg/ml. The purified protein was used as analyte (1000nM-12.5nM). Env-Ab association (k_on_) was monitored for 300s, followed by dissociation (k_off_) for 600s. 10X Kinetics Buffer, Pall ForteBio was used as buffer for all steps in the assay. All steps were performed with agitation at 1000rpm. K_d_ (dissociation constant = k_off_/k_on_) was estimated by globally fitting the reference (buffer only) subtracted sensograms to a 1:1 binding model using ForteBio Octet Data Analysis v9 software. For time dependent affinity estimates, purified trimeric proteins C.1086 UFO-v2-RQH**^173^**, UFO-v2-RQY**^173^** were serially diluted (800-12.5 nM) in 10X Kinetics Buffer, Pall ForteBio and incubated at room temperature (25°C) for 0, 4, 5, 8, 10, 12hrs, prior to affinity measurments by BLI.

For competition assays to monitor epitope specificity of the serum by BLI, IgG was purified from immunized rabbit serum. IgGs from pre-bleed and two weeks after the final protein boost were purified. To identify the epitope on C.1086 envelope protein targeted by rabbit purified IgG, we monitored competition between rabbit IgG and envelope specific mAb to bind the epitope in question; e.g. competition between purified rabbit IgG and HJ16 to bind CD4 binding site on the protein. Anti-human Fc biosensor was immobilized with 5µg/ml of envelope specific mAb. Purified C.1086 UFO-v2-RQH^173^ (800nM) was pre-incubated with 6.4µM of purified rabbit IgG in 10X Kinetics Buffer, Pall ForteBio at 4°C, 1hr. This was used as analyte to monitor association with the immobilized biosensor for 300s and dissociation for 600s. (A) Protein only was used to estimate the maximum binding signal with desired mAb. (B) Protein + pre-bleed IgG served as negative control, while (C) signal from buffer + immunized purified IgG was used to monitor background or non-specific signal. (D) signal from protein + immunized purified rabbit IgG. % competition was calculated as 100*(D-C)/Mean(A, (B-C)). No non-specific interaction was observed between mAb immobilized biosensor and purified IgG alone. Area under the curve of the BLI sensogram (t=0 to 600s) was used as “signal” for analyses. Negative traces were inverted to calculate AUC. Competition >40% was considered positive based on examination of signals observed in (B), (D) and (C). Cases where AUC_Signal B_ > 1.5x AUC_Control signal C_ were considered for analyses. All steps were performed with agitation at 1000rpm.

### Neutralization assay

Neutralizing antibody activity in serum and purified IgG samples was measured in 96-well culture plates by using Tat-regulated luciferase (Luc) reporter gene expression to quantify reductions in virus infection in TZM-bl cells. TZM-bl cells were obtained from the NIH AIDS Research and Reference Reagent Program (catalog no 8129). Assays were performed with HIV-1 Envelope pseudotyped viruses produced in 293T cells essentially as previously described (Montefiori, 2009). Serum samples were heat-inactivated at 56°C for 45 minutes prior to assaying; purified IgG were not heat-inactivated prior to assaying. Samples were diluted over a range of 1:20 to 1:43740 in cell culture medium and pre-incubated with virus (∼150,000 relative light unit equivalents) for 1 hr at 37oC before addition of cells. Following a 48 hr incubation, cells were lysed and Luc activity determined using a microtiter plate luminometer and BriteLite Plus Reagent (Perkin Elmer). Neutralization titers were the sample dilution at which relative luminescence units (RLU) were reduced by 50% compared to RLU in virus control wells after subtraction of background RLU in cell control wells.

### Antibody Dependent Cell Mediated Virus Inhibition (ADCVI)

ADCVI assay was performed as previously described with some modifications(Kannanganat et al., 2016). Briefly, on day 1, CEM-NK^r^ cells were spinoculated at 1500xg for 3 hrs with tier2 Clade C SHIV1157ipd3N4 virus at 31 TCID50/mL. On day 2 of the assay, cryopreserved human donor PBMCs were thawed, washed, and counted. Cells were added to a V-bottom plate (Corning incorporated, Corning, NY) at a concentration of 1×10^5^ cells/well and allowed to rest overnight. On day 3, purified rabbit IgG (250µg/ml) was incubated for 2 hrs with 1×10^4^ infected CEM-NK^r^ cells that had been washed 3 times to remove unbound virus. After 2 hrs of incubation, purified rabbit IgG and infected CEM-NK^r^ cells were added to PBMCs. PGT121 (catalog no. 12343; NIH AIDS Reagent Program) and EM4C04 (anti-influenza HA antibody) served as positive and negative controls respectively. Five days post incubation, cells were washed 2 times and fresh media was added to all wells. On day 7 post incubation, plates were spun at 1500 rpm for 5 mins and the supernatant was harvested and frozen until p27 Gap ELISAs could be performed.

### SIV Gag p27 ELISA

High binding ELISA plates (Thermo Scientific, catalog no 44-2404-21) were coated at 0.5µg/mL with goat anti-mouse IgG2b (Southern Biotech, catalog no 1090-05) overnight at 4°C. Next day, plates were washed 6x with PBST (PBS + 0.05%Tween-20) and blocked with 1% BSA in PBST for 30 minutes at RT. Anti-p27 2F12 (catalog no. 2343; NIH AIDS Reagent Program) antibody at 0.5 µg/mL was added to ELISA plates and incubated for 1 hr at 37°C. Plates were again washed 6x with PBST. Supernatant from ADCVI assays which were placed in the 4°C the night before, were treated with TritonX (Sigma) to a make a 0.5% Triton X solution to inactivate virus particles and release Gag. Gag supernatant was diluted 3-fold and added to ELISA plates and allowed to incubate for 1 ½ hour at 37°C. Plates were washed and biotinylated anti-SIV IgG diluted 1:1000 and added to each well. Plates were again incubated for 1 hr at 37°C. After 6x wash, neutralite-avidin peroxidase (N-HRP) (Southern Biotech, Birmingham, AL) was diluted 1:4000 and added to each plate for 30 minutes at RT in the dark. After incubation, bound IgG was detected using tetramethylbenzidine substrate (KPL, Gaithersburg, MD). The reaction was stopped by adding 100 µl 2N H_2_SO_4_. The readings were recorded at 450nm.

### Hydrogen/Deuterium Exchange Mass Spectrometry

5 µgs (52 pmol) per timepoint of each protein (1086.C UFO-v2-RQH^173^ and UFO-v2-RQY^173^ and BG505.664) were incubated in deuterated buffer (20mM PBS, 85% D2O, pH 7.5) for 3s, 1min, 30min, and 20hrs at room temperature. The reaction was stopped via diluting 1:1 in ice-cold quench buffer (200 mM tris(2-chlorethyl) phosphate (TCEP), 8 M urea, 0.2% formic acid) to a final pH of 2.5 and flash frozen in liquid nitrogen followed by storage in −80°C prior to analysis. Zero time points and fully deuterated samples were prepared as described by Verkerke et al. 2016(Verkerke et al., 2016). Online pepsin digestion was performed and analyzed by LC-MS-IMS utilizing a Waters Synapt G2-Si Q-TOF mass spectrometer as described by Verkerke et al. 2016(Verkerke et al., 2016). Deuterium uptake analysis was performed with HD-Examiner (Sierra Analytics) followed by HX-Express v2(Guttman et al., 2014; Weis et al., 2006). The percent exchange was normalized to the fully deuterated samples. Internal exchange standards (Pro-Pro-Pro-Ile [PPPI] and Pro-Pro-Pro-Phe [PPPF]) were included in each reaction to control for variations in ambient temperature during the labeling reactions.

### Dynamic Light Scattering (DLS)

Dynamic light scattering (DLS) measurements were performed on a Dynapro Nanostar (Wyatt Technologies). Trimer samples were diluted to 1 mg/ml in PBS and centrifuged at 15,000xg for 20 min prior to loading of 10μl into a low-volume quartz cuvette. The mean estimated hydrodynamic radius, and polydispersity were generated from 30 acquisitions of 5 s at 20°C. For time dependent DLS experiments, the samples were diluted to 1 mg/mL in PBS and allowed to sit at room temperature (23 °C) for 4 hours prior to centrifugation and measurement by DLS.

## Statistical Analyses

All statistical analyses were performed using GraphPad Prism v8 software. Data represent mean or mean ± SD. Statistical significance between groups was performed by Mann-Whitney’s test and two-tailed Student’s t test (small sample size, comparing time dependent affinity values between two C.1086 proteins for specific mAb tested). *p < 0.05, **p< 0.01, ***p<0.001, ****p<0.0001. p-values color coded by immunization group correspond to comparison with WT.

## Acknowledgements

We thank Dr. Lynn Morris for providing CAP228-16H mAb; Dr. Jens Wrammert for anti-influenza EM4C04 Ab, Dr. Cynthia Derdeyn for pCDNA3.1 C.1086 K160N plasmid and Dr. Pam Kozlowski for anti-SIV IgG protein, and Dr. John P. Moore for pPPI4 BG505 SOSIP T332N plasmid. We also thank Dr. Genevieve Giny Fouda for providing PGT151, CAP256-VRC26.08, PGDM1400 bnAbs. We are extremely thankful to Dr. Du Yuhong, Emory University, GA, USA for letting use the Octet RED384 instrument for the BLI experiments. We thank Dr. Raghavan Varadarajan, IISc, Bangalore, India 560012 and Dr. Robert Sonowal, Emory University, GA, USA for their valuable suggestions.

## Funding

This work was supported in part by National Institutes of Health Grants U19 AI109633 to R.R.A. R01 AI140868 to K.K.L, and NCRR/NIH base grant P51 OD011132 to YNPRC. Work related to negative stain electron-microscopy done at Dr. Andrew B. Ward’s lab was supported by the Bill and Melinda Gates Foundation through the Collaboration for AIDS Vaccine Discovery (CAVD) grant OPP1115782 to A.B.W.

### Author Contributions

R.R.A. and A.S.^1^ designed the study and wrote the manuscript. A.S.^1^ performed experiments and analyzed data. E.A.H did HIC purification, H/D exchange, DLS experiments (supervised by K.L.), C.L. conducted neutralization assay (supervised by D.C.M.), T.T.S did ADCVI, X.S did BAMA assay (supervised by G.D.T.), G.O and W-H.L. did NS-EM (supervised by A.W.). A.S^1^., N.C. and A.S^2^ purified proteins and plasmids. A.S^1^ performed ELISA, BLI binding experiments. D.J.I. provided ISCOM. ^1^Anusmita Sahoo, ^2^Ayalnesh Shiferaw.

## Data and materials Availability

All unique mutant plasmids generated in this study may be requested from the authors with a completed Materials Transfer Agreement. The study did not generate/analyze any dataset/code.

## Supplementary Information

The article contains following supplementary materials.

### Supplementary Figure and Table Legends

**Figure S1.**
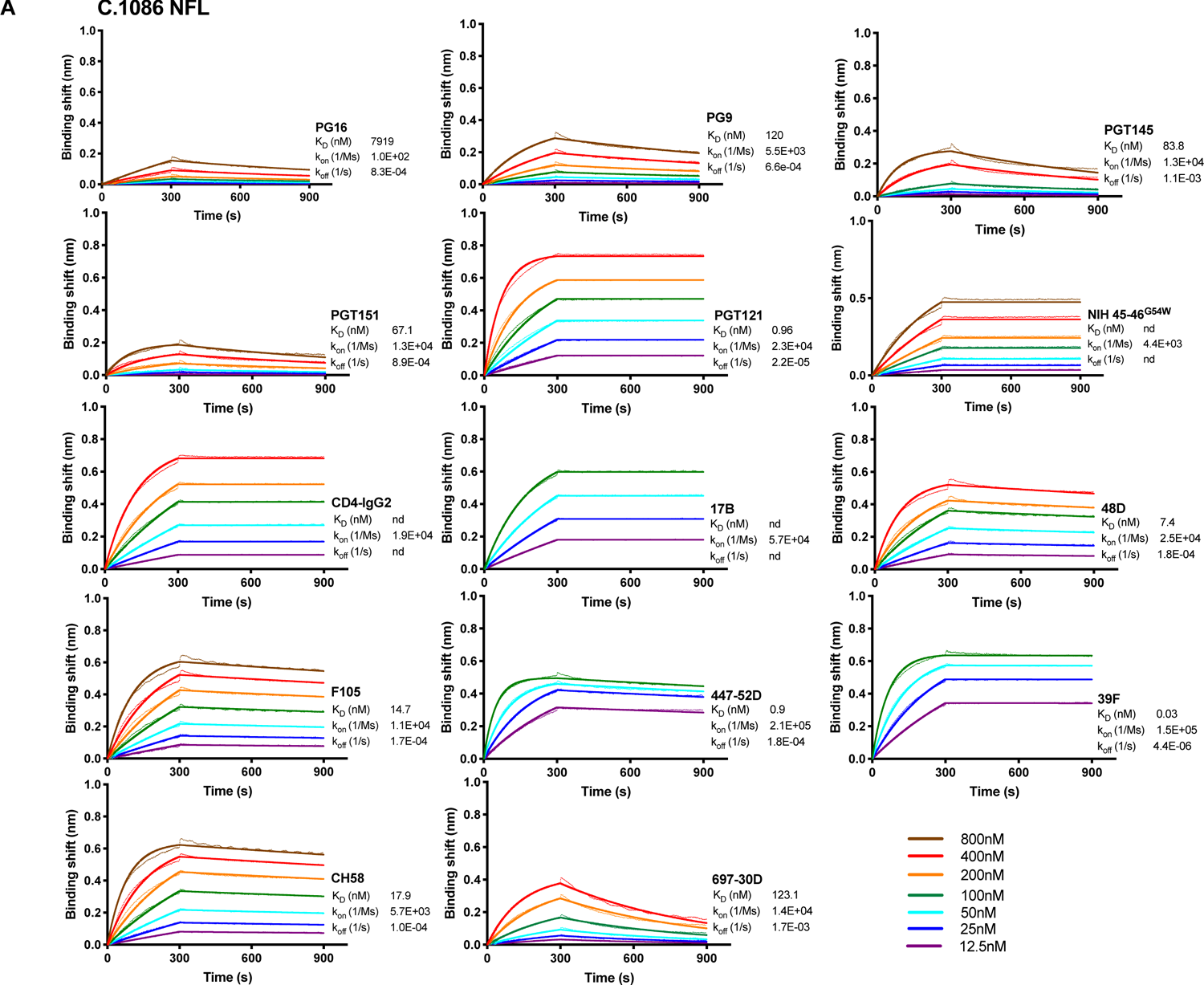
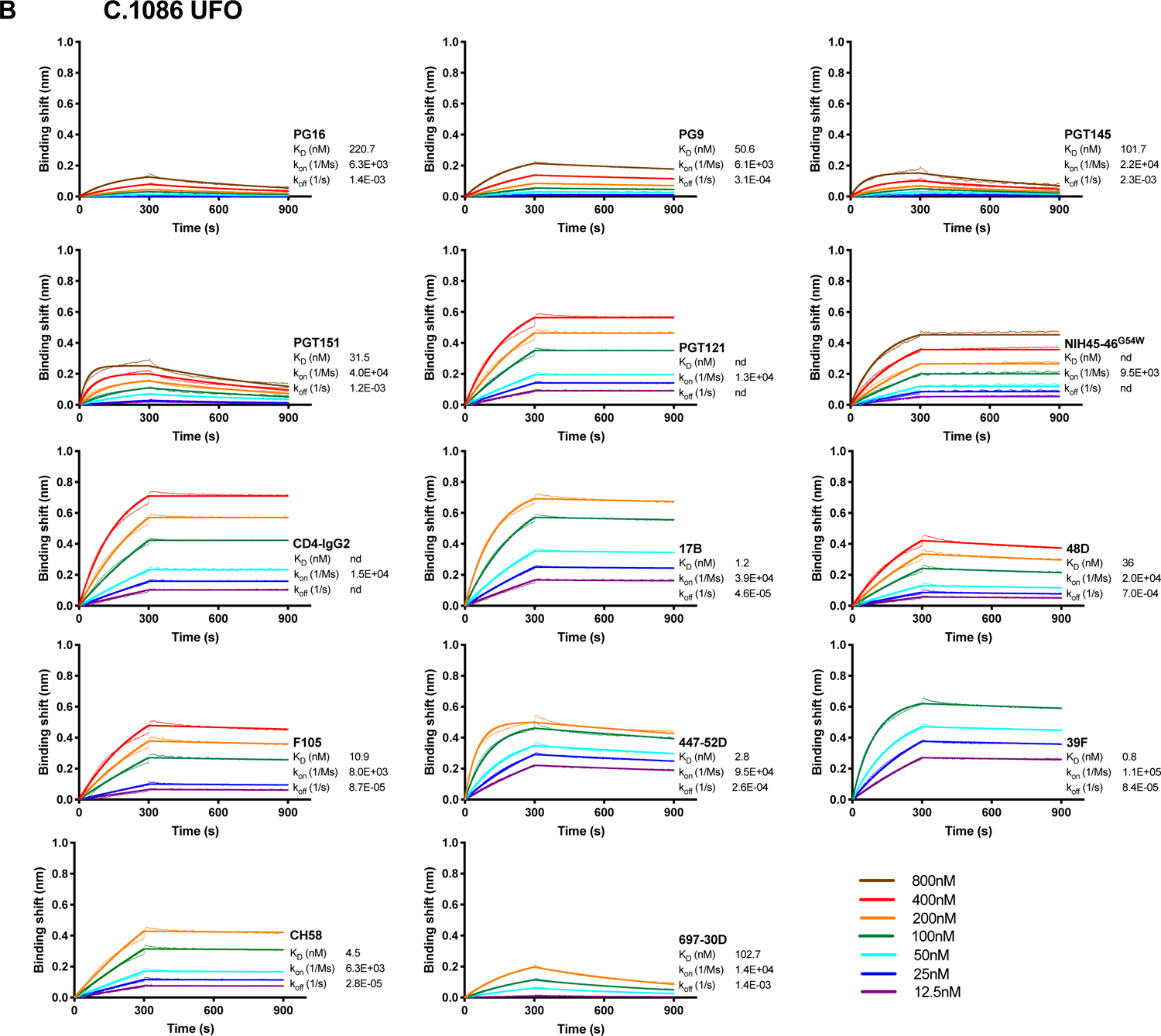
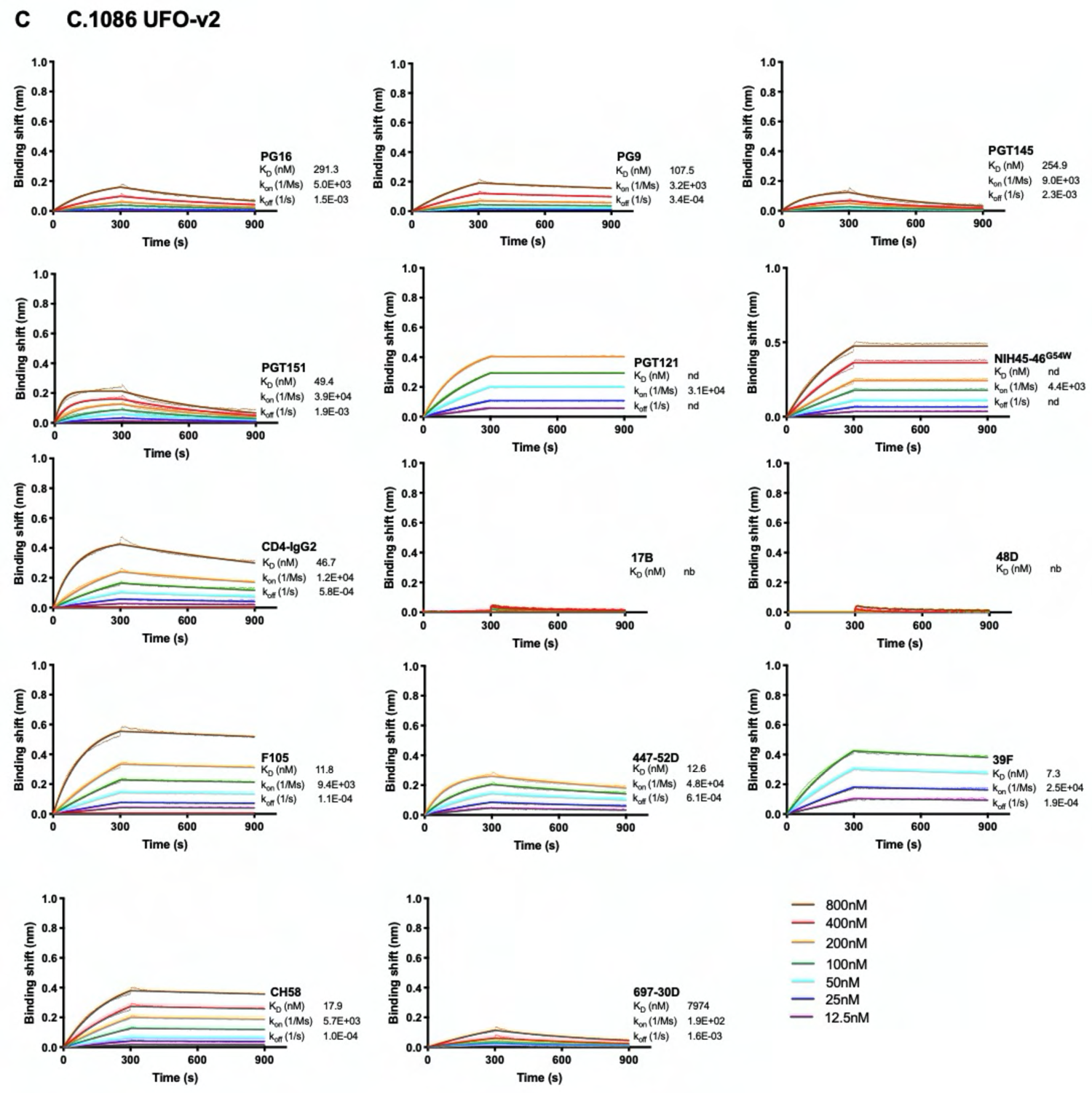
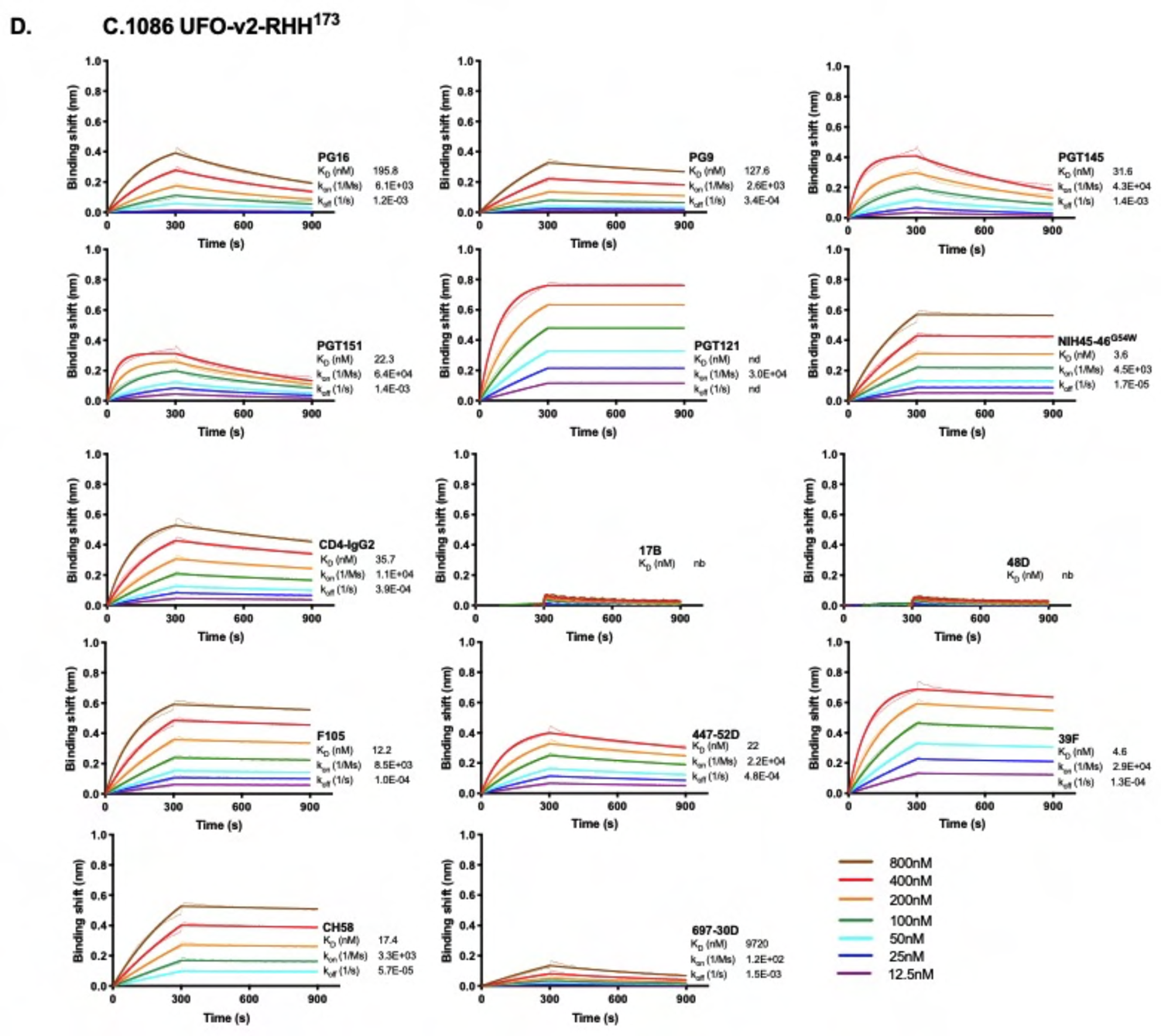
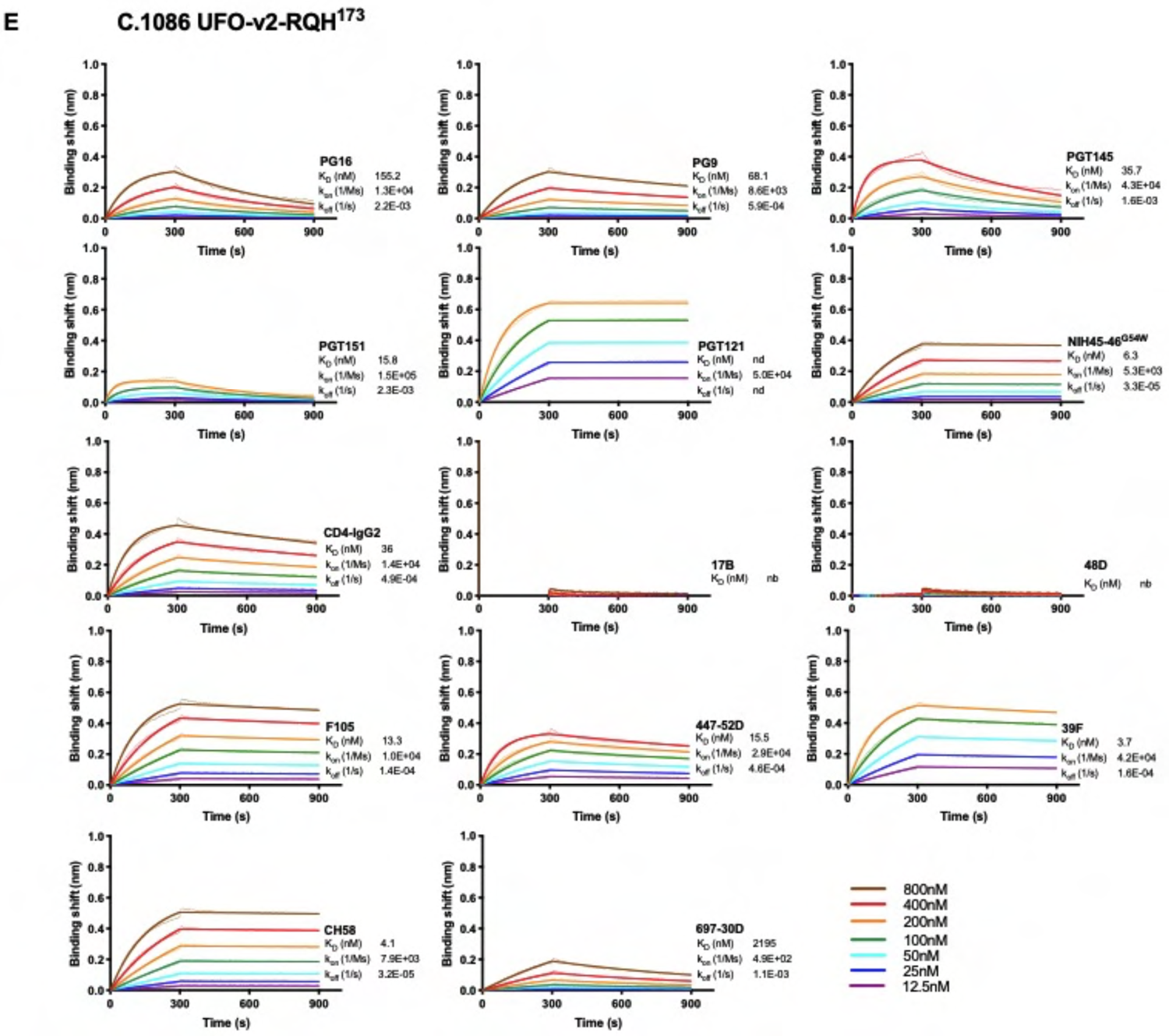
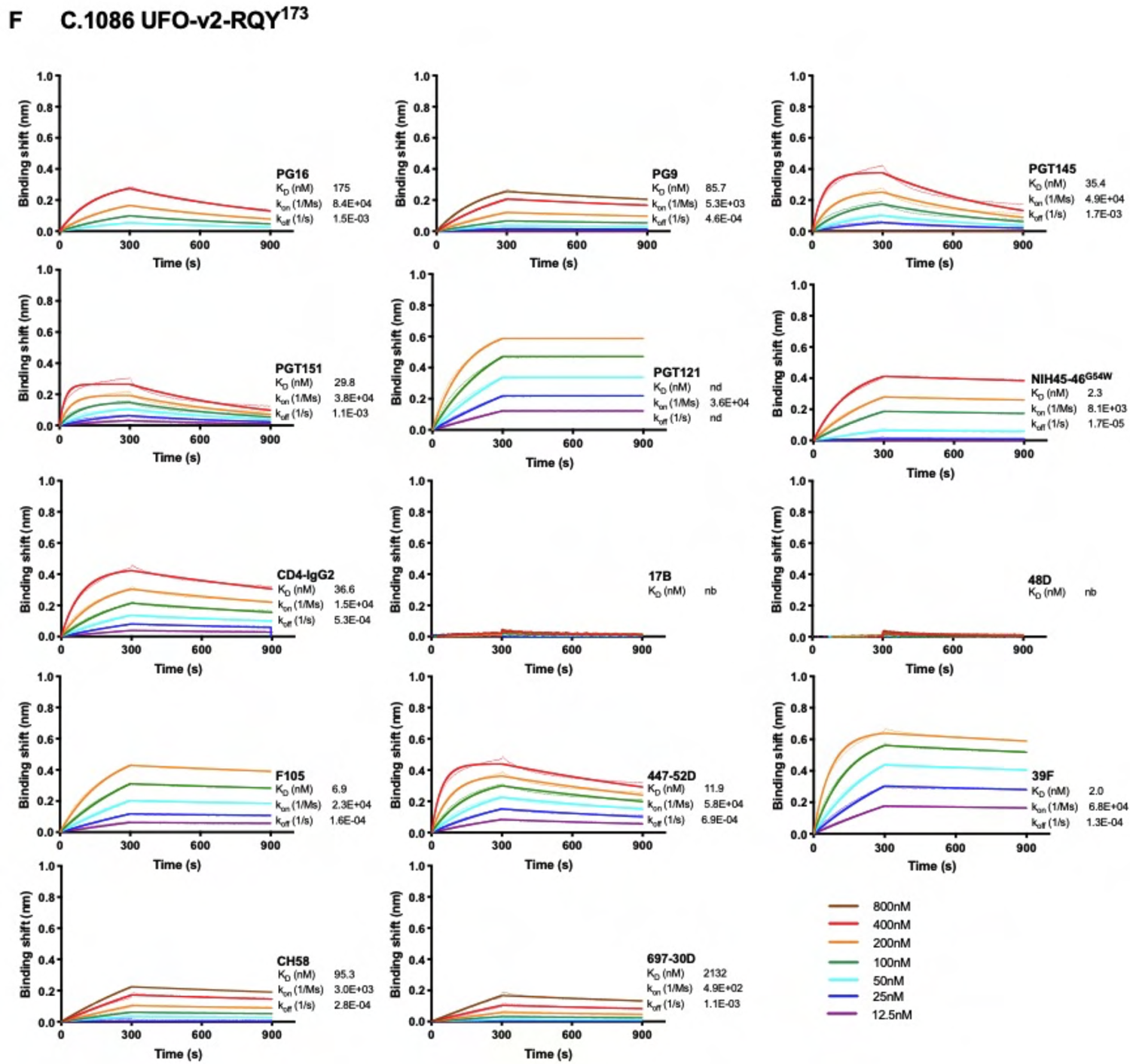
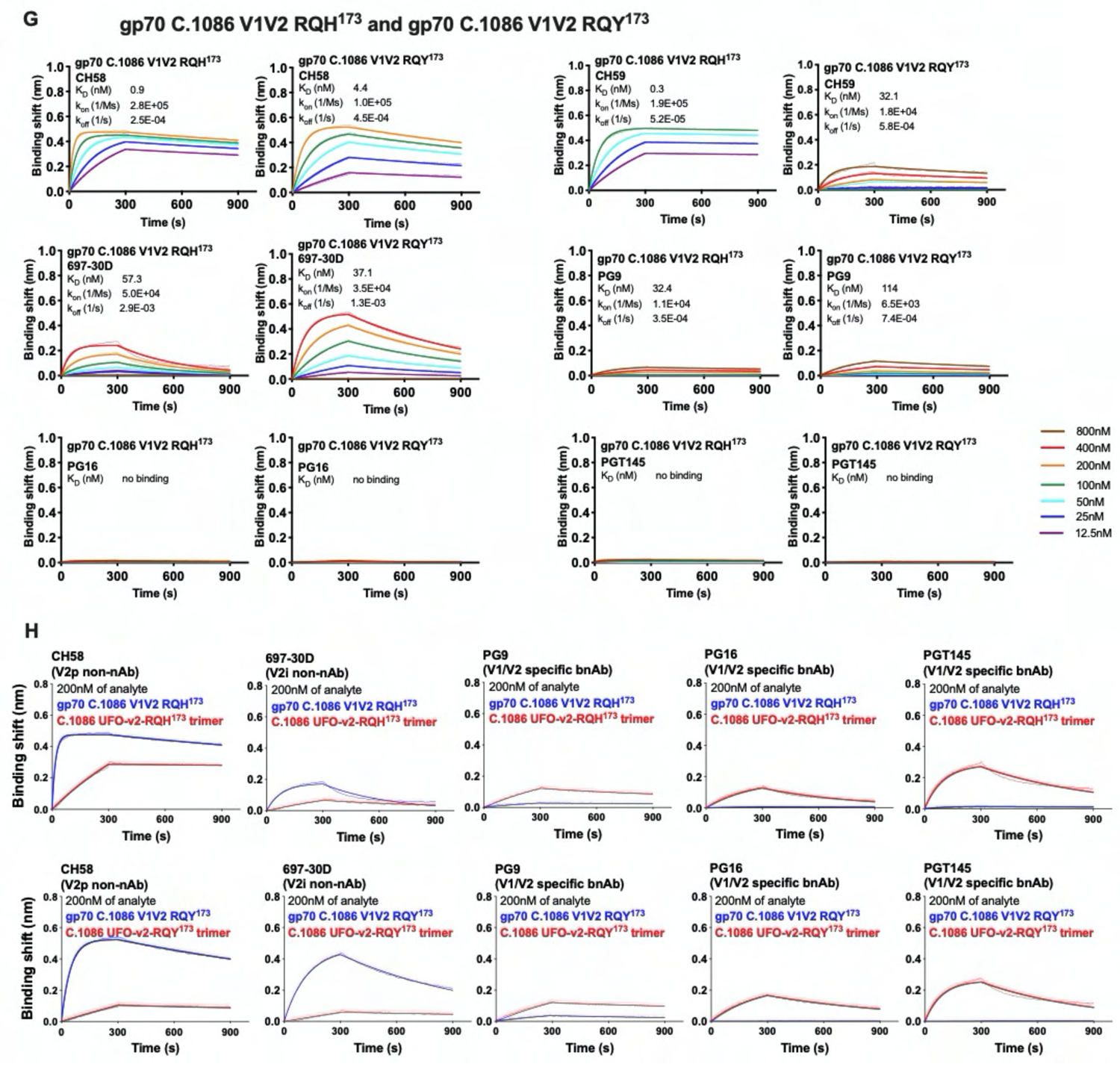
Binding sensograms of purified C.1086 gp140 and gp70-V1V2 variants against envelope specific mAbs monitored by Bio-layer Interferometry (BLI), related to manuscript. Figures 1 and 2. Binding sensograms of purified C.1086 (**A)** NFL, (**B)** UFO, (**C)** UFO-v2, (**D)** UFOv2-RHH^173^, (**E)** UFO-v2-RQH^173^, (**F)** UFO-v2-RQY^173^, (**G)** gp70 C.1086 V1V2 RQ(H/Y)^173^ proteins by BLI. The proteins (analytes) were serially diluted (800-12.5nM) and binding kinetics were measured against desired envelope specific mAb (5µg/ml) immobilized on anti-human Fc Biosensors for 300s association and 600s dissociation on Octet Red384 platform (ForteBio). Each panel shows raw traces (dotted line) monitored at different concentrations of the analyte and traces fitted (solid line) to a global 1:1 binding model. Kinetic parameters indicated were obtained from globally fitting responses from the concentrations which give the best fit to a 1:1 binding model using ForteBio Data Analysis v9 software. Responses from these concentrations shown. (**H)** Overlay of binding sensograms monitored for 200nM, Top: gp70 C.1086 V1V2 RQH^173^ and C.1086 UFO-v2-RQH^173^, Bottom: gp70 C.1086 V1V2 RQY^173^ and C.1086 UFO-v2-RQY^173^ against env specific mAbs. All gp140 variants tested had K160N/V295N/N334S mutations in the protein backbone.

**Figure S2.**
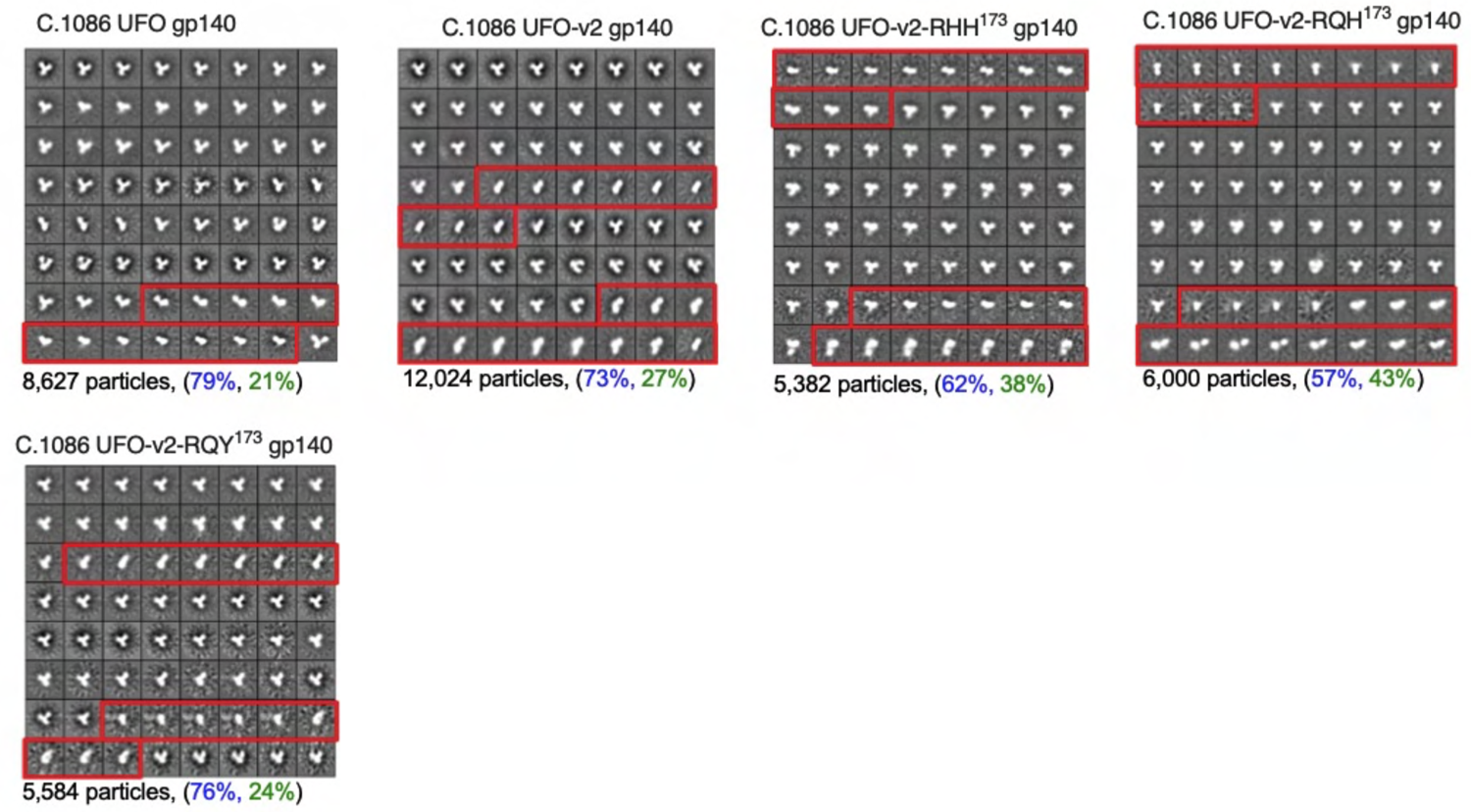
2D Class averages of purified C.1086 gp140 variants by negative-stain electron microscopy, related to manuscript. Figures 1 and 2. 2D class averages of C.1086 protein variants UFO, UFO-v2, UFO-v2-RHH173, UFO-v2-RQH173, UFO-v2-RQY173 with classes representing non-native trimer features (dissociated protomers or malformed trimers) boxed in red. Total particles imaged, %native-like (blue) and non-native like malformed trimers (green) indicated. The proteins used for the negative stain electron microscopy were expressed from 293F cells, purified using *Galanthus nivalis* lectin-based affinity chromatography, followed by isolation of trimer protein by size exclusion chromatography. **Figure**

**S3.**
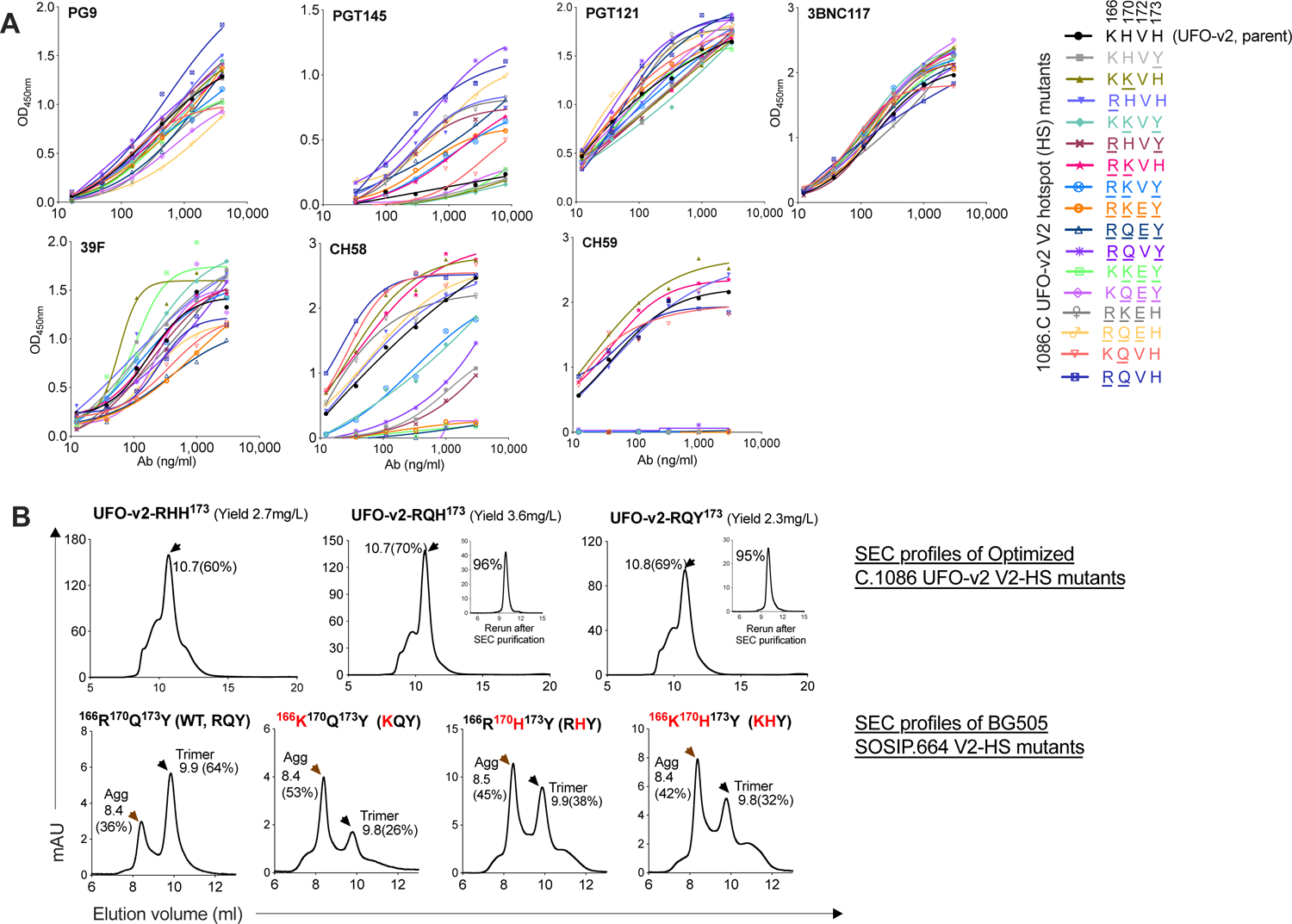
Screening of C.1086 UFO-v2 V2-HS mutants and influence of the V2-HS modifications on protein trimeric proportion, related to manuscript Figure 2. (A) Screening of UFO-v2 V2 hotspot (HS) mutants by ELISA for improved binding to multiple bnAbs, specifically PGT145. Supernatants from C.1086 UFO-v2-V2 hotspot (V2-HS) mutants (listed in right) expressed from 293T cells were assayed for binding to envelope specific mAbs by ELISA. Representative binding traces of two independent experiments. (B) Size exclusion chromatograms of *Galanthus nivalis* lectin affinity purified; Top, C.1086 UFO-v2 V2-HS and Bottom, BG505 SOSIP V2-HS mutants. Inset, Re-run SEC traces (UFO-v2-RQH^173^ and UFO-v2-RQY^173^) of pooled purified trimeric fractions collected after SEC purification step. Elution volume and proportion (%, AUC) of trimeric peak (black arrow) and “Agg” Aggregate/Oligomeric indicated.

**Figure S4.**
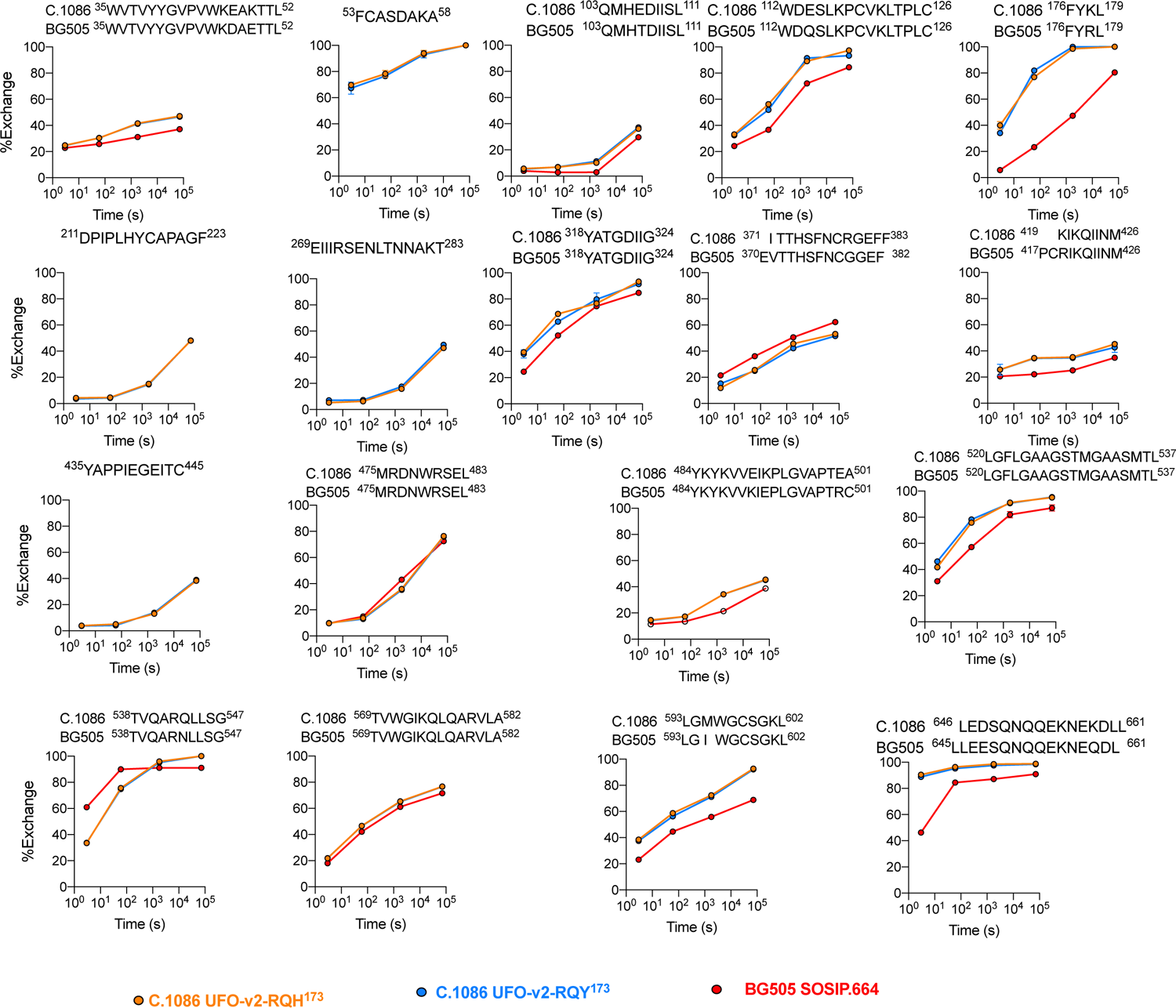
HDX-MS analyses of C.1086 UFO-v2-RQH^173^, UFO-v2-RQY^173^ and homologous peptic peptides in BG505 SOSIP.664 trimers, related to manuscript. Figure 3. Exchange kinetics of individual peptides of purified C.1086 UFO-v2-RQH^173^, UFO-v2-RQY^173^ and BG505 SOSIP.664 proteins at 0s, 1min, 60min, 20hrs. Exchange profiles of BG505 peptides homologous to C.1086 were overlayed onto the profiles of C.1086 peptides for the same region. The percent exchange was normalized to the zero-time point and fully deuterated samples. Mean ± SD values of duplicate experiments plotted.

**Figure S5.**
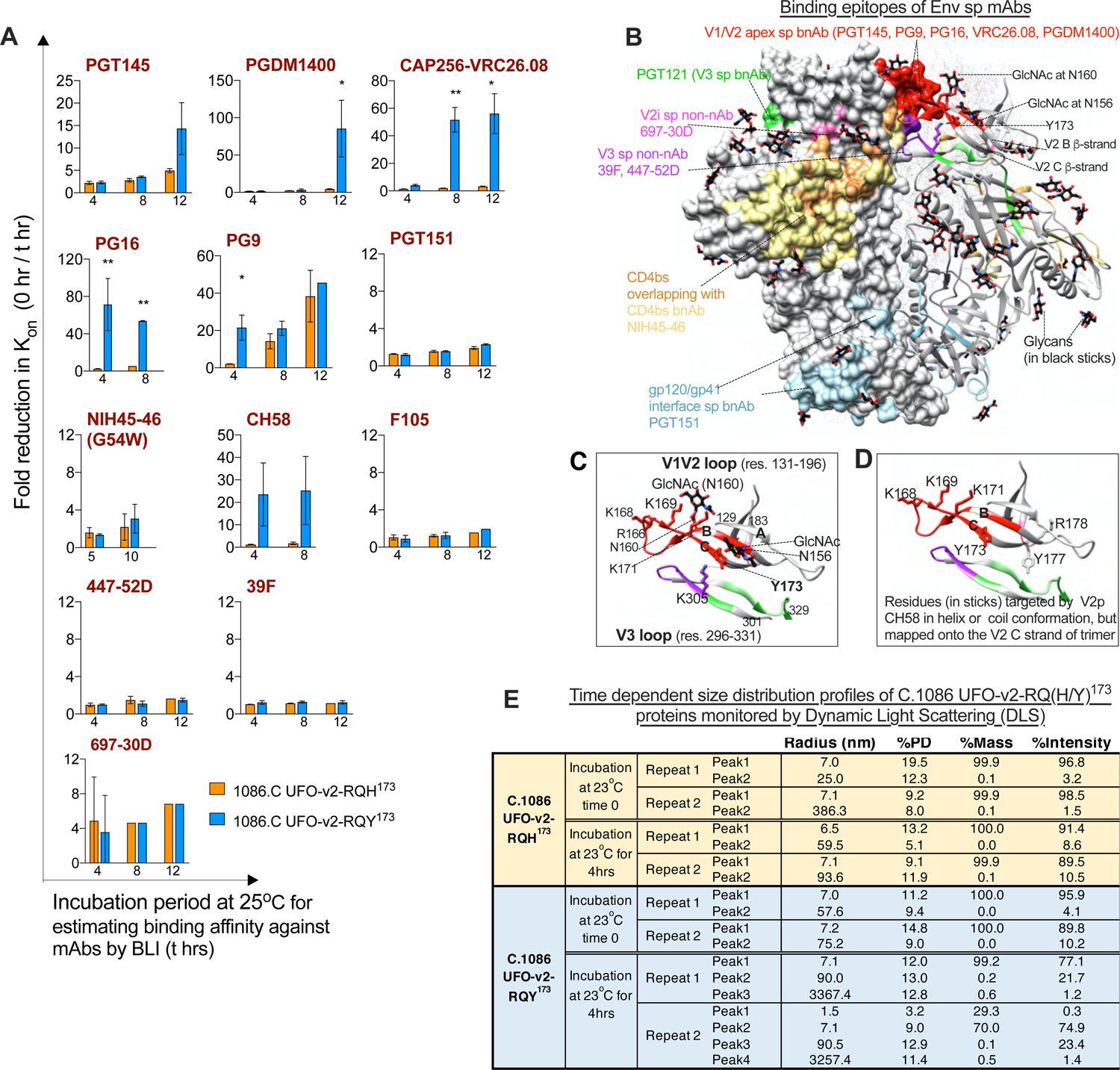
Time dependent structural differences monitored for C.1086 UFO-v2-RQ(H/Y)^173^ variants by BLI and DLS, *in-vitro,* related to manuscript Figure 3. **(A)** Time dependent fold reduction in association rate (K_on_ 0hr / K_on_ t hr) of UFO-v2-RQH^173^ and UFO-v2-RQY^173^ variants against envelope specific mAbs. Binding kinetics of purified C.1086 UFO-v2-RQH^173^ and UFO-v2-RQY^173^ proteins against envelope specific mAbs was estimated by BLI after the proteins (analytes) were incubated at 25°C for t = 0, 4, 8, 12hrs or 0, 5, 10hrs. Association (300s) and dissociation (600s) traces were globally fit to a 1:1 binding model using ForteBio Data Analysis v9 software. Bars show Mean, error bars SD of at-least three independent experiments. Student’s t test for statistical comparisons, *p < 0.05, **p< 0.01, ***p<0.001, ****p<0.0001. (**B)** Glycans (represented as black sticks) and epitopes targeted by various envelope specific mAbs (color coded), V1V2 B, C β-strands, Y173, GlcNAc at N156, N160 highlighted on the structure of unliganded BG505 SOSIP.DS trimer (pdb id 4ZMJ (Kwon et al., 2015)). Protomers in solid, ribbon and dotted surface representation. (**C-D**) Enlarged view of V1V2 and V3 loop. Refer panel (**B)** for epitope specific color codes. V2 B-C strands harboring V1V2 apex sp bnAbs epitope, red. (**C)** Y173 (V2 C strand) and its adjacent glycan at N156 position (V2 B strand), cationic residues and N160 glycan in the V2 B-C strand shown. (**D)** Residues targeted by V2p CH58 non-nAb in helix/coil conformation are mapped onto the V2 C strand of the trimer. (**B-D)** were made using UCSF Chimera v1.14 (Pettersen et al., 2004). (**E)** Time dependent size distribution profiles of purified C.1086 UFO-v2-RQ(H/Y)^173^ proteins monitored by Dynamic Light Scattering (DLS). For readings at 4hrs, the samples were diluted to 1 mg/mL in PBS and allowed to sit at room temperature (23°C) for 4 hours prior to centrifugation and measurement. The mean estimated hydrodynamic radius, and polydispersity (PD) were generated from 30 acquisitions of 5 s at 20°C.

**Figure S6.**
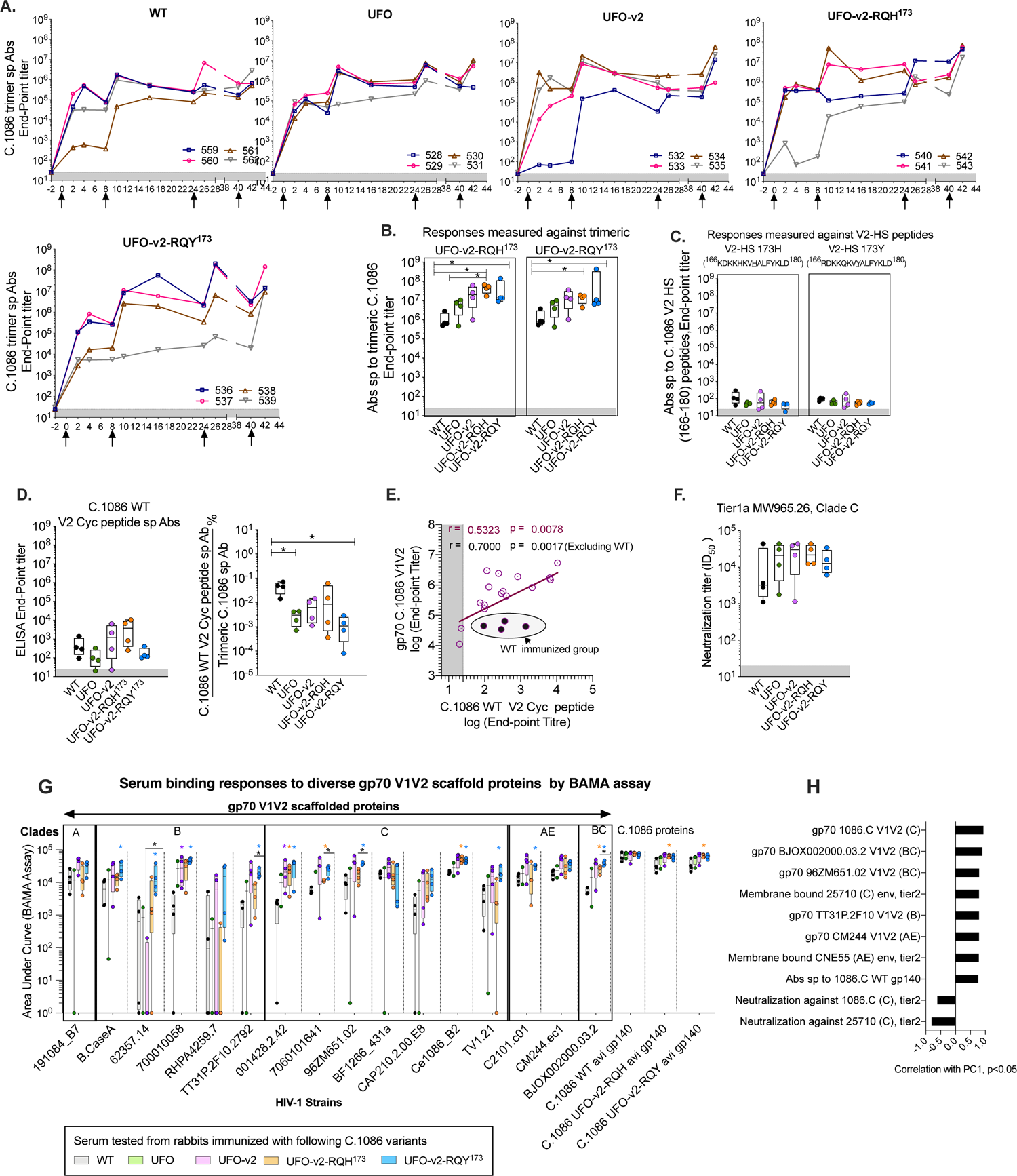
Characterization of antibody responses elicited by C.1086 variants in rabbits, related to manuscript. Figures 4 and 5**. (A)** Serum from rabbits immunized with C.1086 constructs WT, UFO, UFO-v2, UFO-v2-RQH**^173^**, UFO-v2-RQY**^173^** were assayed longitudinally (0 – 42 weeks) to monitor C.1086 UFO-v2-RQH**^173^** trimer specific binding antibodies by ELISA. Mean end-point titers calculated by ELISA from two independent experiments were plotted. Immunization time points indicated by black arrows. Responses of serum (2 weeks after final protein boost) against (**B)** UFO-v2-RQH^173^ and UFO-v2-RQY^173^ by ELISA, (**C)** C.1086 V2 hotspot (HS) peptides V2-HS 173H (166KDKKHKV<u>H</u>ALFYKLD1^80^) and V2-HS 173Y (^166^RDKKQKV<u>Y</u>ALFYKLD^180^), (**D)** C.1086 WT V2 cyclic peptides (left) and response normalized to trimer specific responses (End point titer measured against C.1086 WT V2 cyc peptide / End point titer measured against C.1086 UFO-v2-RQH^173^ (right). (**E)** Spearman’s correlation between responses against gp70 C.1086 V1V2 (End-point titer by ELISA) and C.1086 WT V2 cyc peptide (End-point titer by ELISA). Correlations w/o responses from WT immunized group shown. (**F)** Neutralization ID_50_ (dilution of the serum required to neutralize 50% infection) of serum (2 weeks after final protein boost) monitored against Tier1a Clade C MW965.26 pseudotyped virus by standard TZM-bl assay (Montefiori, 2009). (**G)** Binding Antibody Multiplex Assay (BAMA) analyses (binding AUC) (Zolla-Pazner et al., 2014) of serially diluted serum from WT, UFO, UFO-v2-RQH^173^ and UFO-v2-RQY^173^ immunized rabbits to V1V2 from diverse 16 cross-clade HIV-1 strains (grafted onto gp70 scaffold protein, referred as V1V2-scaffold) and C.1086 WT, UFO-v2-RQH^173^, UFO-v2-RQY^173^ C terminal avi tagged proteins. (**H)** Top most significant (p < 0.05) variables of the PCA analyses done using serum characterization data of different immunized groups. Correlation of variables with PC1, p <0.05 have been plotted. PCA was carried out using the R. (**A-F)** Shaded area represents background signal based on responses to pre-bleed serum. (**G, H)** Serum collected two weeks after final protein boost used for analyses. (**B-D, F, G)** Box and whiskers plots where box extends from 25th to 75th percentile, median indicated by line, minimum and maximum values indicated by whiskers. Statistical comparisons between groups by Mann-Whitney test (*p < 0.05, **p< 0.01, ***p<0.001, ****p<0.0001). (**G)** p-values immunized group color coded correspond to comparison with WT. All values plotted are the average of at-least two independent experiments.

**Figure S7.**
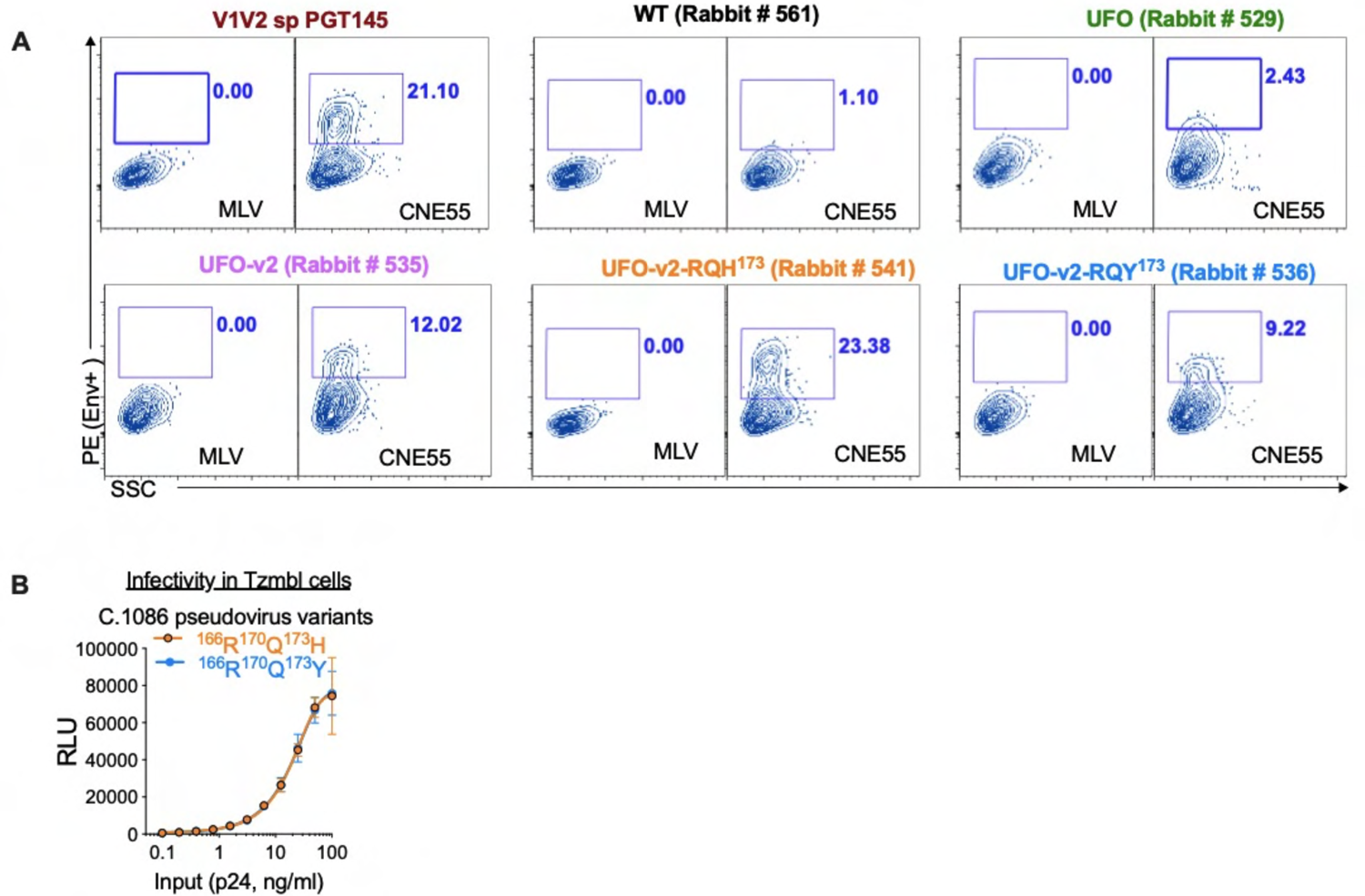
Representative flow plots showing binding of purified serum IgGs to 293T cells expressing membrane anchored gp160 and comparing infectivity of TZM-bl cells by C.1086 RQ(HY)^173^ pseudoviruses, related to manuscript. Figure 4**. (A)** Representative flow plots showing binding of purified serum IgGs (1µg/ml) to live transiently transfected 293T cells expressing membrane anchored gp160. Shown here are representative binding flow plots to tier2 CNE55 env^+^ 293T cells. Binding signal of rabbit IgG to MLV transfected cells used as -ve control to define binding signal gate. Binding of cells expressing envelopes to env sp bnAbs (here PGT145) used as +ve control. (**B)** Infectivity of TZM-bl cells by serially diluted C.1086 RQH^173^ and RQY^173^ pseudoviruses (input virus equalized based on p24 concentration), measured by luminescence (RLU, mean ± SD of two independent experiments, each done in duplicates).

**Table S1.** **Binding kinetics of C.1086 gp140 variants estimated by Bio-Layer Interferometry, related to manuscript Figures 1 and 2**. Binding affinities of C.1086 gp140 proteins (NFL, UFO, UFO-v2, UFO-v2-RQH^173^, UFO-v2-RHH^173^, UFO-v2-RQY^173^) were monitored against envelope specific mAbs by BioLayer Interferometry using Octet Red384 platform. Env specific mAbs were immobilized on anti-human Fc biosensor. The proteins were used as analyte (800-12.5nM serially diluted). The data was globally fit to a 1:1 binding model. Mean ± SD values calculated from more than two independent experiments reported. ^a^nd: No dissociation observed, hence K_D_ could not be calculated. All variants tested had K160N/V295N/N334S mutations in the protein backbone.

| Epitope specificity | Env sp mAb | C.1086<br>K160N/V295N/N334S<br>gp140 | K<sub>D</sub> (nM) | k<sub>on</sub>(1/Ms) | k<sub>dis</sub>(1/s) |
| --- | --- | --- | --- | --- | --- |
| V1/V2 apex<br>sp bnAb | PG16 | NFL | 11044±4414 | 1.1±0.03E+02 | 1.2±0.5E-03 |
|  |  | UFO | 252±44 | 6.6±0.5E+03 | 1.7±0.4E-03 |
|  |  | UFO-v2 | 275±23 | 5.5±0.8E+03 | 1.5±0.11E-03 |
|  |  | UFO-v2-RHH<sup>173</sup> | 215±27 | 5.8±0.4E+03 | 1.3±0.09E-03 |
|  |  | UFO-v2-RQH<sup>173</sup> | 153±17 | 1.3±0.1E+04 | 2±0.2E-03 |
|  |  | UFO-v2-RQY<sup>173</sup> | 217±84 | 7.3±3.8E+03 | 1.3±0.08E-03 |
|  | PG9 | NFL | 167±67 | 4.4±1.6E+03 | 6.9±0.4E-04 |
|  |  | UFO | 74±33 | 5.7±0.6E+03 | 4.1E-04 |
|  |  | UFO-v2 | 108 | 3.2E+03 | 3.4E-04 |
|  |  | UFO-v2-RHH<sup>173</sup> | 100±39 | 4±1.9E+03 | 3.6±0.3E-04 |
|  |  | UFO-v2-RQH<sup>173</sup> | 70±10 | 8.3±0.8E+03 | 5.8±0.3E-04 |
|  |  | UFO-v2-RQY<sup>173</sup> | 91±7 | 4.3±0.9E+03 | 3.9±0.7E-04 |
|  | PGT145 | NFL | 136±56 | 1.3±0.2E+04 | 1.7±0.6E-03 |
|  |  | UFO | 111±53 | 1.6±0.9E+04 | 1.6±0.6E-03 |
|  |  | UFO-v2 | 197±82 | 1.2±0.5E+04 | 2.2±0.08E-03 |
|  |  | UFO-v2-RHH<sup>173</sup> | 34±4 | 3.7±0.8E+04 | 1.3±0.1E-03 |
|  |  | UFO-v2-RQH<sup>173</sup> | 37±6 | 4.3±1.1E+04 | 1.6±0.2E-03 |
|  |  | UFO-v2-RQY<sup>173</sup> | 34±3 | 5.1±0.3E+04 | 1.7±0.03E-03 |
| gp120/gp41<br>bnAb | PGT151 | NFL | 97±38 | 1.2±0.2E+04 | 1.1±0.2E-03 |
|  |  | UFO | 35±5 | 3±0.9E+04 | 1±0.2E-03 |
|  |  | UFO-v2 | 54±18 | 3.8±1.7E+04 | 1.9±0.3E-03 |
|  |  | UFO-v2-RHH<sup>173</sup> | 24±6 | 4.6±2E+04 | 1.1±0.3E-03 |
|  |  | UFO-v2-RQH<sup>173</sup> | 20±2 | 7.5±0.8E+04 | 1.50E-03 |
|  |  | UFO-v2-RQY<sup>173</sup> | 34±6 | 3.7±0.3E+04 | 1.2±0.1E-03 |
| V3 glycan sp<br>bnAn | PGT121 | NFL | 2±0.3 | 1.4±0.5E+04 | 2±0.3E-05 |
|  |  | UFO | <sup>a</sup>nd | 1.4±0.5E+04 | <sup>a</sup>nd |
|  |  | UFO-v2 | <sup>a</sup>nd | 2.2±1.3E+04 | <sup>a</sup>nd |
|  |  | UFO-v2-RHH<sup>173</sup> | <sup>a</sup>nd | 2.8±0.3E+04 | <sup>a</sup>nd |
|  |  | UFO-v2-RQH<sup>173</sup> | <sup>a</sup>nd | 1.9±0.6E+04 | <sup>a</sup>nd |
|  |  | UFO-v2-RQY<sup>173</sup> | <sup>a</sup>nd | 1.5±0.5E+04 | <sup>a</sup>nd |
| CD4bs bnAb | NIH45-46(G54W) | NFL | 2±1 | 7±2.2E+03 | 1.5±1.1E-05 |
|  |  | UFO | <sup>a</sup>nd | 1.1±0.2E+03 | <sup>a</sup>nd |
|  |  | UFO-v2 | <sup>a</sup>nd | 3.1±2.1E+03 | <sup>a</sup>nd |
|  |  | UFO-v2-RHH<sup>173</sup> | 4 | 4.5E+03 | 1.7E-05 |
|  |  | UFO-v2-RQH<sup>173</sup> | 4±1 | 1.3±0.1E+04 | 5.4±0.8E-05 |
|  |  | UFO-v2-RQY<sup>173</sup> | 2±1 | 8.1±2.3E+03 | 1.7±0.5E-05 |
| CD4 mimic | CD4-IgG2 | NFL | <sup>a</sup>nd | 1.4±0.2E+04 | <sup>a</sup>nd |
|  |  | UFO | <sup>a</sup>nd | 1.2±0.2E+04 | <sup>a</sup>nd |
|  |  | UFO-v2 | 43±5 | 1.2±0.04E+04 | 5.3±0.7E-04 |
|  |  | UFO-v2-RHH<sup>173</sup> | 37±2 | 1.1±0.1E+04 | 4±0.2E-04 |
|  |  | UFO-v2-RQH<sup>173</sup> | 39±4 | 1.5±0.2E+04 | 5.9±1.4E-04 |
|  |  | UFO-v2-RQY<sup>173</sup> | 43±9 | 1.2±0.4E+04 | 5±0.5E-04 |
| CD4i non-nAb | 17B | NFL | 3 | 3.4±0.4E+04 | 1.1E-04 |
|  |  | UFO | 3±2 | 2.8±0.5E+04 | 7.9±4.3E-05 |
|  |  | UFO-v2 |  | no apparent binding |  |
|  |  | UFO-v2-RHH<sup>173</sup> |  | no apparent binding |  |
|  |  | UFO-v2-RQH<sup>173</sup> |  | no apparent binding |  |
|  |  | UFO-v2-RQY<sup>173</sup> |  | no apparent binding |  |
|  | 48D | NFL | 15±7 | 3.7±2.6E+04 | 6.3±6.4E-04 |
|  |  | UFO | 27±13 | 1.6±0.5E+04 | 4.7±3.3E-04 |
|  |  | UFO-v2 |  | no apparent binding |  |
|  |  | UFO-v2-RHH<sup>173</sup> |  | no apparent binding |  |
|  |  | UFO-v2-RQH<sup>173</sup> |  | no apparent binding |  |
|  |  | UFO-v2-RQY<sup>173</sup> |  | no apparent binding |  |
| CD4bs non-nAb | F105 | NFL | 16±2 | 1.1±0.1E+04 | 1.7±0.07E-04 |
|  |  | UFO | 7±4 | 1.1±0.3E+04 | 7.1±3.8E-05 |
|  |  | UFO-v2 | 15±5 | 9±0.7E+03 | 1.3±0.3E-04 |
|  |  | UFO-v2-RHH<sup>173</sup> | 16±5 | 8.6±0.2E+03 | 1.3±0.4E-04 |
|  |  | UFO-v2-RQH<sup>173</sup> | 10±5 | 1.3±0.4E+04 | 1.2±0.2E-04 |
|  |  | UFO-v2-RQY<sup>173</sup> | 7±1 | 1.8±0.7E+04 | 1.3±0.4E-04 |
| V3 non-nAb | 447-52D | NFL | 2±1 | 1.4±0.6E+05 | 2.2±0.6E-04 |
|  |  | UFO | 2±1 | 7±1.8E+04 | 1.8±0.8E-04 |
|  |  | UFO-v2 | 15±3 | 3.4±2.1E+04 | 4.7±2E-04 |
|  |  | UFO-v2-RHH<sup>173</sup> | 18±5 | 2.2±0.09E+04 | 4.1±1E-04 |
|  |  | UFO-v2-RQH<sup>173</sup> | 15±1 | 3.6±0.9E+04 | 5.2±0.8E-04 |
|  |  | UFO-v2-RQY<sup>173</sup> | 10±2 | 5.8±0.6E+04 | 5.5±1.9E-04 |
|  | 39F | NFL | 1±1 | 1.1±0.3E+05 | 9.3±12E-05 |
|  |  | UFO | 1±0.4 | 7.4±4.4E+04 | 4.8±5E-05 |
|  |  | UFO-v2 | 7±0.05 | 2.2±0.4E+04 | 1.6±0.3E-04 |
|  |  | UFO-v2-RHH<sup>173</sup> | 6±1 | 2.6±0.3E+04 | 1.5±0.2E-04 |
|  |  | UFO-v2-RQH<sup>173</sup> | 4±0.2 | 4.8±0.8E+04 | 1.7±0.2E-04 |
|  |  | UFO-v2-RQY<sup>173</sup> | 3±1 | 5.7±1.6E+04 | 1.5±0.3E-04 |
| V2p non-nAb | CH58 | NFL | 13±3 | 1.4±0.2E+04 | 1.8±0.1E-04 |
|  |  | UFO | 4±2 | 1.3±0.3E+04 | 6±4.4E-05 |
|  |  | UFO-v2 | 20±3 | 3.7±2.9E+03 | 7±4.7E-05 |
|  |  | UFO-v2-RHH<sup>173</sup> | 21±6 | 4.6±1.9E+03 | 1±0.7E-04 |
|  |  | UFO-v2-RQH<sup>173</sup> | 5±0.3 | 9.5±0.3E+03 | 4.9±0.5E-05 |
|  |  | UFO-v2-RQY<sup>173</sup> | 94±3 | 3.3±0.5E+03 | 3.1±0.3E-04 |
| V2i non-nAb | 697-30D | NFL | 151±39 | 1.2±0.3E+04 | 1.7±0.04E-03 |
|  |  | UFO | 107±6 | 1±0.5E+04 | 1.1±0.5E-03 |
|  |  | UFO-v2 | 7483±694 | 1.7±0.3E+02 | 1.3±0.4E-03 |
|  |  | UFO-v2-RHH<sup>173</sup> | 7213±3546 | 8.8±4.3E+01 | 7.1±6.2E-04 |
|  |  | UFO-v2-RQH<sup>173</sup> | 2387±1211 | 5.5±4E+02 | 9.7±3.5E-04 |
|  |  | UFO-v2-RQY<sup>173</sup> | 2403±1089 | 4.3±4.9E+02 | 7.1±4.9E-04 |

<sup>a</sup>nd: No dissociation observed, hence K<sub>D</sub> could not be calculated.

**Table S2.** **Time dependent binding kinetics assessment of C.1086 UFO-v2-RQ(H/Y)^173^ variants against envelope specific mAbs by BLI at 25°C, related to manuscript** **Figure 3** **and Figure S5.** Binding kinetics parameters of C.1086 UFO-v2-RQH^173^ and UFO-v2-RQY^173^ proteins measured against envelope specific mAbs by BLI after the proteins (analytes) were incubated at 25°C for t = 0, 4, 8, 12hrs or 0, 5, 10hrs. Association (300s) and dissociation (600s) traces were globally fit to a 1:1 binding model using ForteBio Data Analysis v9 software. Mean ± SD values of at-least three independent experiments reported. ^a^ntd: binding affinity not able to determine due to low binding signal to reliably globally fit the raw data, ^b^nd: no dissociation observed, hence K_D_ could not be calculated, ^c^ND: not determined.

| Epitope specificity | Envelope sp mAbs | Time of Incubation at 25°C (t, hr) | C.1086 UFO-v2-RQH<sup>173</sup> |  |  | C.1086 UFO-v2-RQY<sup>173</sup> |  |  |
| --- | --- | --- | --- | --- | --- | --- | --- | --- |
|  |  |  | K<sub>D</sub> (nM) | k<sub>on</sub>(1/Ms) | k<sub>dis</sub>(1/s) | K<sub>D</sub> (nM) | k<sub>on</sub>(1/Ms) | k<sub>dis</sub>(1/s) |
| V1/V2 apex sp bnAb | PG16 | 0 | 153±17 | 1.3±0.1E+04 | 2±0.2E-03 | 217±84 | 7.3±3.8E+03 | 1.3±0.08E-03 |
|  |  | 4 | 332±26 | 5.3±0.5E+03 | 1.8±0.1E-03 | 18309±6469 | 7.7±3.4E+01 | 1.3±0.1E-03 |
|  |  | 8 | 719 | 2.50E+03 | 1.70E-03 | 11138±746 | 9.4±1.4E+01 | 1±0.08E-03 |
|  |  | 12 |  | <sup>a</sup>ntd |  | 17521 | 5.8E+01 | 1.00E-03 |
|  | PG9 | 0 | 70±10 | 8.3±0.8E+03 | 5.8±0.3E-04 | 91±7 | 4.3±0.9E+03 | 3.9±0.7E-04 |
|  |  | 4 | 153±34 | 3.9±0.7E+03 | 5.8±0.2E-04 | 1847±615 | 1.8±0.4E+02 | 3.2±0.3E-04 |
|  |  | 8 | 1071±443 | 5.9±2.3E+02 | 5.8±0.2E-04 | 1730±344 | 1.8±0.2E+02 | 3.1±0.3E-04 |
|  |  | 12 | 2550±829 | 2.4±1E+02 | 5.6±0.9E-04 | 2006 | 8.6E+01 | 1.80E-04 |
|  | PGT145 | 0 | 35±1 | 4.5±0.3E+04 | 1.6±0.1E-03 | 43±4 | 3.1±0.3E+04 | 1.4±0.08E-03 |
|  |  | 4 | 67±7 | 2.3±0.5E+04 | 1.5±0.2E-03 | 91±17 | 1.4±0.04E+04 | 1.3±0.2E-03 |
|  |  | 8 | 90±7 | 1.7±0.2E+04 | 1.6±0.2E-03 | 152±45 | 8.3±0.8E+03 | 1.3±0.3E-03 |
|  |  | 12 | 154±12 | 10±1.4E+03 | 1.5±0.1E-03 | 567±80 | 2.2±0.8E+02 | 1.2±0.3E-03 |
|  | PGDM1400 | 0 | 46±4 | 1.5±0.2E+04 | 6.5±0.3E-04 | 67±15 | 1±0.3E+04 | 6.7±1.1E-04 |
|  |  | 4 | 76±23 | 9.7±2E+04 | 7.1±0.5E-04 | 117±33 | 6.2±0.09E+03 | 7.2±1.3E-04 |
|  |  | 8 | 98±3 | 6.5±0.8E+03 | 6.3±0.6E-04 | 212±9 | 2.8±0.3E+03 | 5.9±0.4E-04 |
|  |  | 12 | 208±64 | 3.5±0.9E+03 | 7.1±0.5E-04 | 5623±661 | 1.1±0.1E+02 | 6.4±1.3E-04 |
|  | CAP256-VRC26.08 | 0 | 27±9 | 1±0.4E+04 | 2.5±0.3E-04 | 35±2 | 6.7±1.2E+03 | 2.4±0.3E-04 |
|  |  | 4 | 35±8 | 8.8±1.1E+03 | 3±0.3E-04 | 165±12 | 1.1±1E+03 | 2.7±0.2E-04 |
|  |  | 8 | 53±12 | 6.5±0.7E+03 | 3.4±0.4E-04 | 2364±176 | 1.3E+02 | 3.1±0.2E-04 |
|  |  | 12 | 91±31 | 3.7±0.9E+03 | 3.4±0.4E-04 | 2732±800 | 1.1±0.3E+02 | 3.2±1.7E-04 |
| gp120/gp41 interface sp bnAb | PGT151 | 0 | 20±2 | 7.5±0.8E+04 | 1.50E-03 | 34±6 | 3.7±0.3E+04 | 1.2±0.1E-03 |
|  |  | 4 | 28±4 | 5.9±0.4E+04 | 1.6±0.1E-03 | 48±4 | 3.1±0.08E+04 | 1.5±0.2E-03 |
|  |  | 8 | 34±5 | 4.8±0.2E+04 | 1.7±0.2E-03 | 69±0.7 | 2.3±0.06E+04 | 1.6±0.1E-03 |
|  |  | 12 | 42±6 | 3.9±0.9E+04 | 1.7±0.2E-03 | 103±20 | 1.6±0.06E+04 | 1.6±0.3E-03 |
| V3 glycan sp bnAn | PGT121 | 0 | <sup>b</sup>nd | 3±0.2E+04 | <sup>b</sup>nd | <sup>b</sup>nd | 2±0.8E+04 | <sup>b</sup>nd |
|  |  | 4 | <sup>b</sup>nd | 2.3±0.5E+04 | <sup>b</sup>nd | <sup>b</sup>nd | 1.4±0.1E+04 | <sup>b</sup>nd |
|  |  | 8 | <sup>b</sup>nd | 1.9±0.6E+04 | <sup>b</sup>nd | <sup>b</sup>nd | 1.5±0.5E+04 | <sup>b</sup>nd |
|  |  | 12 | <sup>b</sup>nd | 2.20E+04 | <sup>b</sup>nd | <sup>b</sup>nd | 1.1E+04 | <sup>b</sup>nd |
|  |  |  | K<sub>D</sub> (nM) | k<sub>on</sub>(1/Ms) | k<sub>dis</sub>(1/s) | K<sub>D</sub> (nM) | k<sub>on</sub>(1/Ms) | k<sub>dis</sub>(1/s) |
| CD4bs bnAb | NIH45-46(G54W) | 0 | 4±1 | 1.3±0.1E+04 | 5.4±0.8E-05 | 2±1 | 8.1±2.3E+03 | 1.7±0.5E-05 |
|  |  | 5 | 5±0.6 | 8.5±2E+03 | 4.5±0.6E-05 | 2±1 | 6±2.1E+03 | 1.5±1.3E-05 |
|  |  | 10 | 6±0.7 | 7.2±4E+03 | 4.7±3E-05 | 4 | 3.2±2.3E+03 | 1.90E-05 |
|  |  | 15 | 11 | 7.50E+03 | 8.10E-05 | ° ND |  |  |
| CD4bs non-nAb | F105 | 0 | 11 | 1.5±0.2E+04 | 1.8±0.02E-04 | 16±6 | 1.2±0.7E+04 | 1.6±0.4E-04 |
|  |  | 4 | 12±2.3 | 1.5±0.2E+04 | 1.8±0.1E-04 | 17±12 | 1.6±1.4E+04 | 1.8±0.4E-04 |
|  |  | 8 | 13±3 | 1.3±0.04E+04 | 1.6±0.5E-04 | 21±14 | 1±0.8E+04 | 1.6±0.3E-04 |
|  |  | 12 | 19 | 1.10E+04 | 2.00E-04 | 42 | 3.5E+04 | 1.50E-04 |
| V3 non-nAb | 447-52D | 0 | 12±3 | 4.8±1E+04 | 5.5±0.2E-04 | 22±8 | 2.9±2E+04 | 5.3±1.5E-04 |
|  |  | 4 | 10±1 | 4.9±0.1E+04 | 5±0.6E-04 | 23±10 | 3±2.2E+04 | 5.4±1E-04 |
|  |  | 8 | 16±6 | 3.2±0.2E+04 | 5.1±1.6E-04 | 27±16 | 3.1±2.9E+04 | 5.3±0.8E-04 |
|  |  | 12 | 21 | 3.30E+04 | 6.80E-04 | 40±0.9 | 1.2±0.3E+04 | 4.8±1.5E-04 |
|  | 39F | 0 | 3±0.6 | 4.2±0.4E+04 | 1.2±0.E-04 | 7±4 | 2.7±2E+04 | 1.4±.2E-04 |
|  |  | 4 | 4±0.3 | 4±0.3E+04 | 1.6±0.003E-04 | 10.4±3 | 2.1±1.2E+04 | 2±0.7E-04 |
|  |  | 8 | 5±0.1 | 3.6±0.4E+04 | 1.9±0.2E-04 | 12±6 | 9.7±4.2E+03 | 2.1±0.7E-04 |
|  |  | 12 | 6 | 3.30E+04 | 2.10E-04 | 15±7 | 1.3±0.3E+04 | 1.9±0.5E-04 |
| V2p non-nAb | CH58 | 0 | 5±0.3 | 9.5±0.3E+03 | 4.9±0.5E-05 | 94±3 | 3.3±0.5E+03 | 3.1±0.3E-04 |
|  |  | 4 | 7±2 | 8.3±1.7E+03 | 5.2±0.5E-05 | 1421±379 | 1.6±0.8E+02 | 2±0.3E-04 |
|  |  | 8 | 13±1.6 | 5.6±1.8E+03 | 7.4±3.3E-05 | 1250±27 | 1.5±0.7E+02 | 1.9±0.9E-04 |
|  |  | 12 | 14 | 6.80E+03 | 9.70E-05 | °ntd |  |  |
| V2i non-nAb | 697-30D | 0 | 2387±1211 | 5.5±4E+02 | 9.7±3.5E-04 | 2403±1089 | 4.3±4.9E+02 | 7.1±4.9E-04 |
|  |  | 4 | 7960±774 | 1.2±0.03E+02 | 9.5±0.7E-04 | 3753±4118 | 1.3±0.09E+02 | 4.5±4.7E-04 |
|  |  | 8 | 3624 | 2.10E+02 | 7.70E-04 | 3624 | 2.1E+02 | 7.70E-04 |
|  |  | 12 | 3964 | 1.50E+02 | 5.70E-04 | 3964 | 1.5E+02 | 5.70E-04 |

°ntd: binding affinity not able to determine due to low binding signal to reliably globally fit the raw data, °nd: no dissociation observed, hence K<sub>D</sub> could not be calculated, °ND: not determined.

**Table S3.**
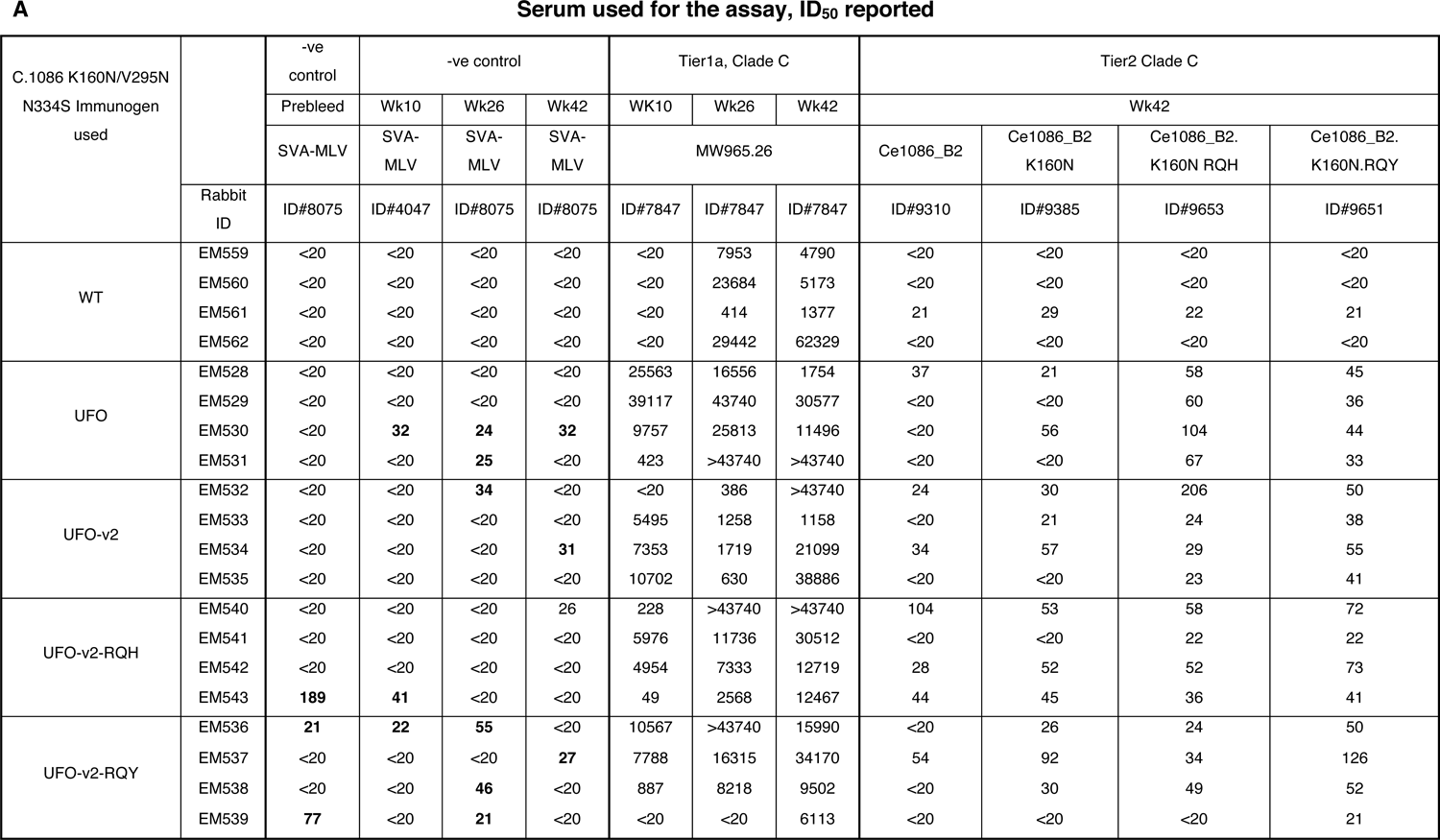
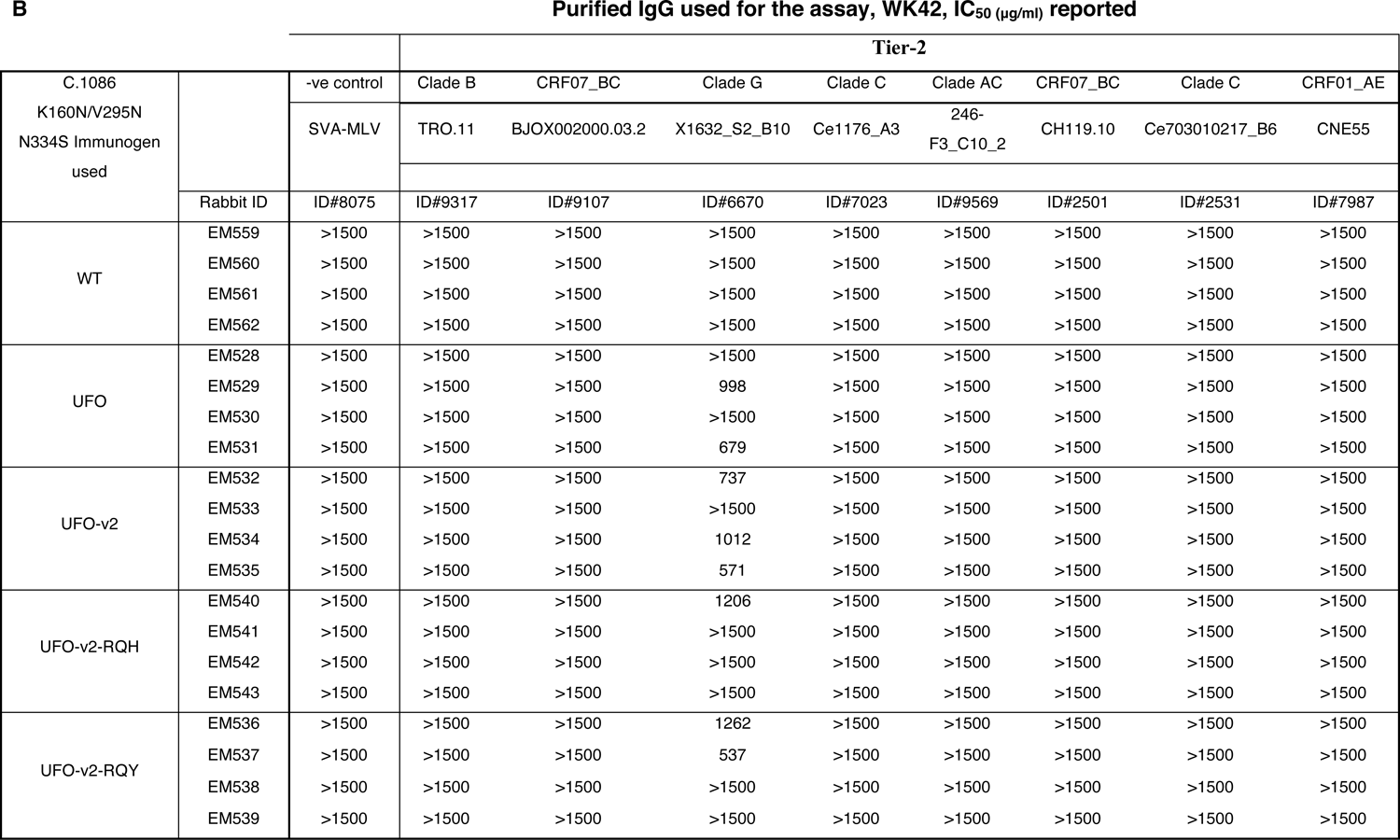
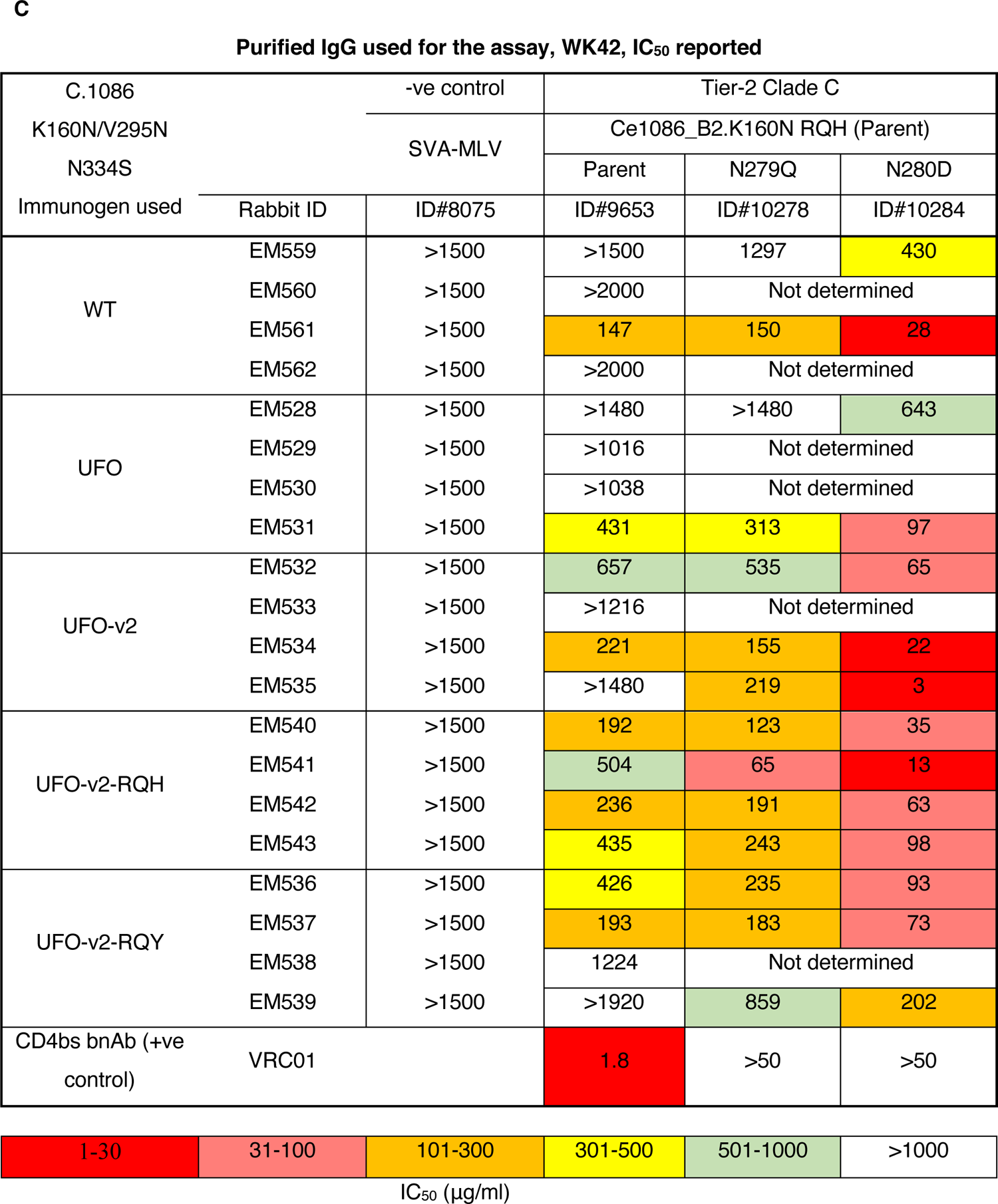
**Neutralization sensitivity of serum from C.1086 immunized rabbits, related to manuscript** **Figure 4. (A)** Neutralization titers (ID_50_, serum dilution to acheive 50% inhibition of viral infection by pseudotyped TZM-bl assay) using serum collected 2 weeks after 2^nd^ (wk 10), 3^rd^ (wk 26) and 4^rth^ (wk 42) immunizations against Tier-1 Clade C MW965.26, and wk42 responses against C.1086 pseudotyped virus variants. Serum showing non-specific activity against negative control MLV (background cut-off < 20) have been highlighted in bold. **(B)** Neutralization titers (IC_50_ µg/ml, concentration of IgG required to achieve 50% neutralization of the pseudovirus) of purified IgG from serum collected on wk 42 i.e., 2 weeks after last immunization against global HV-1 Tier-2 panel. (**C)** Neutralization IC_50_ (concentration of IgG required for 50% neutralization of the virus) values of rabbit purified IgG (week 42 i.e. 2 weeks after the final protein boost) from various immunized groups against C.1086 K160N RQH (parent) and CD4bs mutants N279Q, N280D monitored by standard TZM-bl assay (Montefiori, 2009).

## References

1. Aldon, Y., McKay, P.F., Allen, J., Ozorowski, G., Felfodine Levai, R., Tolazzi, M., Rogers, P., He, L., de Val, N., Fabian, K., et al. (2018). Rational Design of DNA-Expressed Stabilized Native-Like HIV-1 Envelope Trimers. Cell Rep 24, 3324–3338 e3325.

2. Arunachalam, P.S., Charles, T.P., Joag, V., Bollimpelli, V.S., Scott, M.K.D., Wimmers, F., Burton, S.L., Labranche, C.C., Petitdemange, C., Gangadhara, S., et al. (2020). T cell-inducing vaccine durably prevents mucosal SHIV infection even with lower neutralizing antibody titers. Nat Med 26, 932–940.

3. Bale, S., Martine, A., Wilson, R., Behrens, A.J., Le Fourn, V., de Val, N., Sharma, S.K., Tran, K., Torres, J.L., Girod, P.A., et al. (2018). Cleavage-Independent HIV-1 Trimers From CHO Cell Lines Elicit Robust Autologous Tier 2 Neutralizing Antibodies. Front Immunol 9, 1116.

4. Barouch, D.H., Liu, J., Li, H., Maxfield, L.F., Abbink, P., Lynch, D.M., Iampietro, M.J., SanMiguel, A., Seaman, M.S., Ferrari, G., et al. (2012). Vaccine protection against acquisition of neutralization-resistant SIV challenges in rhesus monkeys. Nature 482, 89–93.

5. Berendam, S.J., Styles, T.M., Morgan-Asiedu, P.K., Tenney, D., Kumar, A., Obregon-Perko, V., Bar, K.J., Saunders, K.O., Santra, S., De Paris, K., et al. (2021). Systematic Assessment of Antiviral Potency, Breadth, and Synergy of Triple Broadly Neutralizing Antibody Combinations against Simian-Human Immunodeficiency Viruses. J Virol 95.

6. Binley, J.M., Lybarger, E.A., Crooks, E.T., Seaman, M.S., Gray, E., Davis, K.L., Decker, J.M., Wycuff, D., Harris, L., Hawkins, N., et al. (2008). Profiling the specificity of neutralizing antibodies in a large panel of plasmas from patients chronically infected with human immunodeficiency virus type 1 subtypes B and C. J Virol 82, 11651–11668.

7. Bontempo, A., Garcia, M.M., Rivera, N., and Cayabyab, M.J. (2020). A Systematic Approach to HIV-1 Vaccine Immunogen Selection. AIDS Res Hum Retroviruses 36, 762–770.

8. Bricault, C.A., Yusim, K., Seaman, M.S., Yoon, H., Theiler, J., Giorgi, E.E., Wagh, K., Theiler, M., Hraber, P., Macke, J.P., et al. (2019). HIV-1 Neutralizing Antibody Signatures and Application to Epitope-Targeted Vaccine Design. Cell Host Microbe 25, 59–72 e58.

9. Brouwer, P.J.M., Antanasijevic, A., Berndsen, Z., Yasmeen, A., Fiala, B., Bijl, T.P.L., Bontjer, I., Bale, J.B., Sheffler, W., Allen, J.D., et al. (2019). Enhancing and shaping the immunogenicity of native-like HIV-1 envelope trimers with a two-component protein nanoparticle. Nat Commun 10, 4272.

10. Burton, S., Spicer, L.M., Charles, T.P., Gangadhara, S., Reddy, P.B.J., Styles, T.M., Velu, V., Kasturi, S.P., Legere, T., Hunter, E., et al. (2019). Clade C HIV-1 Envelope Vaccination Regimens Differ in Their Ability To Elicit Antibodies with Moderate Neutralization Breadth against Genetically Diverse Tier 2 HIV-1 Envelope Variants. Journal of virology 93, e01846–01818.

11. Cimbro, R., Gallant, T.R., Dolan, M.A., Guzzo, C., Zhang, P., Lin, Y., Miao, H., Van Ryk, D., Arthos, J., Gorshkova, I., et al. (2014). Tyrosine sulfation in the second variable loop (V2) of HIV-1 gp120 stabilizes V2-V3 interaction and modulates neutralization sensitivity. Proc Natl Acad Sci U S A 111, 3152–3157.

12. de Taeye, S.W., Ozorowski, G., Torrents de la Pena, A., Guttman, M., Julien, J.P., van den Kerkhof, T.L., Burger, J.A., Pritchard, L.K., Pugach, P., Yasmeen, A., et al. (2015). Immunogenicity of Stabilized HIV-1 Envelope Trimers with Reduced Exposure of Non-neutralizing Epitopes. Cell 163, 1702–1715.

13. deCamp, A., Hraber, P., Bailer, R.T., Seaman, M.S., Ochsenbauer, C., Kappes, J., Gottardo, R., Edlefsen, P., Self, S., Tang, H., et al. (2014). Global panel of HIV-1 Env reference strains for standardized assessments of vaccine-elicited neutralizing antibodies. J Virol 88, 2489–2507.

14. Doria-Rose, N.A., Klein, R.M., Manion, M.M., O’Dell, S., Phogat, A., Chakrabarti, B., Hallahan, C.W., Migueles, S.A., Wrammert, J., Ahmed, R., et al. (2009). Frequency and phenotype of human immunodeficiency virus envelope-specific B cells from patients with broadly cross-neutralizing antibodies. J Virol 83, 188–199.

15. Dubrovskaya, V., Guenaga, J., de Val, N., Wilson, R., Feng, Y., Movsesyan, A., Karlsson Hedestam, G.B., Ward, A.B., and Wyatt, R.T. (2017). Targeted N-glycan deletion at the receptor-binding site retains HIV Env NFL trimer integrity and accelerates the elicited antibody response. PLoS Pathog 13, e1006614.

16. Dubrovskaya, V., Tran, K., Ozorowski, G., Guenaga, J., Wilson, R., Bale, S., Cottrell, C.A., Turner, H.L., Seabright, G., O’Dell, S., et al. (2019). Vaccination with Glycan-Modified HIV NFL Envelope Trimer-Liposomes Elicits Broadly Neutralizing Antibodies to Multiple Sites of Vulnerability. Immunity 51, 915–929 e917.

17. Escolano, A., Steichen, J.M., Dosenovic, P., Kulp, D.W., Golijanin, J., Sok, D., Freund, N.T., Gitlin, A.D., Oliveira, T., Araki, T., et al. (2016). Sequential Immunization Elicits Broadly Neutralizing Anti-HIV-1 Antibodies in Ig Knockin Mice. Cell 166, 1445–1458 e1412.

18. Excler, J.L., Ake, J., Robb, M.L., Kim, J.H., and Plotkin, S.A. (2014). Nonneutralizing functional antibodies: a new “old” paradigm for HIV vaccines. Clin Vaccine Immunol 21, 1023–1036.

19. Geretti, A.M., Harrison, L., Green, H., Sabin, C., Hill, T., Fearnhill, E., Pillay, D., Dunn, D., and Resistance, U.K.C.G.o.H.D. (2009). Effect of HIV-1 subtype on virologic and immunologic response to starting highly active antiretroviral therapy. Clin Infect Dis 48, 1296–1305.

20. Gorny, M.K., Pan, R., Williams, C., Wang, X.H., Volsky, B., O’Neal, T., Spurrier, B., Sampson, J.M., Li, L., Seaman, M.S., et al. (2012). Functional and immunochemical cross-reactivity of V2-specific monoclonal antibodies from HIV-1-infected individuals. Virology 427, 198–207.

21. Guenaga, J., de Val, N., Tran, K., Feng, Y., Satchwell, K., Ward, A.B., and Wyatt, R.T. (2015a). Well-ordered trimeric HIV-1 subtype B and C soluble spike mimetics generated by negative selection display native-like properties. PLoS Pathog 11, e1004570.

22. Guenaga, J., Dubrovskaya, V., de Val, N., Sharma, S.K., Carrette, B., Ward, A.B., and Wyatt, R.T. (2015b). Structure-Guided Redesign Increases the Propensity of HIV Env To Generate Highly Stable Soluble Trimers. J Virol 90, 2806–2817.

23. Guenaga, J., Garces, F., de Val, N., Stanfield, R.L., Dubrovskaya, V., Higgins, B., Carrette, B., Ward, A.B., Wilson, I.A., and Wyatt, R.T. (2017). Glycine Substitution at Helix-to-Coil Transitions Facilitates the Structural Determination of a Stabilized Subtype C HIV Envelope Glycoprotein. Immunity 46, 792–803 e793.

24. Guttman, M., Garcia, N.K., Cupo, A., Matsui, T., Julien, J.P., Sanders, R.W., Wilson, I.A., Moore, J.P., and Lee, K.K. (2014). CD4-induced activation in a soluble HIV-1 Env trimer. Structure 22, 974–984.

25. Guzzo, C., Ichikawa, D., Park, C., Phillips, D., Liu, Q., Zhang, P., Kwon, A., Miao, H., Lu, J., Rehm, C., et al. (2017). Virion incorporation of integrin alpha4beta7 facilitates HIV-1 infection and intestinal homing. Sci Immunol 2.

26. Guzzo, C., Zhang, P., Liu, Q., Kwon, A.L., Uddin, F., Wells, A.I., Schmeisser, H., Cimbro, R., Huang, J., Doria-Rose, N., et al. (2018). Structural Constraints at the Trimer Apex Stabilize the HIV-1 Envelope in a Closed, Antibody-Protected Conformation. mBio 9.

27. Haynes, B.F., Gilbert, P.B., McElrath, M.J., Zolla-Pazner, S., Tomaras, G.D., Alam, S.M., Evans, D.T., Montefiori, D.C., Karnasuta, C., Sutthent, R., et al. (2012). Immune-correlates analysis of an HIV-1 vaccine efficacy trial. N Engl J Med 366, 1275–1286.

28. He, L., Kumar, S., Allen, J.D., Huang, D., Lin, X., Mann, C.J., Saye-Francisco, K.L., Copps, J., Sarkar, A., Blizard, G.S., et al. (2018). HIV-1 vaccine design through minimizing envelope metastability. Sci Adv 4, eaau6769.

29. Hraber, P., Seaman, M.S., Bailer, R.T., Mascola, J.R., Montefiori, D.C., and Korber, B.T. (2014). Prevalence of broadly neutralizing antibody responses during chronic HIV-1 infection. AIDS 28, 163–169.

30. Huang, D., Tran, J.T., Olson, A., Vollbrecht, T., Tenuta, M., Guryleva, M.V., Fuller, R.P., Schiffner, T., Abadejos, J.R., Couvrette, L., et al. (2020). Vaccine elicitation of HIV broadly neutralizing antibodies from engineered B cells. Nat Commun 11, 5850.

31. Jain, P.C., and Varadarajan, R. (2014). A rapid, efficient, and economical inverse polymerase chain reaction-based method for generating a site saturation mutant library. Anal Biochem 449, 90–98.

32. Jones, A.T., Shen, X., Walter, K.L., LaBranche, C.C., Wyatt, L.S., Tomaras, G.D., Montefiori, D.C., Moss, B., Barouch, D.H., Clements, J.D., et al. (2019). HIV-1 vaccination by needle-free oral injection induces strong mucosal immunity and protects against SHIV challenge. Nat Commun 10, 798.

33. Joyce, M.G., Georgiev, I.S., Yang, Y., Druz, A., Geng, H., Chuang, G.Y., Kwon, Y.D., Pancera, M., Rawi, R., Sastry, M., et al. (2017). Soluble Prefusion Closed DS-SOSIP.664-Env Trimers of Diverse HIV-1 Strains. Cell Rep 21, 2992-3002.

34. Kannanganat, S., Wyatt, L.S., Gangadhara, S., Chamcha, V., Chea, L.S., Kozlowski, P.A., LaBranche, C.C., Chennareddi, L., Lawson, B., Reddy, P.B., et al. (2016). High Doses of GM-CSF Inhibit Antibody Responses in Rectal Secretions and Diminish Modified Vaccinia Ankara/Simian Immunodeficiency Virus Vaccine Protection in TRIM5alpha-Restrictive Macaques. J Immunol 197, 3586–3596.

35. Kasturi, S.P., Kozlowski, P.A., Nakaya, H.I., Burger, M.C., Russo, P., Pham, M., Kovalenkov, Y., Silveira, E.L., Havenar-Daughton, C., Burton, S.L., et al. (2017). Adjuvanting a Simian Immunodeficiency Virus Vaccine with Toll-Like Receptor Ligands Encapsulated in Nanoparticles Induces Persistent Antibody Responses and Enhanced Protection in TRIM5alpha Restrictive Macaques. J Virol 91.

36. Klasse, P.J., Ketas, T.J., Cottrell, C.A., Ozorowski, G., Debnath, G., Camara, D., Francomano, E., Pugach, P., Ringe, R.P., LaBranche, C.C., et al. (2018). Epitopes for neutralizing antibodies induced by HIV-1 envelope glycoprotein BG505 SOSIP trimers in rabbits and macaques. PLoS Pathog 14, e1006913.

37. Klasse, P.J., LaBranche, C.C., Ketas, T.J., Ozorowski, G., Cupo, A., Pugach, P., Ringe, R.P., Golabek, M., van Gils, M.J., Guttman, M., et al. (2016). Sequential and Simultaneous Immunization of Rabbits with HIV-1 Envelope Glycoprotein SOSIP.664 Trimers from Clades A, B and C. PLoS Pathog 12, e1005864.

38. Kong, L., He, L., de Val, N., Vora, N., Morris, C.D., Azadnia, P., Sok, D., Zhou, B., Burton, D.R., Ward, A.B., et al. (2016). Uncleaved prefusion-optimized gp140 trimers derived from analysis of HIV-1 envelope metastability. Nat Commun 7, 12040.

39. Kulp, D.W., Steichen, J.M., Pauthner, M., Hu, X., Schiffner, T., Liguori, A., Cottrell, C.A., Havenar-Daughton, C., Ozorowski, G., Georgeson, E., et al. (2017). Structure-based design of native-like HIV-1 envelope trimers to silence non-neutralizing epitopes and eliminate CD4 binding. Nat Commun 8, 1655.

40. Kwon, Y.D., Pancera, M., Acharya, P., Georgiev, I.S., Crooks, E.T., Gorman, J., Joyce, M.G., Guttman, M., Ma, X., Narpala, S., et al. (2015). Crystal structure, conformational fixation and entry-related interactions of mature ligand-free HIV-1 Env. Nat Struct Mol Biol 22, 522–531.

41. Landais, E., Huang, X., Havenar-Daughton, C., Murrell, B., Price, M.A., Wickramasinghe, L., Ramos, A., Bian, C.B., Simek, M., Allen, S., et al. (2016). Broadly Neutralizing Antibody Responses in a Large Longitudinal Sub-Saharan HIV Primary Infection Cohort. PLoS Pathog 12, e1005369.

42. Lander, G.C., Stagg, S.M., Voss, N.R., Cheng, A., Fellmann, D., Pulokas, J., Yoshioka, C., Irving, C., Mulder, A., Lau, P.W., et al. (2009). Appion: an integrated, database-driven pipeline to facilitate EM image processing. J Struct Biol 166, 95–102.

43. Lee, J.H., Andrabi, R., Su, C.Y., Yasmeen, A., Julien, J.P., Kong, L., Wu, N.C., McBride, R., Sok, D., Pauthner, M., et al. (2017). A Broadly Neutralizing Antibody Targets the Dynamic HIV Envelope Trimer Apex via a Long, Rigidified, and Anionic beta-Hairpin Structure. Immunity 46, 690–702.

44. Li, Y., Svehla, K., Louder, M.K., Wycuff, D., Phogat, S., Tang, M., Migueles, S.A., Wu, X., Phogat, A., Shaw, G.M., et al. (2009). Analysis of neutralization specificities in polyclonal sera derived from human immunodeficiency virus type 1-infected individuals. J Virol 83, 1045–1059.

45. Liao, H.X., Bonsignori, M., Alam, S.M., McLellan, J.S., Tomaras, G.D., Moody, M.A., Kozink, D.M., Hwang, K.K., Chen, X., Tsao, C.Y., et al. (2013). Vaccine induction of antibodies against a structurally heterogeneous site of immune pressure within HIV-1 envelope protein variable regions 1 and 2. Immunity 38, 176–186.

46. Liu, Q., Acharya, P., Dolan, M.A., Zhang, P., Guzzo, C., Lu, J., Kwon, A., Gururani, D., Miao, H., Bylund, T., et al. (2017). Quaternary contact in the initial interaction of CD4 with the HIV-1 envelope trimer. Nat Struct Mol Biol 24, 370–378.

47. Lynch, R.M., Wong, P., Tran, L., O’Dell, S., Nason, M.C., Li, Y., Wu, X., and Mascola, J.R. (2015). HIV-1 fitness cost associated with escape from the VRC01 class of CD4 binding site neutralizing antibodies. J Virol 89, 4201–4213.

48. McGuire, A.T. (2019). Targeting broadly neutralizing antibody precursors: a naive approach to vaccine design. Curr Opin HIV AIDS 14, 294–301.

49. Montefiori, D.C. (2009). Measuring HIV neutralization in a luciferase reporter gene assay. Methods Mol Biol 485, 395–405.

50. Ogura, T., Iwasaki, K., and Sato, C. (2003). Topology representing network enables highly accurate classification of protein images taken by cryo electron-microscope without masking. J Struct Biol 143, 185–200.

51. Pancera, M., Shahzad-Ul-Hussan, S., Doria-Rose, N.A., McLellan, J.S., Bailer, R.T., Dai, K., Loesgen, S., Louder, M.K., Staupe, R.P., Yang, Y., et al. (2013). Structural basis for diverse N-glycan recognition by HIV-1-neutralizing V1-V2-directed antibody PG16. Nat Struct Mol Biol 20, 804–813.

52. Pauthner, M., Havenar-Daughton, C., Sok, D., Nkolola, J.P., Bastidas, R., Boopathy, A.V., Carnathan, D.G., Chandrashekar, A., Cirelli, K.M., Cottrell, C.A., et al. (2017). Elicitation of Robust Tier 2 Neutralizing Antibody Responses in Nonhuman Primates by HIV Envelope Trimer Immunization Using Optimized Approaches. Immunity 46, 1073–1088 e1076.

53. Pejchal, R., Walker, L.M., Stanfield, R.L., Phogat, S.K., Koff, W.C., Poignard, P., Burton, D.R., and Wilson, I.A. (2010). Structure and function of broadly reactive antibody PG16 reveal an H3 subdomain that mediates potent neutralization of HIV-1. Proc Natl Acad Sci U S A 107, 11483–11488.

54. Perez, L.G., Martinez, D.R., deCamp, A.C., Pinter, A., Berman, P.W., Francis, D., Sinangil, F., Lee, C., Greene, K., Gao, H., et al. (2017). V1V2-specific complement activating serum IgG as a correlate of reduced HIV-1 infection risk in RV144. PLoS One 12, e0180720.

55. Pettersen, E.F., Goddard, T.D., Huang, C.C., Couch, G.S., Greenblatt, D.M., Meng, E.C., and Ferrin, T.E. (2004). UCSF Chimera--a visualization system for exploratory research and analysis. J Comput Chem 25, 1605–1612.

56. Plotnik, D., Guo, W., Cleveland, B., von Haller, P., Eng, J.K., Guttman, M., Lee, K.K., Arthos, J., and Hu, S.L. (2017). Extracellular Matrix Proteins Mediate HIV-1 gp120 Interactions with alpha4beta7. J Virol 91.

57. Rademeyer, C., Korber, B., Seaman, M.S., Giorgi, E.E., Thebus, R., Robles, A., Sheward, D.J., Wagh, K., Garrity, J., Carey, B.R., et al. (2016). Features of Recently Transmitted HIV-1 Clade C Viruses that Impact Antibody Recognition: Implications for Active and Passive Immunization. PLoS Pathog 12, e1005742.

58. Rerks-Ngarm, S., Pitisuttithum, P., Nitayaphan, S., Kaewkungwal, J., Chiu, J., Paris, R., Premsri, N., Namwat, C., de Souza, M., Adams, E., et al. (2009). Vaccination with ALVAC and AIDSVAX to prevent HIV-1 infection in Thailand. N Engl J Med 361, 2209–2220.

59. Robb, M.L., Rerks-Ngarm, S., Nitayaphan, S., Pitisuttithum, P., Kaewkungwal, J., Kunasol, P., Khamboonruang, C., Thongcharoen, P., Morgan, P., Benenson, M., et al. (2012). Risk behaviour and time as covariates for efficacy of the HIV vaccine regimen ALVAC-HIV (vCP1521) and AIDSVAX B/E: a post-hoc analysis of the Thai phase 3 efficacy trial RV 144. Lancet Infect Dis 12, 531–537.

60. Roederer, M., Keele, B.F., Schmidt, S.D., Mason, R.D., Welles, H.C., Fischer, W., Labranche, C., Foulds, K.E., Louder, M.K., Yang, Z.Y., et al. (2014). Immunological and virological mechanisms of vaccine-mediated protection against SIV and HIV. Nature 505, 502–508.

61. Sanders, R.W., Derking, R., Cupo, A., Julien, J.P., Yasmeen, A., de Val, N., Kim, H.J., Blattner, C., de la Pena, A.T., Korzun, J., et al. (2013). A next-generation cleaved, soluble HIV-1 Env trimer, BG505 SOSIP.664 gp140, expresses multiple epitopes for broadly neutralizing but not non-neutralizing antibodies. PLoS Pathog 9, e1003618.

62. Sanders, R.W., and Moore, J.P. (2017). Native-like Env trimers as a platform for HIV-1 vaccine design. Immunol Rev 275, 161–182.

63. Sanders, R.W., van Gils, M.J., Derking, R., Sok, D., Ketas, T.J., Burger, J.A., Ozorowski, G., Cupo, A., Simonich, C., Goo, L., et al. (2015). HIV-1 VACCINES. HIV-1 neutralizing antibodies induced by native-like envelope trimers. Science 349, aac4223.

64. Sharma, S.K., de Val, N., Bale, S., Guenaga, J., Tran, K., Feng, Y., Dubrovskaya, V., Ward, A.B., and Wyatt, R.T. (2015). Cleavage-independent HIV-1 Env trimers engineered as soluble native spike mimetics for vaccine design. Cell Rep 11, 539–550.

65. Sliepen, K., Ozorowski, G., Burger, J.A., van Montfort, T., Stunnenberg, M., LaBranche, C., Montefiori, D.C., Moore, J.P., Ward, A.B., and Sanders, R.W. (2015). Presenting native-like HIV-1 envelope trimers on ferritin nanoparticles improves their immunogenicity. Retrovirology 12, 82.

66. Styles, T.M., Gangadhara, S., Reddy, P.B.J., Hicks, S., LaBranche, C.C., Montefiori, D.C., Derdeyn, C.A., Kozlowski, P.A., Velu, V., and Amara, R.R. (2019). Human Immunodeficiency Virus C.1086 Envelope gp140 Protein Boosts following DNA/Modified Vaccinia Virus Ankara Vaccination Fail To Enhance Heterologous Anti-V1V2 Antibody Response and Protection against Clade C Simian-Human Immunodeficiency Virus Challenge. J Virol 93.

67. Suloway, C., Pulokas, J., Fellmann, D., Cheng, A., Guerra, F., Quispe, J., Stagg, S., Potter, C.S., and Carragher, B. (2005). Automated molecular microscopy: the new Leginon system. J Struct Biol 151, 41–60.

68. Tassaneetrithep, B., Tivon, D., Swetnam, J., Karasavvas, N., Michael, N.L., Kim, J.H., Marovich, M., and Cardozo, T. (2014). Cryptic determinant of alpha4beta7 binding in the V2 loop of HIV-1 gp120. PLoS One 9, e108446.

69. Tomaras, G.D., Yates, N.L., Liu, P., Qin, L., Fouda, G.G., Chavez, L.L., Decamp, A.C., Parks, R.J., Ashley, V.C., Lucas, J.T., et al. (2008). Initial B-cell responses to transmitted human immunodeficiency virus type 1: virion-binding immunoglobulin M (IgM) and IgG antibodies followed by plasma anti-gp41 antibodies with ineffective control of initial viremia. J Virol 82, 12449–12463.

70. Torrents de la Pena, A., Julien, J.P., de Taeye, S.W., Garces, F., Guttman, M., Ozorowski, G., Pritchard, L.K., Behrens, A.J., Go, E.P., Burger, J.A., et al. (2017). Improving the Immunogenicity of Native-like HIV-1 Envelope Trimers by Hyperstabilization. Cell Rep 20, 1805–1817.

71. van Eeden, C., Wibmer, C.K., Scheepers, C., Richardson, S.I., Nonyane, M., Lambson, B., Mkhize, N.N., Vijayakumar, B., Sheng, Z., Stanfield-Oakley, S., et al. (2018). V2-Directed Vaccine-like Antibodies from HIV-1 Infection Identify an Additional K169-Binding Light Chain Motif with Broad ADCC Activity. Cell Rep 25, 3123–3135 e3126.

72. Verkerke, H.P., Williams, J.A., Guttman, M., Simonich, C.A., Liang, Y., Filipavicius, M., Hu, S.L., Overbaugh, J., and Lee, K.K. (2016). Epitope-Independent Purification of Native-Like Envelope Trimers from Diverse HIV-1 Isolates. J Virol 90, 9471–9482.

73. Voss, J.E., Andrabi, R., McCoy, L.E., de Val, N., Fuller, R.P., Messmer, T., Su, C.Y., Sok, D., Khan, S.N., Garces, F., et al. (2017). Elicitation of Neutralizing Antibodies Targeting the V2 Apex of the HIV Envelope Trimer in a Wild-Type Animal Model. Cell Rep 21, 222–235.

74. Weis, D.D., Wales, T.E., Engen, J.R., Hotchko, M., and Ten Eyck, L.F. (2006). Identification and characterization of EX1 kinetics in H/D exchange mass spectrometry by peak width analysis. J Am Soc Mass Spectrom 17, 1498–1509.

75. Xu, K., Acharya, P., Kong, R., Cheng, C., Chuang, G.Y., Liu, K., Louder, M.K., O’Dell, S., Rawi, R., Sastry, M., et al. (2018). Epitope-based vaccine design yields fusion peptide-directed antibodies that neutralize diverse strains of HIV-1. Nat Med 24, 857–867.

76. Yates, N.L., Liao, H.X., Fong, Y., deCamp, A., Vandergrift, N.A., Williams, W.T., Alam, S.M., Ferrari, G., Yang, Z.Y., Seaton, K.E., et al. (2014). Vaccine-induced Env V1-V2 IgG3 correlates with lower HIV-1 infection risk and declines soon after vaccination. Sci Transl Med 6, 228ra239.

77. Zolla-Pazner, S., Alvarez, R., Kong, X.P., and Weiss, S. (2019). Vaccine-induced V1V2-specific antibodies control and or protect against infection with HIV, SIV and SHIV. Curr Opin HIV AIDS 14, 309–317.

78. Zolla-Pazner, S., deCamp, A., Gilbert, P.B., Williams, C., Yates, N.L., Williams, W.T., Howington, R., Fong, Y., Morris, D.E., Soderberg, K.A., et al. (2014). Vaccine-induced IgG antibodies to V1V2 regions of multiple HIV-1 subtypes correlate with decreased risk of HIV-1 infection. PLoS One 9, e87572.

79. Zolla-Pazner, S., deCamp, A.C., Cardozo, T., Karasavvas, N., Gottardo, R., Williams, C., Morris, D.E., Tomaras, G., Rao, M., Billings, E., et al. (2013). Analysis of V2 antibody responses induced in vaccinees in the ALVAC/AIDSVAX HIV-1 vaccine efficacy trial. PLoS One 8, e53629.

## References

80. Kwon, Y.D., Pancera, M., Acharya, P., Georgiev, I.S., Crooks, E.T., Gorman, J., Joyce, M.G., Guttman, M., Ma, X., Narpala, S., et al. (2015). Crystal structure, conformational fixation and entry-related interactions of mature ligand-free HIV-1 Env. Nat Struct Mol Biol 22, 522–531.

81. Montefiori, D.C. (2009). Measuring HIV neutralization in a luciferase reporter gene assay. Methods Mol Biol 485, 395–405.

82. Pettersen, E.F., Goddard, T.D., Huang, C.C., Couch, G.S., Greenblatt, D.M., Meng, E.C., and Ferrin, T.E. (2004). UCSF Chimera--a visualization system for exploratory research and analysis. J Comput Chem 25, 1605–1612.

83. Zolla-Pazner, S., deCamp, A., Gilbert, P.B., Williams, C., Yates, N.L., Williams, W.T., Howington, R., Fong, Y., Morris, D.E., Soderberg, K.A., et al. (2014). Vaccine-induced IgG antibodies to V1V2 regions of multiple HIV-1 subtypes correlate with decreased risk of HIV-1 infection. PLoS One 9, e87572.

